# Human Body Single-Cell Atlas of 3D Genome Organization and DNA Methylation

**DOI:** 10.1101/2025.03.23.644697

**Authors:** Jingtian Zhou, Yue Wu, Hanqing Liu, Wei Tian, Rosa G Castanon, Anna Bartlett, Zuolong Zhang, Guocong Yao, Dengxiaoyu Shi, Ben Clock, Samantha Marcotte, Joseph R. Nery, Michelle Liem, Naomi Claffey, Lara Boggeman, Cesar Barragan, Rafael Arrojo e Drigo, Annika K. Weimer, Minyi Shi, Johnathan Cooper-Knock, Sai Zhang, Michael P. Snyder, Sebastian Preissl, Bing Ren, Carolyn O’Connor, Shengbo Chen, Chongyuan Luo, Jesse R. Dixon, Joseph R. Ecker

**Author notes:** Corresponding authors. (J.Z.); (J.R.D.); (J.R.E.). Co-first authors (Jingtian Zhou, Yue Wu).

## Abstract

Higher-order chromatin structure and DNA methylation are critical for gene regulation, but how these vary across the human body remains unclear. We performed multi-omic profiling of 3D genome structure and DNA methylation for 86,689 single nuclei across 16 human tissues, identifying 35 major and 206 cell subtypes. We revealed extensive changes in CG and non-CG methylation across almost all cell types and characterized 3D chromatin structure at an unprecedented cellular resolution. Intriguingly, extensive discrepancies exist between cell types delineated by DNA methylation and genome structure, indicating that the role of distinct epigenomic features in maintaining cell identity may vary by lineage. This study expands our understanding of the diversity of DNA methylation and chromatin structure and offers an extensive reference for exploring gene regulation in human health and disease.

## Main Text

Distinct cellular identities are established by cooperative gene regulatory programs that lead to cell type-specific gene expression patterns. Proper patterns of lineage-specific gene expression require coordination between distinct chromatin regulatory processes, including open chromatin, transcription factor binding, histone modifications, DNA methylation, and higher-order 3D chromatin structure (*1*). The development of single-cell technologies to profile gene expression (*2*) or aspects of the chromatin regulatory landscape (*3–7*) has led to efforts to comprehensively profile these features across human cell types to understand the diversity of gene expression regulation across individual human cells (*8–11*). These efforts have led to the identification of novel human cell states (*12*) and the global characterization of regulatory elements that contribute to human disease risk (*13–17*). This has been most notably accomplished using single-cell methods to profile gene expression or open chromatin data. In contrast, single-cell methods that analyze other critical mechanisms of gene regulation, such as DNA methylation (*6*, *18*, *19*) or 3D genome architecture (*20–23*), have only been applied in limited tissues and cell types.

Previous efforts to broadly characterize these epigenomic features across human cell types have focused on using bulk tissues to profile DNA methylation (*24*, *25*) or 3D genome organization (*26*). Similarly, these methods have been applied to diverse sets of cultured primary or cancer cell lines (*27–31*). These studies have revealed distinct differences in DNA methylation or 3D genome architecture, for example, identifying the presence of non-CG methylation in human brain samples and identifying the presence of partially methylated domains in cancer cell lines and some human tissues. More recently, DNA methylation patterns have been profiled across a broad spectrum of human cell types using cell sorting to isolate specific populations (*32*). However, cell sorting has limitations, such as an inability to isolate subpopulations of cells, such as neuronal subtypes, without the use of genetically encoded markers, which is impossible in human tissue samples. Similarly, cell sorting and related techniques have enabled insights into cell type-specific patterns of 3D chromatin compartments, domains, and chromatin loops between cultured cells or tissues (*26–28*, *33*). However, these methods have not been comprehensively applied to distinguish human cell types from *in vivo* tissue samples, limiting our understanding of how distinct DNA methylation states or aspects of 3D chromatin folding differ among human cell types. Further, most previous efforts have studied these features in isolation, so our understanding of the relationship between these distinct chromatin regulatory mechanisms in single cells remains unexplored.

To better understand the diversity of DNA methylation and 3D genome structure and how these contribute to cell type-specific patterns of gene expression, we have applied the single-nucleus methyl-3C (*22*) method to generate a “body map” of DNA methylation and 3D genome architecture for single cells derived from 16 human tissues. Using this data, we have identified 35 major cell types and 206 cellular subtypes across human tissues. This has allowed us to characterize DNA methylation patterns, including the widespread presence of low-level non-CG methylation across human cell types, and to identify partially methylated-like domains in a wide variety of cell types. Further, we have used this to identify chromatin compartments, domains, and loops across human cells. Interestingly, comparing 3D chromatin structure and DNA methylation, we find instances where the two modalities show differing clustering patterns between single cells, suggesting that these two genomic features show distinct dynamics during cellular differentiation. Taken together, our study represents the first single-cell body map atlas of human cells’ DNA methylation or 3D genome architecture. This dataset will provide insights into the variability of these features across a large number of distinct human cell types and will be a valuable resource for understanding the role these gene regulatory mechanisms play in establishing cellular identity.

### A single-cell atlas of DNA methylation and 3D genome structure across human tissues

To better understand the diversity of DNA methylation and 3D chromatin conformation across human cell types, we used the single nucleus methyl-3C-seq (snm3C-seq) assay to profile cells from 16 human tissues (Fig. 1A). Snm3C-seq generates information on DNA methylome and 3D genome structure simultaneously within single cells (Fig. 1, B and C). After quality control and doublet removal (materials and methods), we obtained 86,689 cells with 195 billion non-clonal methylation reads and 18 billion long-range chromatin contacts in total. For each tissue, we profiled cells from at least two human donors (table S1). For 12/16 tissues, the samples were previously profiled as part of the ENTEx collaborative project (*34*), such that there are rich public datasets for these tissues, including single-nucleus assay for transcriptome accessible chromatin sequencing (snATAC-seq) from the same tissue samples (*14*). The remaining four tissue types were chosen to broaden the diversity of cell types and lineages represented in the atlas, including brain (primary motor cortex), placenta, isolated pancreatic islets, and peripheral blood. Together, this data represents the first single-cell atlas of DNA methylation or 3D genome architecture across human tissues.

**Fig. 1.**
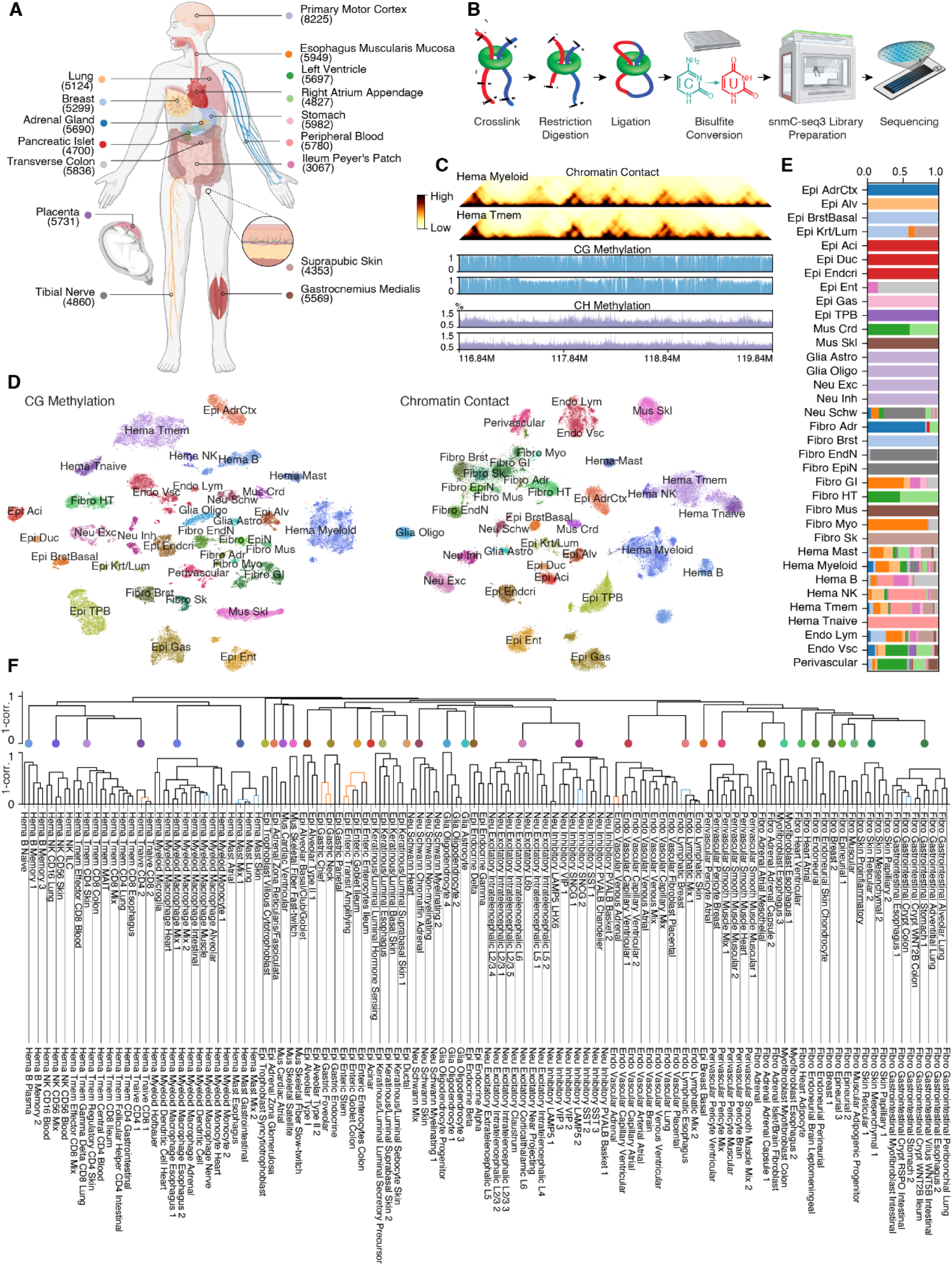
Single-cell atlas of human DNA methylation and 3D genome structure. (**A**) Schematic showing 16 human tissues profiled by the snm3C-seq assay. Below each tissue are the number of single cells passing quality control filters. Each tissue is assigned a specific color that is used throughout the manuscript. (**B**) Workflow diagram of the snm3C-seq assay. (**C**) Browser shot of DNA methylation and chromatin contacts in Myeloid (Hema Myeloid) or T-memory cells (Hema Tmem) after clustering and merging single cells into pseudobulk cell types. (**D**) t-distributed Stochastic Neighbor Embedding (t-SNE) of all single cells (n=86,689) using either CpG cytosine methylation (mCG, left) or chromatin contact (right) data colored by major types. The same color palette is used for major types throughout the manuscript. (**E**) The proportion of cells derived from each tissue for the 35 major cell types. The colors for each tissue are labeled in panel A. (**F**) Dendrogram of major cell types (top) and subtypes (bottom). The x-axis coordinate of each major type is manually aligned to the center of the highest branch in the subtype dendrogram of the major type.

To analyze the large dataset across donors, tissues, and modalities, several analytical challenges were addressed with novel or improved computational methods. We generated an exclusion list for 3D genome contacts to remove the artifactual contacts introduced by misalignments over repeat regions due to the 3-base pair genome alignment (fig. S1). We introduced methods to cluster single cells with DNA methylation across 5kb genomic bins (fig. S2, A and B), which in many tissues outperformed our 100kb-bin-based clustering method to cluster brain cells. We used an anchor-based method for the integration of data between donors and geometric distance-based filtering to refine the anchors (fig. S2, C and D). We used the combined information of DNA methylation and 3D genome structure to classify cells into 35 major types (Fig. 1D and fig. S2E). Major cell types were a mixture of cells that were exclusive to single tissues (for example, excitatory neurons from primary motor cortex) or that were shared across different tissues (for example, memory T-cells) (Fig. 1E). We further sub-classified major type cells into 206 cell subtypes (Fig. 1F). This was accomplished by both jointly and separately clustering cells by DNA methylation and chromatin contact data and further integrating with scRNA-seq and snATAC-seq data (figs. S3 to 5, see materials and methods for details). Finally, to facilitate the exploration of the data, we have developed an interactive cell browser (https://humancellepigenomeatlas.arcinstitute.org) to display both the DNA methylation and chromatin contact data across tissues, major types and subtypes.

We compared mCG and chromatin contacts across cell types and donors to validate the quality of our major and subtype clustering. We observed the strongest correlation between the identical cell type across different donors for both CG methylation (mCG) and chromatin contacts (fig. S6). In contrast, we observed a notably lower correlation in mCG or chromatin contacts between different cell subtypes or different major types, confirming the quality of the clustering and cell assignment. We also compared the DNA methylation profiles generated in this study with previous studies using sorted primary cells (*32*) or bulk human tissues (*24*). We observed strong correlations in DNA methylation profiles between the cells classified from our snm3C-seq data with previous methylomes from the corresponding sorted cell populations and bulk tissues (fig. S7). The clearest discrepancies between the DNA methylation patterns in this dataset versus the prior sorted or bulk tissues are related to tissue sources. For example, the previous study of sorted methylomes included tissues (i.e. prostate) not profiled here. Likewise, some sorted populations (i.e., “neurons,” fig. S7A) contain heterogeneous subtypes of cells that we are able to distinguish using our single-cell-based methods that are not readily discerned using cell surface markers. Our dataset represents a valuable resource for the community to study cell-type-resolved patterns of DNA methylation and 3D chromatin conformation across diverse tissues and cell types.

### Variability of CG methylation across cell types

Cytosine DNA methylation occurs in a CG dinucleotide context across diverse cell types. Next, we analyzed patterns of mCG across major cell types and observed a wide variety of average mCG frequencies within each major cell type (Fig. 2A). The cells with the lowest frequency of mCG were epithelial trophoblast cells (Epi TPB), with an average of 59% methylation. In contrast, the highest frequency of mCG was observed in inhibitory neurons (Neu Inh), with an average of 81% methylation. Examining the distribution of methylation of individual cytosines, we observed that most cell types show a bimodal distribution of mCG, where cytosines are either largely unmethylated (<10%) or fully methylated (>70%), with relatively few major cell types showing evidence of a high frequency of partially methylated CpGs throughout the genome (Fig. 2B). A subset of major cell types, however, were enriched for partially methylated cytosines, including trophoblast epithelial cells (Epi TPB), memory B-cells (Hema Bmem), and acinar epithelial cells (Epi Aci) (Fig. 2B). The partially methylated CpGs in these lineages are found in large discrete domains (Fig. 2C), previously termed Partially Methylated Domains (PMDs) (*31*). PMDs have been described as a prominent feature in cancer cell lines and cultured fibroblasts (*35*), and previous studies have also found PMDs in bulk DNA methylation profiling from trophoblast tissue (*36*).

**Fig. 2.**
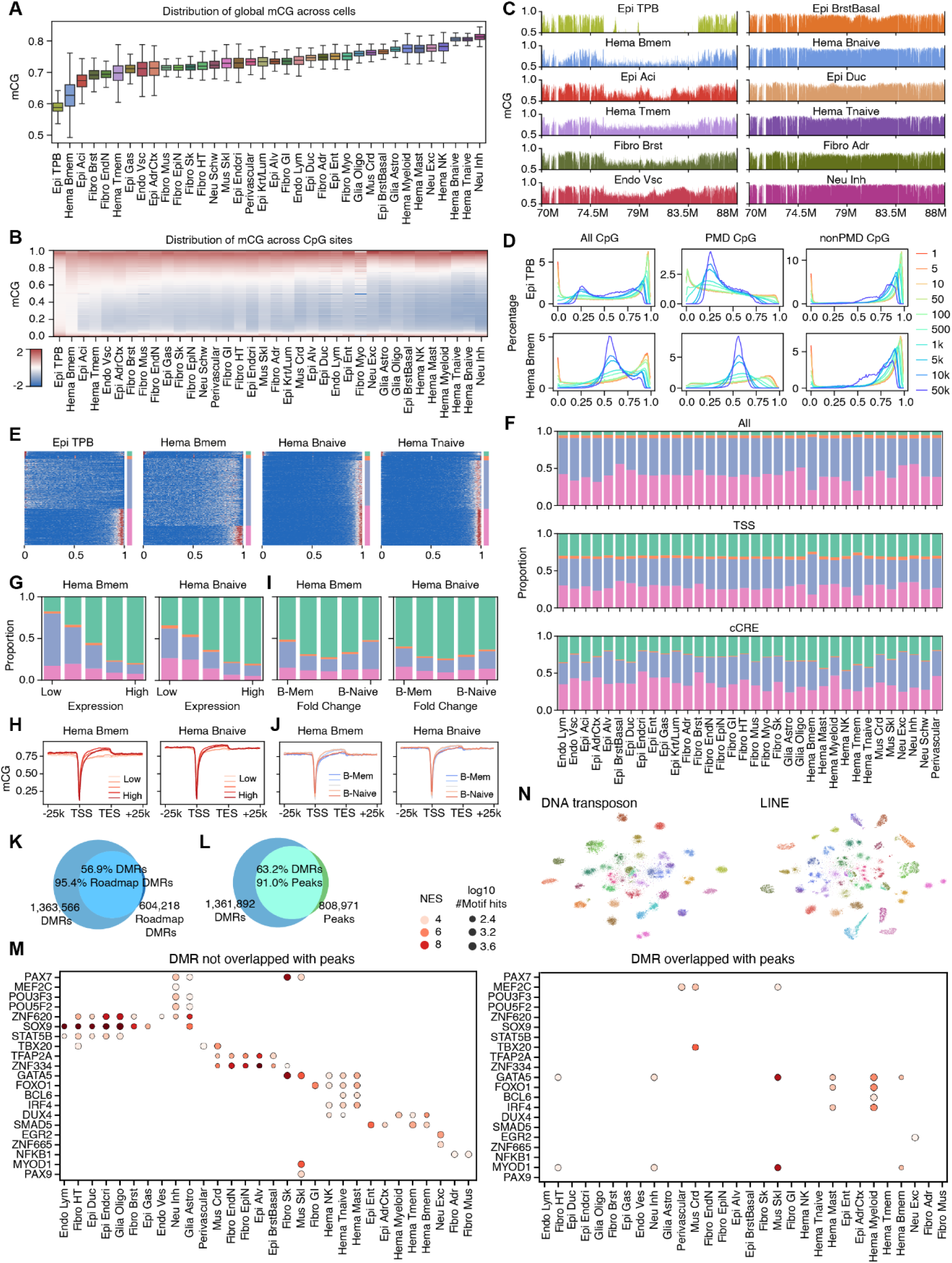
CpG methylation across cell types. (**A**) Boxplot showing the distribution of average genome wide mCG of single cells grouped by major types. For all box plots throughout the manuscript, the center line denotes the median; box limits denote first and third quartiles; and whiskers denote 1.5 × the interquartile range. (**B**) The distribution of mCG of individual CpGs in each major type. (**C**) Genome browser views of mCG at chr2:70,000,000-88,000,000 showing the presence of partially methylated domains (PMDs). (**D**) Distribution of mCG in trophoblast epithelial cells (Epi TPB, top) or memory B-cells (Hema Bmem, bottom) throughout the genome using varying bin sizes over all CpGs (left), CpGs within regions called as PMDs (middle), and CpGs in non-PMD regions (right) based on assigning regions into DNA methylation compartments. (**E**) Distribution of mCG over CpGs for 10kb bins (row) ordered by methylation compartments. Color bars from top to bottom: fully unmethylated, bimodal, partially methylated, and fully methylated. The same color palette is used for methylation compartments throughout the manuscript. (**F**) Fraction of the genome associated with each methylation compartment for all 10kb bins (top), transcription start sites (TSS - middle), or candidate cis-regulatory elements (cCREs - bottom). (**G-J**) Fraction of genes associated with each methylation compartment (G and I) or gene body mCG (H and J) in memory (Hema Bmem) or naive (Hema Bnaive) B cells stratified by gene expression level (G and H) or fold-change between Naive and Memory B cells (I and J). (**K**) Overlap between all DMRs from this study and DMRs from Schultz et al. 2015. (**L**) Overlap between DMRs identified in this study and previous scATAC-seq peaks from matching cell types. Of note, this DMR set excludes trophoblast and Naive B cells because of a lack of matching ATAC-seq. (**M**) Enrichment of selected transcription factor (TF) binding motifs in major cell types for non-peak DMRs (left) and peak DMRs (right). Normalized enrichment scores (NES) are row-wise Z-scored. (**N**) t-SNE of downsampled cells (n=26,423) using mCG at DNA transposon (left) and long terminal repeats (LTR, right) colored by major types.

To characterize the presence of PMDs across human cell types, we first applied published tools to identify PMDs from the pseudobulk mCG in major cell types (*37*). However, the ability to accurately call PMDs appeared to vary by lineage. For example, in lineages with visually apparent PMDs, DNMTools(*38*) missed a large number of partially methylated regions in memory B cells (Hema Bmem), while it completely misclassified the hypo-methylated regions as PMDs in memory T cells (Hema Tmem) (fig. S8A). We therefore developed a computational framework to identify PMDs. Given that PMDs are over a kilobase long and enriched for partially methylated CpGs, we clustered 10kb genomic bins into groups according to the distribution of methylation across CG sites within the 10kb bin (fig. S8B). The method accurately identifies PMDs (fig. S8A), with the partially methylated CpGs enriched in PMDs and depleted from non-PMD regions (Fig. 2D).

If we identified PMDs in one lineage with readily apparent PMDs, such as memory B-cells (Hema Bmem), and then examined mCG levels over the same genomic regions in related types without apparent PMDs such as naive B-cells (Hema Bnaive), we observed that these regions show lower methylation levels compared with non-PMD regions even in cell types that do not show obvious signatures of PMDs (fig. S8C). This suggests that PMDs may exist along a continuum across cell types. Further, given that the PMD region shows lower mCG levels even in cells that lack obvious PMDs, this suggests that the genome may be divided into “DNA methylation compartments” based on the local distribution of mCG levels across cell types. To better classify the genome according to the continuity of mCG within DNA methylation compartments, we applied the clustering framework to all the cell types (fig. S8D). This showed that the genome can be divided into four compartments based on the patterns of mCG frequency in each cell type. These consist of two major compartments comprising most of the genome that showed patterns of full cytosine methylation (pink) or partial cytosine methylation (blue), and two minor clusters that show a complete absence of mCG (green) and a bimodal distribution of mCG (orange) (Fig. 2E and fig. S8, B and D). When we stratify regions of the genome by methylation compartments in cells without strong PMDs (Hema Tnaive, Hema Bnaive), we observe that in related lineages (Hema Tmem or Hema Bmem) that have readily apparent PMDs, the partially methylated compartments show greater reduction in their mCG compared to the fully methylated compartments (fig. S8, E and F). This is consistent with the idea that, across cell types, the level of depletion of mCG within the PMD-like compartment exists on a continuum, with some lineages showing only mild depletion of mCG within the PMD-like compartment and others with more pronounced depletion.

Examining the patterns of methylation compartments, we find that the overall distribution of the four compartments is similar across major cell types (fig. S8D). However, some lineages with obvious PMDs also show an expansion of the partially methylated compartment (Fig. 2F). The partially and fully methylated compartment occupies 34-72% and 20-56% of the genome, respectively (90-92% in total; Fig. 2F, top panel). In contrast, the fully unmethylated compartment (green) is strongly enriched for transcription start sites (TSS, Fig. 2F, middle panel) and candidate *cis*-regulatory elements (cCREs, Fig. 2F bottom panel). Consistent with the association with active regulatory elements, we observe that the hypomethylated compartment is strongly associated with higher gene expression levels (Fig. 2G and fig. S9A). Likewise, we observe that higher expressed genes have low levels of promoter mCG and higher gene body mCG compared with lower expressed genes that have higher promoter methylation and lower gene body methylation (Fig. 2H and fig. S9B). Interestingly, in contrast to the observation that the hypomethylated compartment is more likely to contain gene TSSs compared to the genome-wide background, we find that the most variable genes are more likely to reside in the fully or partially methylated compartments compared with genes that do not show expression differences (Fig. 2, I and J, and fig. S9, C to F). This suggests that genes that reside within the partially or fully methylated compartments may be associated with mechanisms of dynamic gene regulation.

To understand patterns of dynamic mCG at a higher genomic resolution, we identified differentially methylated regions (DMRs) between major types and subtypes within each major type. In total, we identified 1,364,566 DMRs across all cell types, with an average length of 377bp, covering 17.9% of the genome in total. 39.3% of the DMRs are differentially methylated between both major types and subtypes. In comparison, 48.2% are differential between subtypes but not major types, suggesting the boosted power of predicting cCREs by a higher cell type resolution. Compared to tissue-level DMRs previously identified from bulk tissue methylome profiling, we identified 95.4% of previous DMRs while identifying 587,648 more DMRs (Fig. 2K). 1,249,253 (91.5%) of DMRs are >2kb from TSS, representing a large repertoire of distal cCREs. DMRs are enriched near TSS but depleted from a core promoter region of ±1kb surrounding TSS (fig. S10A). DMRs are usually hypomethylated in specific cell types while hypermethylated in most other cell types, with only 1% DMRs hypomethylated in >90% subtypes and 96% DMRs hypomethylated in <10% subtypes, while only 10% DMRs hypermethylated in <10% subtypes (fig. S10, B and C).

We also compared DMRs with snATAC-seq data generated from the same tissue samples. Within a given major type, 48% of ATAC-seq peaks overlapped DMRs, while 33% of DMRs overlapped ATAC-seq peaks (fig. S11A). Overall, 91.9% of ATAC-seq peaks overlapped DMRs across all tissues, yet 61.9% of DMRs overlapped ATAC-seq peaks (Fig. 2L). Examining the patterns of DNA methylation across cell types at DMRs, we observed that the DMR methylation levels were highly correlated between similar lineages (fig. S11B). This was similar for DMRs that overlapped ATAC-seq peaks (peak DMR) and those that did not (non-peak DMR; fig. S11B), suggesting that both classes of DMRs contain information regarding the cis-regulatory landscape of cells.

Analysis of transcription factor motifs within cell type-specific DMRs could show evidence of which transcription factors contributed to the establishment of lineage-specific patterns of gene expression. For instance, we identified TBX20, an evolutionarily conserved master regulator for cardiac cell functions (*39*), enriched in cardiac muscle cells, heart fibroblast, and perivascular cells. IRF4, FOXO1, and BCL6 motifs are enriched in the immune cell lineage(*40–42*). Additionally, PAX7, MYOG, and MYOD1, which are important players in myogenesis (*43*), were identified in the skeletal muscle (Fig. 2M). These enrichments were similar for peak and non-peak DMRs (fig. S11C). Overall, 559 motifs were found across major types that were shared between peak and non-peak DMRs, with 514 motifs unique to peak DMRs and 1249 motifs unique to non-peak DMRs. Notably, many TF motifs regulating cell development and functions are only found by non-peak DMRs but not peak DMRs in the corresponding cell type. Such TFs are from various families, including the POU domain family TFs like POU3F3 and POU5F2, which are enriched in the neuronal lineage(*44*), and Zinc-finger family TFs like ZNF620, ZNF334, and ZNF665 that are enriched in epithelial cells, fibroblasts, and neuronal cells (Fig. 2M).

Across all lineages, we observed a depletion of DNA methylation at ATAC-seq peaks for both mCG and mCH, with generally stronger depletion of methylation in the cell types where the ATAC-seq peaks were called (fig. S12, A and B). Although the vast majority of peaks have depleted mCG, we also noticed a small fraction of them are hyper-methylated (fig. S12C). Analyses of the motif enrichment between hyper-methylated peaks against hypo-methylated ones identified many TFs that have been reported as preferring binding methylated motifs (fig. S12, D and E).

Repeat regions have also been suggested to host a considerable amount of regulatory elements and can be important to phenotype-level gene functions. We therefore examined the cell-type specificity of DNA methylation at different categories of repeats across the genome. Interestingly, many types of repeat elements could separate most major cell types with their mCG, including DNA transposons, LINE, and LTR (Fig. 2N and fig. S13). Shorter regions (e.g., SINE, satellite DNA, retroposon) showed lower performance of cell clustering (Fig. 2N), which could result from a weaker specificity or less accurate quantification due to coverage limitation of single-cell data. Since many repeat elements overlap genes and ATAC peaks, we further tested whether we could still distinguish the cell types after removing all the peaks within 2kb of a gene or overlapping with ATAC peaks in the corresponding cell type. Notably, excluding these regions decreased the performance of cell type classification, but even after removing both of them, most of the major types could still be separated by mCG (Fig. 2N and fig. S13), suggesting the specificity of mCG on genomic regions outside of cCREs and genes may also be functional.

### Patterns of non-CG methylation across cell types

In addition to methylation of cytosines in the canonical CG dinucleotide context, non-CG methylation (referred to as CH methylation or mCH, where H is A, T, or C) has been reported to occur in neurons, glia, and stem cells, and at low or undetectable levels (<1%) in other human tissues (*45*). To better understand the cell types that contain mCH, we first quantified the genome-wide average mCH level across single cells grouped by cell type (Fig. 3A). As expected, neurons had the highest levels of mCH across major cell types (Fig. 3A - rightmost panel). Apart from neuronal populations, we observe modest mCH in glial populations (Glia Oligo, Glia Astro) as well as in muscle-associated cells (skeletal muscle - Mus Skl, muscle fibroblasts - Fibro Mus, cardiomyocytes - Mus Crd). However, it is apparent that mCH largely exists on a continuum outside of neuronal lineages and at low levels (Fig. 3A). Previous reports of mCH showed that this frequently occurs in a CHG context in stem cells (*31*), so we also examined the trinucleotide contexts for mCH across major cell types (Fig. 3B). We observe that on average across major cell types, mCH most frequently occurs in a CAC context. Interestingly, four of the five most frequent CH contexts are in a CpA context, while the lowest seven trinucleotide contexts contain CpC or CpT, suggesting that the presence of a purine following cytosine may increase the likelihood of mCH (Fig. 3B). To validate that the low-level mCH patterns reflected endogenous patterns of methylation and were not the result of incomplete bisulfite conversion, we also examined mCH patterns on spike in lambda phage DNA that is included as a control for every single cell. Indeed, mCH in nearly all base contexts was enriched above what was seen from control lambda phage DNA (fig. S14A), and similarly showed relatively strong enrichment in specific cell types (neurons, glia, and skeletal muscle) and base contexts (fig. S14B). These results suggest that low-level mCH is pervasive across human cell types and is enriched along a continuum between different lineages.

**Fig. 3.**
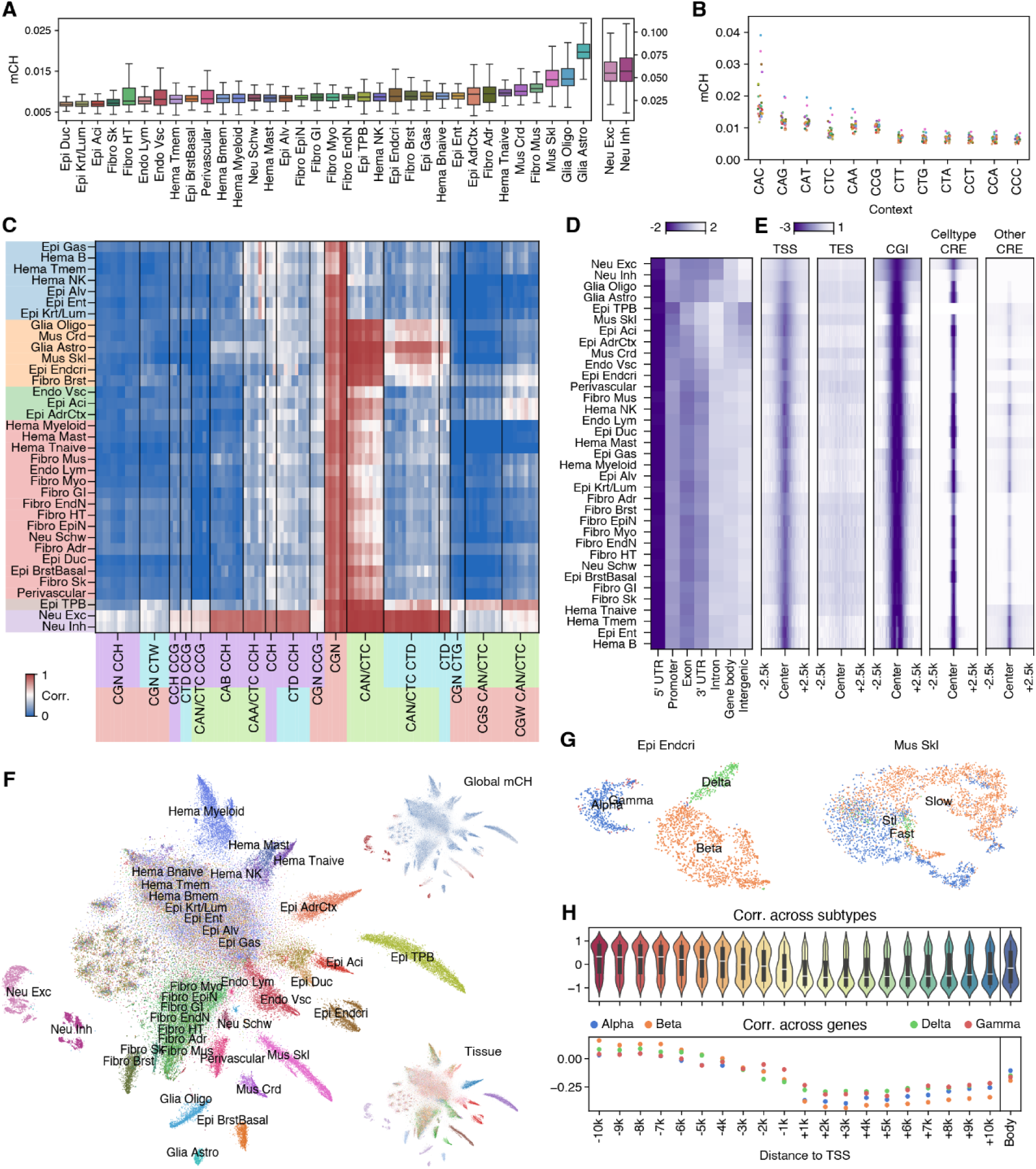
Non-CpG methylation across cell types. (**A**) Distribution of the average non-CG methylation (mCH) level in single cells grouped by major type. The plot is split to allow for the scale to show both neuronal (far right panels) and non-neuronal cell types. (**B**) Average mCH of major cell types excluding neuronal cells grouped by trinucleotide context. (**C**) Correlation of mC between pairwise trinucleotide contexts across 50kb bins in each cell type. Colors show major type grouping (y-axis) or category of trinucleotide contexts (x-axis). (**D**) mCH at different genomic features across major cell types. Values are row-wise Z-scored. (**E**) mCH surrounding different elements. Values are row-wise Z-scored across all five plots together. The last two columns show ATAC-seq defined cCREs present in the given cell type or cCREs remaining after excluding the cCREs of the cell type from cCREs of all cell types. (**F**) t-SNE of all cells (n=86,689) using mCH colored by major types (center), global mCH (top right), or tissue source (bottom right). (**G**) t-SNE of endocrine pancreas epithelial cells (Epi Endocri; n=2,324; left) or skeletal muscle cells (Mus Skl; n=3,973; right) using mCH colored by subtypes. (**H**) Correlations between mCH and expression of differentially expressed genes (DEGs) in Epi Endcri across subtypes for all DEGs (top) or across all DEGs for each cell subtype (bottom). The correlations are calculated using mCH of regions from TSS to different distances on each side of TSS (x-axis) or across the entire gene body (right).

We were also interested in understanding whether patterns of mCH in the genome were non-random. We first calculated the genome-wide correlation of DNA methylation by base context, reasoning that if mCH was non-randomly distributed, we should see a correlation of local patterns of mCH across different trinucleotide contexts. We first compared correlations by trinucleotide context across a range of bin sizes from 50bp to 50kbp (figs. S15 and 16). At small bin sizes (50bp), only methylation in CpG dinucleotide contexts showed consistent correlations across different cell types (fig. S15 and 16). At larger bin sizes, up to 50kbp, strong correlations began to emerge across different CH nucleotide contexts in multiple lineages (fig. S15 and 16). We therefore clustered the correlations between trinucleotide contexts across all major cell types (fig. S15 and 16), and observed six groups of major cell types with similar patterns of CH correlation across base contexts (Fig. 3C and fig. S15). For mCG, we observed a strong correlation between trinucleotide contexts across all major cell types (Fig. 3C), consistent with the known role of mCG in the regulation of cis-regulatory element function in diverse lineages. Similarly, we observe correlations in mCH across a wide variety of trinucleotide contexts in neuronal cell types (Fig. 3C), consistent with the known association of mCH with gene expression. Outside of neurons, we observed correlation in mCH largely in the CTC or CAN trinucleotide contexts (Fig. 3C), which are also the most frequently enriched trinucleotide for mCH across major cell types (Fig. 3B). Notably, CAN/CTC methylation was well correlated across diverse cell types including neurons, different fibroblast populations, and endothelial cells (Fig. 3C - see orange, green, red, gray and purple groups). On the contrary, it was less well correlated in a subset of major types (Fig. 3C - blue group), including some hematopoietic (Hema B, Hema Tmem, Hema NK) or epithelial lineages (Epi Gas, Epi Alv, Epi Ent, Epi Krt/Lum). There is a relatively weak correlation for mCH outside of the CTC/CAN trinucleotide context across most major cell types. This suggests that the mCH can reflect chromatin state differences between different lineages, but this is more pronounced for specific trinucleotide contexts (CAN/CTC) and distinct lineages.

Consistent with the idea that correlations in mCH by base context are suggestive of non-random localization of mCH, we observe that mCH is depleted at regulatory elements across major cell types (Fig. 3D). Specifically, we observe a depletion of mCH at promoters and transcription start sites of genes (Fig. 3, D and E). We noted that major cell types showed different patterns of mCH over promoters and gene bodies. For example, neurons showed relatively low levels of mCH over exons and introns (Fig. 3D), while cells such as skeletal muscle showed higher levels of mCH over these regions. Further, we also observe a depletion of mCH over cCREs, but only when those cCREs are active in the specific major cell types (Fig. 3E - compare two rightmost panels). This suggests that the mCH patterns around regulatory elements may be cell type-specific and harbor information on cell identity. To test this, we clustered cells by mCH and observed the appearance of clusters for many major cell types (Fig. 3F and fig. S17). Similarly, we trained logistic regression models to predict cell type from mCH levels and tested the model’s performance in distinct major cell types (fig. S18A). Some lineages, such as fibroblasts, were globally distinguished by mCH, but specific subsets of fibroblasts were not resolved. In other lineages, we observed that mCH could distinguish relevant cell populations, such as pancreatic islets or skeletal muscle (Fig. 3G and fig. S18A). Taken together, these results suggest that low levels of mCH are present across a wide variety of lineages and retain information regarding regulatory elements and cell identity.

We also explored the relationship between mCH over gene bodies and gene expression to further examine the features separating the cell types with their mCH. Previous studies have shown that mCH in neurons is anticorrelated with gene expression levels, but whether this is true for other lineages is unclear. Therefore, we calculated the correlation between mCH and gene expression across cell subtypes within each major type and observed different relationships between lineages (Fig. 3H and fig. S18B). For example, for both inhibitory and excitatory neurons, mCH and gene expression were anticorrelated at both the regions surrounding the TSS and within the gene body (fig. S18B). In contrast, in cells from the endocrine pancreas, we found that mCH and gene expression were anticorrelated in regions downstream of the TSS but positively correlated in regions upstream of the TSS (Fig. 3H) and showed only a weak correlation with gene expression when considering the entire gene body. Similarly, gene body mCH was mildly anticorrelated with gene expression within skeletal muscle cells but positively correlated with gene expression across NK cell subtypes (fig. S18B). This suggests that the mechanisms relating to mCH and gene activity may vary between lineages and is an important area for future exploration.

### Variability in 3D genome architecture across cell types

Higher-order chromatin structure plays a critical role in regulating diverse processes in the nucleus. We therefore analyzed multiple features of 3D genome architecture, including chromatin contact decay patterns, chromatin loops, and chromatin compartments, across major cell types. Examining contact decay patterns across major cell types, we observe that there are two prominent “peaks” of enrichment of chromatin interactions, one that is most prominent from 100kb-1Mb and the other that is most prominent greater than 10Mb (Fig. 4A). This enrichment in contact decay plots has been described previously, including in previous snm3C-seq studies in human brain samples (*18*, *46*), and has been attributed to the presence of chromatin loops and Topologically Associating Domains (TADs) at shorter distances (100kb-1Mb) and to A/B chromatin compartments at longer distances (>10Mb). Interestingly, the relative enrichment of the short and long-range signals varies across major cell types, with some lineages showing a “domain dominant” phenotype primarily of short-range interactions. In contrast, others show a “compartment dominant” phenotype primarily of long-range interactions. At the same time, many lineages show a mixture of the two (Fig. 4B). Such variability in contact decay patterns has also been previously observed comparing neurons and glial cell types in snm3C-seq data from human brain samples. Neuronal cell types are among the most domain-dominant cells, while five of the six most compartment-dominant lineages are hematopoietic cell types (Fig. 4B). This is consistent with prior observations in human brain data where microglia appear as the most compartment-dominant cells in human brain tissue. Apart from the enrichment of hematopoietic or neuronal lineages, there is no apparent enrichment of other major cell types for compartment or domain-dominant phenotypes either by tissue type or germ layer.

**Fig. 4.**
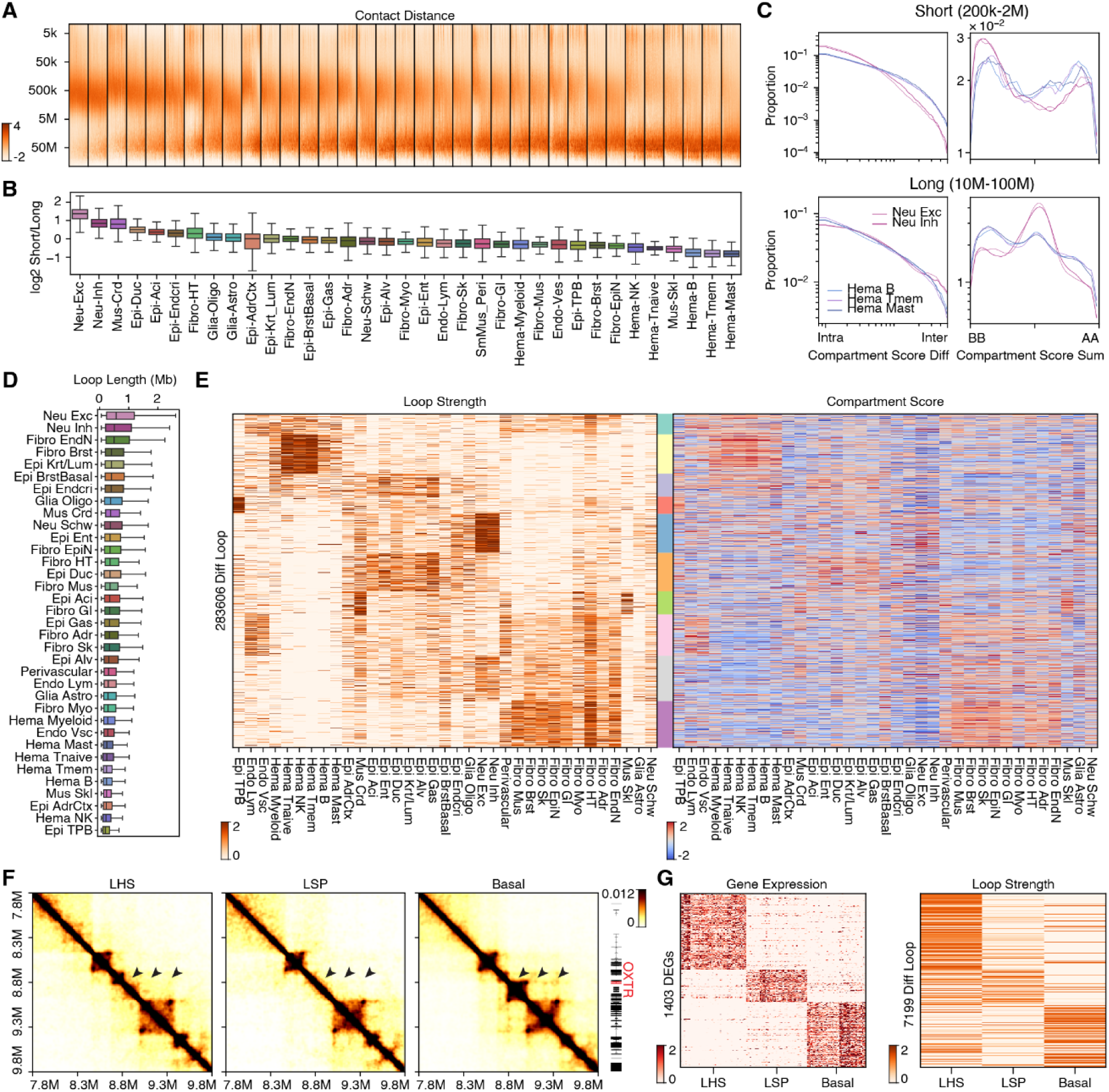
Chromatin contacts across cell types. (**A**) Distribution of contact distances (y-axis) across single cells (n=86,689; x-axis) grouped by major types. (**B**) Ratio of short (200kb-2Mb) to long (10-100Mb) range chromatin contacts among major cell types. Order and labels are the same between (A) and (B). (**C**) Proportion of short (top) or long (bottom) range contacts stratified by the differences (left) or sums (right) of compartment scores at the two interacting regions. (**D**) Distribution of loop lengths by major cell type. (**E**) Interaction strength of differential loops between major types (left) and the compartment scores of loop anchors (right) across major types. Both values are row-wise Z-scored. Colorbar (middle) shows *k*-means clusters of differential loops. (**F**) Browser shot showing subtype specific chromatin loops in breast epithelial cells (Luminal hormone sensitive - LHS; Luminal secretory precursor - LSP; Basal myoepithelial - Basal) at chr3:7,800,000-9,800,000 surrounding basal cell marker *OXTR*. (**G**) Expression level of DEGs whose TSSs are within 2kb of either anchor of the differential loops (left) and interaction strength of differential loops between breast epithelial subtypes (right) across cell types-samples. Values are row-wise Z-scored. Left and right heatmaps share the row orders. When a loop overlaps multiple genes, the loop is repeated in the left heatmap, and vice versa for a gene overlapping multiple loops.

To directly test if the long-range chromatin interactions we observe in compartment dominant lineages are due to A/B compartments, we compared chromatin compartment interactions as a function of genomic distance across major lineages (fig. S19A). Comparing the most domain dominant lineages (excitatory neurons - Neu Exc; inhibitory neurons - Neu Inh) with the most compartment dominant lineages (B-cells - Hema B; memory T-cells - Hema Tmem; Mast cells - Hema Mast), we find that the A/B compartment patterns differ as a function of genomic distance in these lineages. At short distances, the domain-dominant neuronal lineages have relatively more intra-compartment contacts than the hematopoietic lineages (Fig. 4C - upper left panel). This is true for both A-A and B-B compartment interactions (Fig. 4C - upper right panels). In contrast, at longer distances, the hematopoietic cells show relatively stronger within-compartment interactions, while the neuronal lineages appear to be depleted for both intra-B-B and A-A compartment interactions (Fig. 4C - bottom panels). We observed similar patterns across major lineages. Specifically, comparing the ratio of short/long chromatin contacts, domain dominant lineages (higher short/long contact ratios) showed weaker within-compartment interactions (AA or BB) and relatively stronger between-compartment interactions (AB or BA) (fig. S19B). Likewise, calculating the “compartment score” across lineages, we found that the more compartment-dominant lineages (lower short/long ratios) showed stronger compartment strengths (fig. S19B). These observations are consistent with weakened compartment signals in the longer-range (>10Mb) contact decay patterns in neurons and the enriched compartment dominant phenotype seen with long-range interactions in the hematopoietic lineages.

We also examined the presence of chromatin loops across cell types. We observed considerable variability in each lineage’s number (fig. S20A) and size (Fig. 4D) of chromatin loops. Consistent with the domain dominant phenotype in contact decay patterns, neurons were among the lineages that showed the most and longest chromatin loops. At the same time, more compartment-dominant hematopoietic lineages showed shorter chromatin loops (Fig. 4D). We also identified chromatin domains across lineages and observed notably more consistency in domain number (fig. S20B) and size (Fig. S20C) across major lineages. We further compared chromatin loops across lineages and identified 283,606 differential chromatin loops across major cell types (Fig. 4E and fig. S20D). Strikingly, we observed that many differential chromatin loops are shared amongst related cell types (Fig. 4E). The most notable patterns are sets of loops that are shared amongst four primary lineages: fibroblasts, epithelial cells, neurons, and hematopoietic cells (Fig. 4E). We further observe smaller subsets of lineage-specific loops, such as a small group that are specific to trophoblast epithelial cells (Epi TPB) or a set that appears shared between muscle lineages (Mus Crd and Mus Skl) (Fig. 4E). The presence of loops in a specific lineage also appears to correspond with a relative enrichment of those lineages in the A compartment regions, consistent with these being more active regions of the genome (Fig. 4E - right panel).

In addition to major types, we also called loops and identified differential loops between cell subtypes (Fig. 4F), and were able to identify loops that were present in specific lineages. This also allowed us to compare chromatin looping differences with associated gene expression changes. For example, we identified differentially expressed genes between breast epithelial subtypes and were able to show that these are frequently associated with subtype-specific chromatin looping (Fig. 4G; more examples in later sections). Taken together, these results indicate an abundance of variable chromatin loops that differ according to major cell type and subtype lineages and are strongly associated with differential gene expression.

### Association between chromatin conformation and DNA methylation

Having generated DNA methylation and chromatin contact information, we wanted to compare these features to better understand their genomic relationship across cell types. We first computed the correlation between compartment score and methylation level at 100kb resolution. Most major cell types show a negative correlation between mCG and compartment score, aligning with the knowledge that active chromatin is more likely to be A compartment and depleted of mCG (Fig. 5A and fig. S21A). Seven cell types showed a positive correlation between mCG and compartment score, all of which have explicit PMD observed along their genome (Fig. 5A and fig. S21A). This also matches the previous understanding of PMDs that they are likely to be repressive chromatin and have lower mCG compared to the rest of the genome. mCH only showed a negative correlation with compartment scores in the neuronal lineage (Fig. 5B and fig. S21B). This is consistent with our observation in the brain studies that active genes have lower mCH expanding across the whole gene bodies, suggesting different mCH regulation in non-neuronal cell types.

**Fig. 5.**
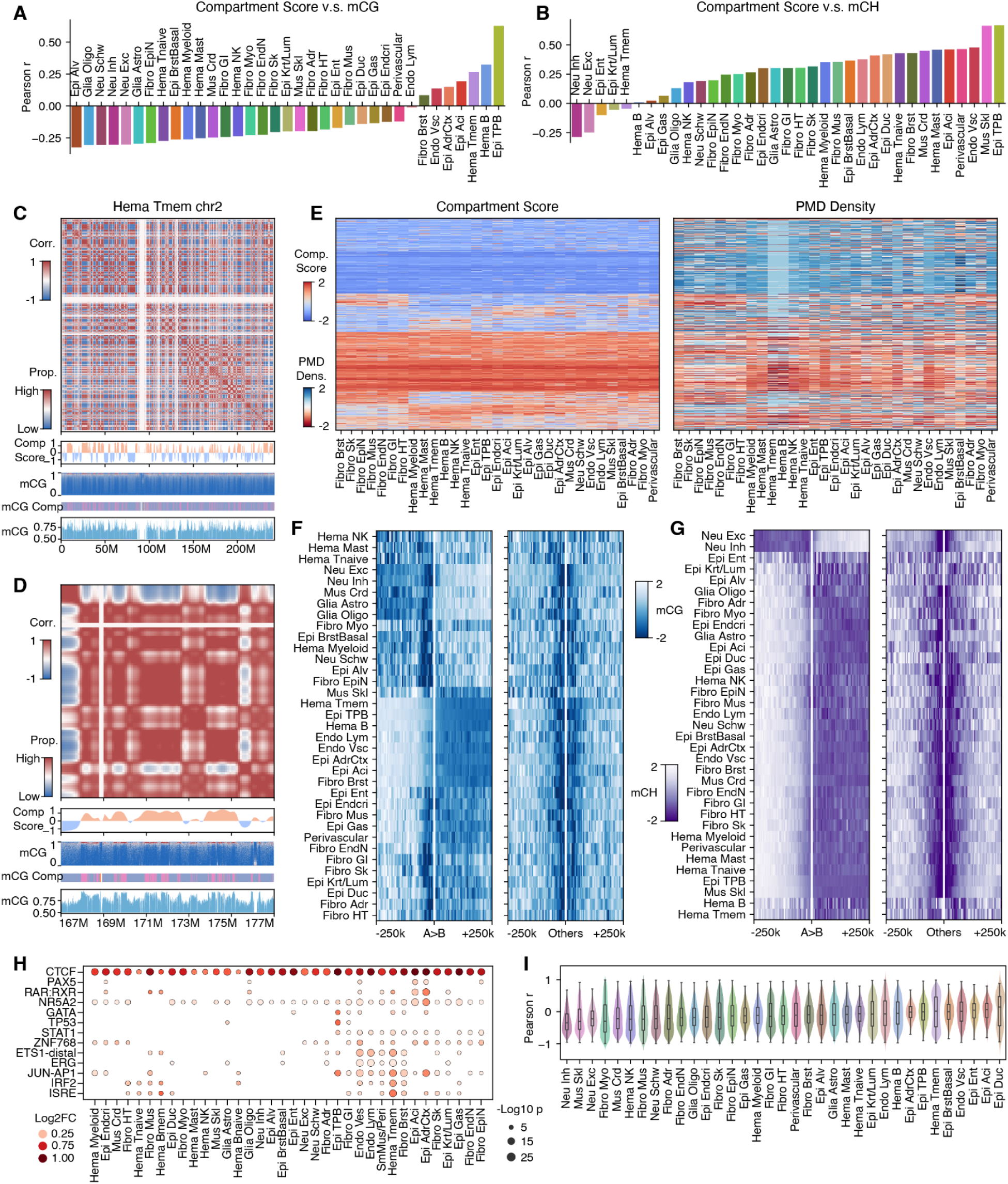
Comparison of DNA methylation and 3D genome structure across cell types. (**A,B**) Correlation between chromatin compartments and mCG (A) or mCH (B) across 100kb bins for all major cell types. (**C,D**) From top to bottom: correlation matrix of distance normalized contact maps of Hema Tmem, compartment score at 100kb resolution, distribution of mCG of individual CpG, methylation compartment, and mCG at 10kb resolution for whole chr2 (C) or chr2:167,000,000-177,000,000 (D). (**E**) Chromatin compartment score (left) and proportion of partially methylated compartment (right; materials and methods) at 100kb resolution across major types. Neurons and glia in the cortex show weak correlations between the two scores and are not shown. (**F,G**) mCG (F) or mCH (G) surrounding chromatin domain boundaries that separates A/B compartments (left) or not (right). (**H**) Enrichment of motifs at loop DMRs versus non-loop DMRs for each major type. The hue represents log2 fold change of proportion of motif-containing DMRs. (**I**) Distribution of correlations between DMRs and differential loops across cell clusters within each major type.

To further investigate the different levels of correlation between mCG and A/B compartment, we compared the 3D genome compartment and the methylation compartment in memory T cells. A clear correspondence of the compartment between the two modalities was observed, with the partially methylated compartment usually matching the B compartment and the fully methylated compartment matching the A compartment (Fig. 5C). When zoomed in to a 10Mb region of the genome, the methylation compartment provides higher-resolution segregation of the genome (Fig. 5D). We further expanded the comparison to all major cell types and observed that most regions of the genome were either stable as A or B compartments across major cell types, with a subset of regions of the genome switching between A and B compartments in select lineages (Fig. 5E - left panel). Comparing the A and B compartment patterns with DNA methylation, we also observed a strong association between B compartments and regions considered as partially methylated DNA methylation compartments. The regions that were constitutively B compartment across lineages were consistently identified as part of the partially methylated compartment across lineages. In cases where there are lineage-specific switches of A/B compartments, we observe that in the lineages where a region is identified as B compartment, it will similarly be identified as the partially methylated compartment (Fig. 5E). The one notable exception to this is in neurons and glial cells in the brain, where there was not an obvious correlation between B compartment regions and the partially methylated compartments. With the exception of neural lineages, these results strongly suggest that the B compartments are the same as the partially methylated compartment we identify through clustering (fig. S8E), indicating that B compartment regions of the genome may have fundamentally distinct mechanisms for maintaining DNA methylation in the genome.

We also compared DNA methylation patterns at chromatin domain boundaries in the genome, stratifying domain boundaries according to whether they separate A and B compartment regions. We performed this analysis for both mCG (Fig. 5F) and mCH (Fig. 5G). The relationship between domains and mCG patterns varies by lineage. Across nearly all lineages, domain boundaries demarcate regions showing enrichment and depletion of mCG (Fig. 5F). Further, most lineages show a relative focal depletion of mCG at or immediately adjacent to the domain boundary (Fig. 5F and fig. S21C). However, when considering domain boundaries that separate A and B compartment regions, the association of mCG with A/B compartments varies between lineages (Fig. 5F and fig. S21C). Consistent with the correlation analyses, lineages that show evidence of PMDs, including trophoblast and some hematopoietic lineages, show lower levels of mCG in the B compartment side adjacent to domain boundaries (Fig. 5F and fig. S21C), while other lineages, including neurons and glia, show a depletion of mCG within A compartment side (Fig. 5F and fig. S21C). On the contrary, patterns of mCH and their relationship with compartments and domain boundaries are simpler. Nearly all lineages show enrichment of mCH in A-compartment regions except for neuronal lineages, which show enrichment of mCH in B-compartment regions (Fig. 5G and fig. S21D). Interestingly, across all lineages, mCH is depleted at domain boundaries that separate compartments of a similar type (Fig. 5G). This is consistent with the general depletion of mCH we observe over cCREs (Fig. 3E), suggesting that lack of mCH at regulatory elements may be a universal phenomenon across human cell types. The difference in the distribution of mCH in neurons and non-neuronal lineages is perplexing, as neurons show much higher levels of mCH than most lineages. Why mCH shows distinct associations with compartments and gene expression in neuronal versus non-neuronal lineages remains unclear.

To investigate the association between the two modalities at a higher genomic resolution, we compared DMRs between cell subtypes within each major type with chromatin loops. DMRs are enriched at loop summit anchors in all major types (fig. S22A). We performed motif enrichment analysis on DMRs associated with loops against those not associated with loops to nominate TFs potentially regulating the chromatin loop formation and maintenance. The analyses showed that the CTCF motif is enriched in almost all the major types, supporting their crucial role in higher-order chromosome structure regulation (Fig. 5H). Intriguingly, we also identified other lineage-specific TF, including IRF2 and ISRE in the immune lineage (*40*), NR5A2 in the pancreas acinar cells (*47*) and GATA factors in the trophoblasts (*48*) (Fig. 5H). Our analyses indicate they could also contribute to the cell type-specific 3D genome structure, possibly through mechanisms such as modulating CTCF accessibility to chromatin.

We further identified differential loops between subtypes and compared them to the DMRs. In most major types, DMRs are more enriched at the loops showing higher variability across subtypes (fig. S22B). Next, we calculated correlations between DNA methylation and chromatin contact across subtypes at differential loops overlapping DMRs. We observed variable relationships between mCG at DMRs and chromatin loops (Fig. 5I). Some lineages, such as neurons, showed strong anticorrelations between DNA methylation and chromatin loops at loop anchors, consistent with the notion that depletion of DNA methylation occurs over active regulatory elements. In contrast, other lineages showed little clear relationship between DNA methylation and chromatin looping, or in some cases, showed bimodal distributions (Fig. 5I), with some loops being positively correlated with DNA methylation while others are negatively correlated with DNA methylation (i.e., Hema Tmem, Epi Duc). Some of this may be related to the presence of partially methylated domains, as the lineages that show the least correlation between DMRs and chromatin looping (Fig. 2C - Hema Tmem, Epi TPB, Epi Aci) also show the strongest presence of PMDs in the genome. This is not a product of the clustering method or data modality, as the same lineages show consistently little correlation between DNA methylation and chromatin looping whether we cluster by DNA methylation, 3D genome organization, or jointly between the two data modalities (fig. S23).

### Discrepancies between DNA methylation and chromatin contact

Overall, we observed frequent correlations between patterns of DNA methylation and 3D chromatin architecture across major cell types. However, we also observed distinct differences, in particular, related to the clustering of major cell types into cell subtypes (Fig. 1F, and figs. S24 and 25). To better understand these differences, we compared sub-clustering using DNA methylation or chromatin contacts within major cell types. We quantified the degree of similarity between the DNA methylation and chromatin compartment clusters using the Adjusted Rand Index (ARI - Fig. 6A). We observed considerable variability in the consistency of clusters between the two modalities. Some lineages showed strong agreement between the two modalities, with neurons, endothelial vascular cells, and distinct subsets of fibroblast, epithelial, and hematopoietic cells showing good agreement in clusters (Fig. 6A). In contrast, other lineages showed little to no agreement in clusters between the two modalities (Fig. 6A - rightmost panels). These results suggest that the association of DNA methylation and 3D genome organization with cellular states may vary across lineages.

**Fig. 6.**
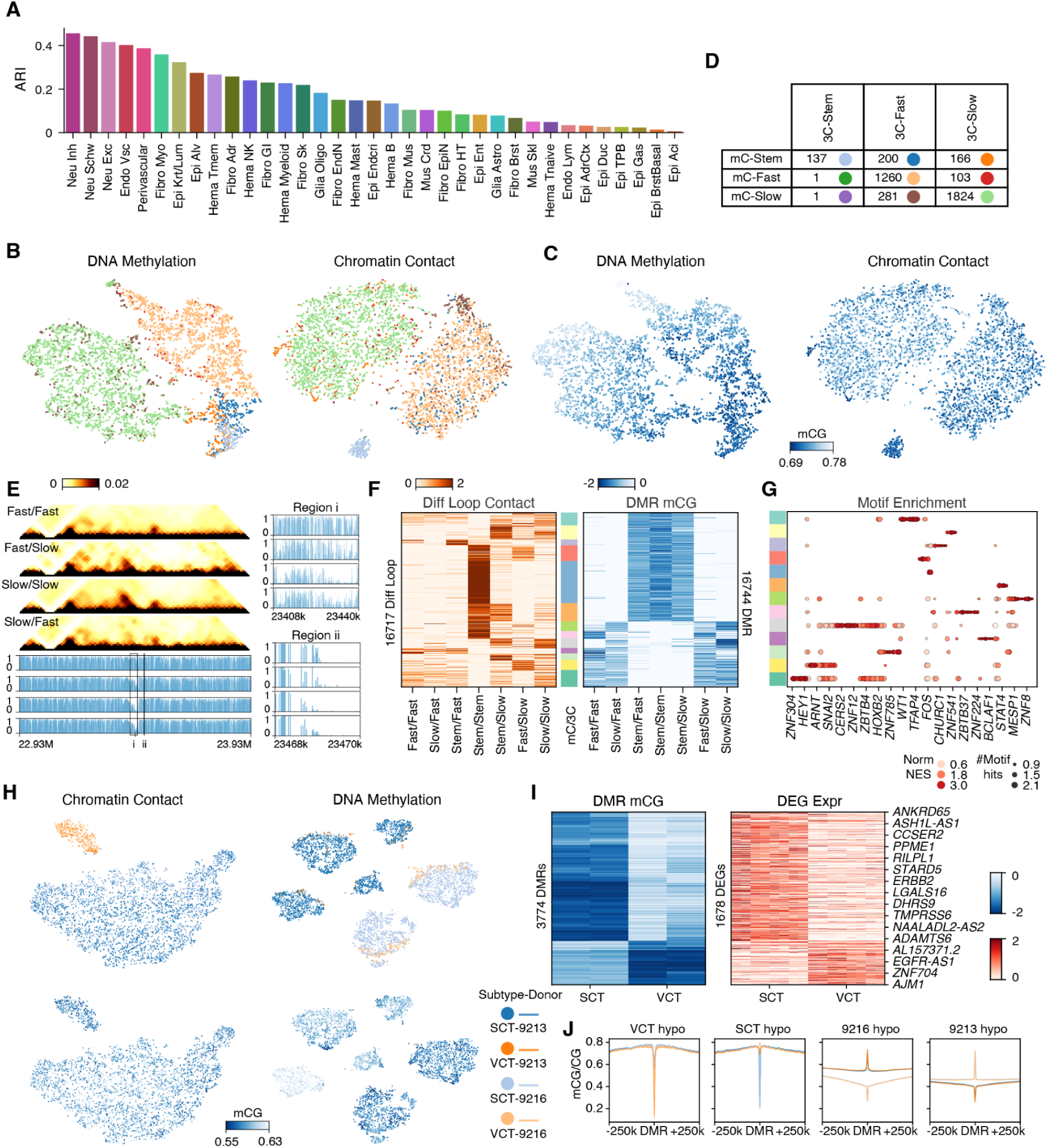
Differences between DNA methylation and 3D genome architecture defined cell types. (**A**) Adjusted Rand Index (ARI) quantifying the consistency between mCG and chromatin contact based clustering of each major type into cell subtypes. Higher ARI corresponds to greater agreement in clusters. (**B,C**) t-SNE of Mus Skl (n=3,973) using mCG (left) or chromatin contact (right) colored by cell groups (B) or global mCG (C). (**D**) Confusion matrix of subtypes defined with mCG (mC-) versus chromatin contacts (3C-). Cell group colors are shared in (C) and (D). (E) Browser shot of chromatin contacts and mCG with the same cell group orders at chr14:22,930,000-23,930,000 (left) surrounding slow twitch fiber marker *MYH7* and zoomed-in view of DMRs (right) of the regions in the boxes. (**F**) Interaction strength of differential loops between cell groups (left) and mCG of DMRs at either anchor of the differential loops (right) across cell groups. Values are Z-score normalized within each row. Colorbar (middle) shows *k*-means clusters of loop-DMR pairs. (**G**) Motif enrichment of DMRs in each group of loop-DMR pairs in (F). (**H**) t-SNE of Epi-TPB (n=5,228) using chromatin contact (left) or DNA methylation (right) colored by subtypes-donors (Villous cytotrophoblast - VCT; syncytiotrophoblast - SCT; top) or global mCG (bottom). (**I**) mCG of DMRs between subtypes (left) and expression of DEGs whose TSSs are within 2kb of DMRs (right) across subtypes-donors. Values are row-wise Z-scored. (**J**) mCG in subtypes-donors at flanking regions of subtype DMRs (left) or donor DMRs (right). Colors are shared between (H) and (J).

We wanted to explore the basis for these discrepancies in greater detail. One lineage with poor agreement between clustering was in skeletal muscle (Mus Skl). Comparing the clustering between DNA methylation and chromatin contacts, both data modalities identify distinct clusters corresponding to satellite cells (muscle stem cells - blue), and slow (green) or fast twitch (orange) fibers (Fig. 6B), but the clusters from DNA methylation appear to separate along a continuum of mCG levels (Fig. 6C). In contrast, chromatin contact data showed the presence of three distinct clusters without obvious DNA methylation gradients (Fig. 6C). The two modalities showed a significant mismatch between the subtype assignment (Fig. 6D). Subsets of methylation annotated stem cells were assigned to mature cell populations with chromatin contacts. The subsets of cells annotated as mature by 3C but stem cells by mCG were mixed throughout the chromatin contact-defined mature cell population instead of positioned on one side of the embedding (Fig. 6B), suggesting their genome architectures have become indistinguishable from other mature cells. On the contrary, almost no cells annotated as mature cells by mCG were annotated as stem cells by 3D genome (Fig. 6D). This suggests that there is a population of muscle fibers that, at the chromatin contact level, resemble mature myofibers but, at the DNA methylation level, resemble stem cells. This finding is consistent with our previous study in the developing human brain, where we observed that the dynamic of 3D genome structures usually happened earlier than DNA methylation (*46*).

In addition, a considerable number of mature myofiber cells were also clustered differently by the two modalities (Fig. 6, B and D). We examined the locus of a slow-twitch muscle marker gene, MYH7. Indeed, the 3D genome structure of the cell group assigned as slow twitch by mCG but fast twitch by chromatin contacts (fast/slow) show weaker chromatin loops between the gene promoter and upstream regions, but a stronger depletion of mCG (Fig. 6E). To analyze this more systematically, we identified differential loops and DMRs between the satellite, slow, and fast twitch fibers (Fig. 6F and fig. S26A). We observe that the differential loops are most distinct between the stem cell population and the mature myofiber clusters (Fig 6F). Similarly, these regions show a depletion of mCG in the stem cell populations (Fig. 6F - right panel). However, in the cells that differ between chromatin contacts and DNA methylation (Stem/Fast or Stem/Slow), they have lost the stem cell chromatin loops and begin to display chromatin loops reminiscent of differentiated cells while retaining the methylation signatures of the stem cell populations (Fig. 6F - compare left and right panels for Stem/Fast or Stem/Slow cells).

To determine the factors contributing to the differences in 3D genome and DNA methylation states in these lineages, we performed motif enrichment analysis at DMRs overlapping differential chromatin loops (Fig. 6G). We found that DMRs hypomethylated in the three methylation-defined stem cell clusters (group 1, 3, 5, 6, 12) are more related to muscle fiber differentiation. For example, *MYF5* is involved in the early commitment of satellite cells to the myogenic lineage(*49*). *SIX1* and *MYF6* are both essential for later-stage fast-twitch muscle development(*50–52*). In contrast, DMRs hypomethylated in the differentiated muscle types ( group 0, 2, 4, 7, 9, 10, 11) are more enriched of TFs involved in muscle cell regeneration and adaptation to outside stimulus. For example, PAX7 is a marker for stem cell population (*49, 53*), MYC is increased after exercise (*49, 53*), and CEBPB is induced upon injury (*54*). These TFs potentiate the differentiated muscle cells for fast adaptation to outside stress. Interestingly, we found *FOXO1* and *NFKB1* in cluster 8, which contains loops that exist only in 3C-defined stem cells and DMRs in methylation-defined stem cell clusters. Both TFs are negative regulators for muscle differentiation (*49*, *53*, *55*, *56*), suggesting that cluster 8 may be critical to the maintenance of muscle stem cell population. These observations may reflect differences in the dynamics of 3D chromatin organization and DNA methylation during terminal differentiation in some lineages.

The inconsistency between cell type classification with different modalities is also observed between other cell subtypes, including the Non-myelinated versus Myelinated Schwann cells, the Memory B cells versus Plasma cells, and Alveolar Type 1 versus Type 2 cells in Lung. In Schwann cells, we also observed relative continuous changes from Non-myelinated cells to Myelinated cells in the mCG embedding (fig. S26B - left), while two discrete clusters of the two subtypes in the chromatin contact embedding (fig. S26B - right). A cluster of cells (c21-c7) clustered with Myelinated cells based on mCG while clustered with Non-myelinated cells with chromatin contacts (fig. S26, B and C). Further examination of the DMRs and differential loops between the clusters showed a similar pattern as that of the muscle cells. For the same genome locus harboring DMRs and differential loops, cluster c21-c7 showed methylation patterns similar to the Myelinated cells while chromatin looping patterns similar to the Non-myelinated cells (fig. S26D).

In contrast to these cell types, other lineages show almost no correlation at all between the DNA methylation and chromatin contact-defined clusters. For example, within placenta epithelial cells, we observe two clusters from chromatin contact data and six clusters for DNA methylation (Fig. 6H). The DNA methylation clusters are largely driven by donor and global methylation level differences (Fig. 6H, right), which are not segregated in the chromatin embedding (Fig. 6H, left). To further test which clustering better represents cell type differences, we identified DMRs between the two sets of clusters. Intriguingly, the DMRs between clusters from chromatin embedding were centered around the marker genes between villous cytotrophoblast (VCT) and syncytiotrophoblast (SCT), and genes highly expressed in the same cell type showed consistent changes of mCG between cell types (Fig. 6I). These DMRs have strong mCG at their flanking regions (Fig. 6J), suggesting they are usually found in the fully methylated compartment discussed in Fig. 2. Contrarily, the DMRs between methylation-based clustering show lower flanking mCG (Fig. 6J), suggesting that they are more likely to be located in the partially methylated compartment. Taken together, these indicated that the 3D genome embedding reflected the cell type differences more precisely in Trophoblast. In contrast, methylation embedding was dominated by large global methylation differences in the partially methylated compartment. Similar to these, the same pattern was observed in Epithelial cells in Stomach and Intestine (fig. S26, E to H). This indicates that global patterns of DNA methylation may lack information regarding cell type but that local methylation patterns over specific genes may be informative. This may suggest that DNA methylation is still playing a critical role in the regulation of these genes and lineages but that the variation in DNA methylation globally across single cells is higher and is not primarily defined by cell type.

## Discussion

DNA methylation and 3D chromatin architecture are critical aspects of proper gene regulation, but we currently lack comprehensive information about how these features vary across human cell types. To address this, we have used single-nucleus methyl-3C sequencing (*22*) to simultaneously profile DNA methylation and 3D chromatin architecture in single cells across human tissues. Our study has generated DNA methylation and 3D chromatin architecture profiles for 35 major cell types and 206 cellular subtypes across 16 human tissues. This has revealed the landscape of DNA methylation and 3D chromatin architecture across human cell types at an unprecedented scale and has revealed novel aspects of how these features are related between cell types.

In examining the presence of mCG across cell types, we have identified the presence of features that we term “DNA methylation compartments”. This stems from examining the patterns of Partially Methylated Domains (PMDs) (*31*) in different cell types, where the degree of partial methylation over PMDs varies across lineages. Indeed, we observe that between related cell types, regions called PMDs in one cell type will show a notable decrease in methylation in other cell-related types (fig. S8C) compared to non-PMD regions of the genome. The patterns of methylation compartments are highly correlated with patterns of A/B compartments (*57*) based on 3D genome interaction data, with the PMD-like compartments being strongly associated with the repressive, heterochromatin, B-compartment regions of the genome (*57*, *58*). Interestingly, the compartment patterns from DNA methylation-based clustering appear to be more fine-grained than the compartments from 100kb bin Hi-C-based analysis. This is consistent with recent observations using ultra-deep (33 billion read) Hi-C sequencing that A/B compartments may be smaller than previously realized (*59*). One advantage of using DNA methylation compartments in this regard is that they may reveal the fine-grained nature of chromatin compartmentalization across cell types more cost-effectively.

The association between the PMD-like compartments and the B-compartment regions of the genome suggests that the mechanisms that maintain DNA methylation vary by genomic context, in particular for heterochromatic regions of the genome. Previous studies have suggested that late replicating regions of the genome, which are also associated with the B-compartment, progressively lose DNA methylation over many cell divisions to create PMDs (*60*). Our data suggest that these mechanisms may also be present in normal or non-dividing cells, with the PMD-like DNA methylation compartments showing lower levels of mCG, albeit with a smaller effect compared to cells with strong PMDs. These observations would fit with a model where the PMD-like DNA methylation compartments are “primed” to become PMDs if a cell type is stimulated to divide rapidly, as might occur between naive lymphoid cells and mature memory lymphoid cells, where we observe some of the clearest changes in mCG over PMD-like compartments between cells.

In addition to canonical mCG, DNA can be methylated in CH contexts in mammalian cells (*31*), previously described as being prominent in neurons and embryonic stem cells (*45*). We compared the levels and patterns of mCH across cell types. Most cells, apart from neuronal lineages, show low levels of mCH. Despite this, the mCH patterns are non-random. Within most cells, we observe local correlation in mCH patterns according to specific trinucleotide contexts, and indeed we find mCH is specifically enriched or depleted at genes and regulatory elements across most cell types. Further, mCH patterns alone can distinguish many of the major cell types we identified in this study (Fig. 3F), suggesting that mCH reflects cell type-specific chromatin states. Mechanistically, within neurons, the high levels of mCH over genes have been shown to influence the binding of the repressor MECP2 (*61*), but whether the lower levels of mCH play a similar role in regulation of chromatin state and gene regulation across diverse cell types remains unclear.

We also characterized patterns of 3D genome organization across an unprecedented number of distinct cell types, identifying patterns such as chromatin compartments, domains, and loops in both major cell types and cell subtypes. For chromatin loops, we observe extensive differences between cell types that are frequently correlated with cell type-specific gene expression of neighboring genes. Interestingly, we also observe that many loops are shared amongst related lineages across tissues, with notable similarities of chromatin loops within hematopoietic cells, epithelial cells, neurons, or fibroblasts (Fig. 4E). These results may suggest that chromatin loops play outsized roles in establishing the gene regulatory programs across major axes of cell type specification. Further, examining the distribution of chromatin interactions as a function of genomic distance, we also observed that distinct lineages can show “compartment dominant” or “domain dominant” phenotypes or a mixture of the two (Fig. 4, A to D). Similarly, prior single-cell 3D genome studies in the human brain (*46*) also show a continuum of compartment domain or domain dominant cell types. Interestingly, the degree of domain dominance versus compartment dominance appears to be anticorrelated, with the most domain dominant cells showing weak compartments and the most compartment dominant cells showing weak domains (Fig. 4A). The basis for compartment versus domain dominance is unclear but is reminiscent of the anticorrelated relationship between chromatin loops and compartments that has been revealed upon depletion of the cohesin complex (*62–64*). These experiments indicate that cohesin loop extrusion both promotes chromatin loop formation while diminishing the presence of chromatin compartments. It is therefore tempting to speculate that the compartment or domain dominant phenotypes may result from differences in global rates of chromatin loop extrusion. How this would be regulated is unclear, but recent evidence has suggested that loop extrusion may be preferentially initiated in euchromatic regions and arrested in B-compartment regions in lymphocytes (*65*). Therefore differences in the type or degree of heterochromatin within a lineage may be a contributing factor.

Comparing DNA methylation and 3D genome structure, we observe multiple ways in which these features correlate with each other throughout the genome, including at the levels of chromatin compartments and domains. Interestingly, we also observed cases where these two features give different pictures of the composition of cells within a tissue. Notably, when we cluster cells either separately or jointly based on DNA methylation and chromatin contacts, we observe some cell types with divergent classifications between the two features. For example, within skeletal muscle, 3D chromatin-based clustering largely separates cells by cell state, whereas DNA methylation appears to separate cells along a continuum from progenitor satellite cells to mature muscle fibers (Fig. 6, B and C). This suggests that DNA methylation may mature over longer time scales in these cells, such that it is reflected in the maturation state of the cells. In contrast, the 3D chromatin classifications appear to reflect more categorical cell type definitions. Understanding the mechanism that gives rise to these divergent patterns of DNA methylation and 3D genome will be an important area for future studies.

## Materials and Methods

### Human Subjects

Among the 32 samples profiled in the study (table S1), 21 samples from 12 tissues were acquired by the ENTEx collaborative project via the GTEx (*66*) collection pipeline (*67*). All human donors were deceased, and informed consent was obtained via next-of-kin consent for the collection and banking of deidentified tissue samples for scientific research. Donor eligibility requirements were as described previously (*67*), and excluded individuals with metastatic cancer and individuals who had received chemotherapy for cancer within the prior two years.

Additional samples for the adrenal gland, heart left ventricle, and heart right atrial appendage were obtained from previously collected tissue from the NIH Epigenomics Roadmap Program (sample STL003)(*68*). In addition, the two motor cortex samples were obtained as part of the BRAIN Initiative Cell Census Network(*69*). Pancreatic islet samples were obtained from Prodo labs and purified using density centrifugation. Peripheral blood mononuclear cells (PBMC) were obtained from AllCells.

The two placenta samples were obtained as part of the ENCODE4 sampling effort. Detailed metadata and biosample-specific information are available on the ENCODE Project website (encodeproject.org) under the following biosample accession numbers: ENCDO384FEJ (https://www.encodeproject.org/human-donors/ENCDO384FEJ) ENCDO023YQQ (https://www.encodeproject.org/human-donors/ENCDO023YQQ). Nuclei isolation was performed according to the “ENCODE Nuclei Isolation from Tissue for 10x multiome” protocol (https://www.encodeproject.org/documents/d18bad0e-9a86-40f0-8ded-2c74072910e5).

### Nuclei Isolation

The nuclei isolation protocol is described in detail in https://www.protocols.io/view/snm3c-seq3-kqdg3x6ezg25/v1. Specifically, human tissue samples were pre-ground in liquid nitrogen and stored at −80°C in aliquots of 50–75 mg. For nuclei isolation, each tissue aliquot was transferred to a Dounce homogenizer containing 2.4 mL of lysis buffer (NIMT: NIM [250 mM Sucrose, 25 mM KCl, 5 mM MgCl₂, 10 mM Tris-Cl pH 8.0] supplemented with 0.1% Triton X-100, 1 mM DTT, and protease inhibitor at a 1:100 dilution). The lysate was then divided into two 2 mL Eppendorf tubes and centrifuged at 1,000 × g for 10 min at 4°C using a swing-bucket rotor. After discarding the supernatant, the nuclear pellet was resuspended in ice-cold DPBS supplemented with RNase inhibitors. The resuspended pellets from both tubes were combined to a final volume of 1 mL per sample.

### Chromosome conformation capturing (3C)

The 3C protocol is based on the 3C kits protocol from Arima Genomics and is described in detail in https://www.protocols.io/view/snm3c-seq3-kqdg3x6ezg25/v1.

#### Crosslinking

Nuclei samples were divided into tubes, each containing 1–2 million nuclei in up to 1 mL of DPBS. Crosslinking was initiated by adding 57 µL of 37% formaldehyde (final concentration: 2%), followed by mixing via inversion (10 times). The samples were incubated at room temperature for 5 min with occasional inversion. To quench the reaction, 91.9 µL of Stop Solution (2.5 M Glycine) was added, followed by additional mixing via inversion (10 times) and a further 5 min incubation at room temperature. The samples were then placed on ice for 5 min. The nuclei were pelleted by centrifugation at either 2,500 × g for 5 min or 1,000 × g for 10 min. The supernatant was carefully removed to avoid residual liquid, and the pellet was resuspended in 1 mL of 1× PBS. This washing step was repeated to ensure complete removal of formaldehyde and glycine. Samples were either processed immediately or stored at −80°C for up to 5 days.

#### 3C Nuclei Conditioning

Each crosslinked nuclei sample was resuspended in 20 µL DPBS. Next, 24 µL of Conditioning Solution (0.5% SDS) was added, and the mixture was gently pipetted to ensure thorough mixing. Samples were incubated at 62°C for 10 min. If using a thermal cycler, the lid temperature was set to 85°C. Following incubation, 20 µL of Stop Solution 2 (10% Triton X-100) was added, and the mixture was gently pipetted to mix. Samples were then incubated at 37°C for 15 min, maintaining a lid temperature of 85°C if using a thermal cycler.

#### 3C Enzymatic Reactions

Mix the sample with 28 µL of the master mix containing enzymes H1, H2, and Cutsmart Buffer (NEB) by gently by pipetting and incubate as directed for 1 hour at 37°C followed by inactivation at 65C°.

#### Ligation

Digested chromatin ends were ligated by adding 82µL of the ligation master mix containing buffer C and enzyme C. Samples were gently mixed by pipetting and incubated at room temperature for 15 minutes. DPBS containing RNase and protease inhibitors was added to bring the volume up to 1 mL (approximately 800 µL added), and the sample was gently mixed. Nuclei were pelleted by centrifugation at 2,500 × g for 5 minutes at 4°C, and the supernatant was carefully removed. The pellet was resuspended in 250 µL of DPBS supplemented with 1% BSA, followed by the addition of 750 µL more to reach a final volume of 1 mL.

To stain the nuclei, Hoechst 33342 was added (5 µL per 1 mL sample, final dilution 1:1000). The Hoechst stock was first diluted 1:10 (5 µL in 45 µL DPBS) before use. The stained sample was incubated for at least 5 min, then filtered through a 40-µm cell strainer and transferred to a polypropylene tube for sorting. A 10 µL aliquot was taken for cell counting using the Bio-Rad TC20 automated cell counter. The final sample was then filtered through a 40 µm cell strainer and transferred to a polypropylene tube for fluorescence-activated cell sorting (FACS). A 10 µL aliquot was taken for a second nuclei count, if possible.

#### FACS sorting

Nuclei were sorted into 384-well plates pre-loaded with a digestion buffer containing proteinase K. Sorting was performed using a BD Influx cell sorter in single-drop mode, employing 2N gating based on Hoechst staining to isolate intact nuclei. A total of 8–10 plates were prepared per sort. After sorting, nuclei were spun down in the plate, and proteinase K lysis was carried out using a thermal cycler program (20 minutes at 50°C). Plates were stored at −20°C prior to library preparation.

### Library preparation and sequencing

The library preparation protocol is described in detail in https://www.protocols.io/view/snm3c-seq3-kqdg3x6ezg25/v1. This protocol was automated using a Beckman Biomek i7 and Tecan Freedom Evo instrument to facilitate large-scale applications. The snmC-seq3 and snm3C-seq libraries were sequenced on an Illumina NovaSeq 6000 instrument, using one S4 flow cell per 16 384-well plates and using a 150 bp paired-end configuration. The following software were used during this process: BD Influx (v.1.2.0.142; for flow cytometry), Freedom EVOware (v.2.7; for library preparation), Illumina MiSeq control (v.3.1.0.13) and NovaSeq 6000 control (v.1.6.0/RTA, v.3.4.4; for sequencing), and Olympus cellSens Dimension 1.8 (for image acquisition).

### Identification of false positive signals in 3C contact matrices (exclusion list)

We have observed unusually high signals at certain near regions, both near diagonal and off-diagonal (fig. S1A). These regions often overlap with highly repeated regions and ENCODE exclusion regions and can confuse the downstream domain and loop-calling tasks. We used the m3c mapping pipeline in yap (https://github.com/lhqing/cemba_data) (*70*) to process the snmC-seq3 data (without chromosome conformation capturing), and observed the same signals (fig. S1A), suggesting that these signals are likely due to mapping errors of the bisulfite converted reads. We observed highly consistent signals between split reads and unsplit reads (fig. S1A), suggesting these signals are not due to the read-splitting process of our mapping pipeline. We also evaluated the consistency of these signals between samples from different donors and different cell types, and observed highly correlated results between two different donors of MTG snmC-seq3, two donors of M1 snm3C-seq, and between L2/3 excitatory neurons, inhibitory neurons, and muscle stem cells (fig. S1A).

Given these observations, we used snmC-seq3 data (without chromosome conformation capture) from two MTG samples and 12 PBMC samples to determine the loci-pairs across the genome that have high signals not resulting from ligation. We mapped snmC-seq data of 768 cells from each sample with yap and “m3c” mapping mode. Because these data are generated from experiments without crosslinking and ligation of interacting DNA, the chromatin contacts detected in these cells should come from mapping errors or random chimeric reads (possibly due to template switching by Klenow polymerase during first strand synthesis). We then summed up the contact matrices at 10kb resolution of the cells from the same donor and normalized the total number of contacts in each matrix to the average depth of all 14 samples. The 10kb bin-pairs with median signal >1 are selected as exclusion list, and the contacts with both ends overlapping with the exclusion list are removed from the contact file of each single cell after snm3C-seq data mapping.

The snmC-seq3 data mapped with m3c mapping pipeline were also used to decide the lower distance threshold of read pairs to be considered as a contact. snmC-seq3 data show a peak of alignments at short ranges (<2.5kb), consistent with the fact these originate from continuous DNA molecules, while the snm3C-seq method that assays chromatin contacts shows enriched alignments at larger genomic distances. The differences in distance dependent decay patterns in the two methods was used to define a cutoff (2.5kb) for classifying a read pair as a chromatin contact (Fig. S1B).

### Workflow for clustering cells with mCG and chromatin contacts

The multi-modality multi-level cell clustering was achieved with the following steps. 1. Compute mCG embedding of each cell. 2. Integrate the mCG embedding between donors. 3. Compute 3C embedding of each cell. 4. Integrate the 3C embedding between donors. 5. Combine mCG and 3C embedding. 6. Perform consensus clustering on joint embedding.

The whole pipeline consists of iterations of these steps.

a. For each tissue and each modality, separately compute the embedding and perform integration between donors and clustering (Modality tissue L1 cluster).
b. For each modality tissue L1 cluster, identify integration anchors between the two donors.
c. For all cells, go through steps 1-6 using all the anchors identified in step b, and annotate the joint consensus clusters into major types.
d. For each major type, go through step 1-6 using all the anchors identified in step b, and identify initial clusters.
e. Merge the initial clusters through supervised models to identify 432 final clusters.
f. Merge the final clusters to identify 206 subtypes.

Details of the clustering and integration steps are described below.

### Cell embedding with mCG

To increase the sensitivity of cell type identification, we developed a method for clustering single-cell methylation data using 5kb genomic bins as features. Neuronal cells have highly diverse methylation patterns, including the mCH signatures across the whole gene bodies and the large super-enhancer-like regions with differential CG methylation across cell types. Together, these make it possible to use methylation levels at 100kb resolution to resolve cell types at high granularity in our previous publications. Contrarily in other tissues where fewer long and specific features were explicitly observed, classifying cell types using DNA methylation data only remains challenging. Existing methods often focus on the methylation level of promoters or predefined enhancers. However, recent studies have also revealed limited methylation diversity near TSSs across cell types, while relying on annotated enhancers limits the generalizability of the analysis in the samples without histone data. To develop a universally adaptable framework, we use 5kb genomic bins as features to perform dimension reduction, with the rationale to capture methylation dynamics at shorter regulatory elements. Given the coverage of typical scDNAme assays (millions of reads per cell), 90% of 5kb bins across the mammalian genome are covered by at least one read, making the full cell-by-5k bin matrix a dense matrix and challenging to be saved either in memory or on disk. To alleviate this issue, ALLCools compute a hypo-methylation score in each 5kb bin for every single cell, assuming regulatory elements are usually hypo-methylated. The score accounts for the methylation level and coverage with a binomial model. Only the bins with a score > 0.9 are stored, while all other bins are saved as 0. The resulting cell-by-bin matrix is a sparse matrix with ∼2% non-zero values, considerably reducing computation cost and disk usage. After generating a hypo-methylation score matrix, we binarize the matrix with a threshold and perform latent semantic indexing (LSI) for dimension reduction, similar to scATAC data analysis.

Specifically, in a single cell *i*, we modeled its mCG base call *M_ij_* for a 5kb bin *j* using a binomial distribution *M_ij_*∼ *Bi*(*cov_ij_*, *P*_i_), where *P* represents the global mCG level of the cell. We then computed *P*(*M_ij_*> *mc_ij_*) as the hypomethylation score of cell *i* at bin *j*. The less likely to observe smaller or equal methylated basecalls, the more hypomethylated the bin is. We next binarized the hypomethylation score matrix *M* by setting the values greater than 0.95 as 1, otherwise 0, to generate a sparse binary matrix *A*. Latent semantic analysis with log term frequency was applied to *A* to compute the embedding. Specifically, we selected the columns having 1 in more than 5 rows, then computed the column sum of the matrix 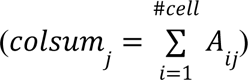 and kept only the bins with Z-scored log_2_colsum between −2 and 2. The filtered matrix was normalized by dividing the row sum of the matrix to generate *TF*, and further converted to *X* used for singular value decomposition *X* = USV^T^, where 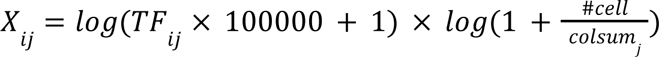. Top dimensions of *U* were then used for t-SNE visualization and cluster analysis.

We first benchmarked this method on the whole mouse brain dataset(*71*) and observed better capability to separate the annotated cell types, especially in the brain stem regions (fig. S2A). To further test the generalizability of this method, we used a PBMC dataset(*72*) where eight specific cell populations were selected by FACS and profiled with snmC-seq2. This dataset collects Naive helper T cells (CD3+, CD4+, CCR7+, CD45RA+), Memory helper T cells (CD3+, CD4+, CD45RA-), Naive cytotoxic T cells (CD3+, CD8+, CCR7+, CD45RA+), Memory cytotoxic T cells (CD3+, CD8+, CD45RA-), B cells (CD3-, CD19+), Monocytes (CD3-, CD19-, CD14+), NK cells (CD3-, CD19-, CD14-, CD16+, CD56+), and other cells (CD3-, CD19-, CD14-, CD16-, CD56-), with the FACS providing a standard for cell-type annotation. The 5kb LSI framework in ALLCools can separate the above cell types and resolve more rare cell types (fig. S2B), including CD16+ monocytes, Dendritic cells (DC), and Plasmacytoid DC (pDC). Together, these results demonstrate the accuracy and scalability of the 5kb feature-based clustering framework.

### Removal of potential doublet nuclei

We observed potential doublets in the NeuN-sorted samples of the primary motor cortex and the lung samples, shown as high-coverage cell groups between cell types in the embedding space. We developed a doublet simulation and k-NN-based method (similar to Scrublet) to remove potential doublets with 5kb mCG features. The key difference is to expand the Scrublet framework to use methylation data as input. We simulate doublet 5kb mCG profiles by adding the mc basecalls and cov basecalls of each 5kb bin of two randomly selected cells. We also required the cells to have less than 3M total reads to be included in the simulation to avoid the final dataset having too many doublets and multiplets. The hypomethylation score of the simulated doublets were computed using the summed basecalls, and LSI was used to generate embedding of the combined dataset with the observed nuclei and simulated doublets. The doublet score was computed using the same method as in Scrublet (*73*, *74*), and the threshold for excluding doublets was selected based on the score distribution of observed nuclei and simulated doublets.

### Cell embedding with chromatin contact

scHiCluster was used to generate single-cell embedding at 100 kb resolution(*75*). For each chromosome, the upper triangle within 1Mb distance of the contact matrix from each single cell was reshaped to a 1D array and single cells are concatenated vertically. Then singular value decomposition (SVD) was performed on chromatin contacts for each chromosome, with singular values divided from the matrix and only keeping the first 50 dimensions. In the next step, all chromosomes were concatenated horizontally and SVD was performed again without dividing the singular values. We then took the significant dimensions that reached the 0.01 threshold to produce a t-SNE embedding of the cells.

### Integration of data across donors

The integration was performed with a canonical correlation analysis (CCA) based method for anchor identification and data transform through these anchors(*76*). To avoid overcorrection of biological variation between cell types, we only use the anchors identified between the two donors of the same tissue to align the datasets.

For each tissue, we first identified the anchors with CCA on the LSI-transformed matrices (mCG) or concatenated SVD matrices across chromosomes (3C). Using the anchors successfully integrated cells from two donors together, but resulted in the mixing of different cell types in the integrated space (fig. S2C, left). This is due to the inaccurate identification of anchors connecting different cell types, particularly when the proportion of cell types varies significantly between donors. Therefore, we examined the distances between the two cells of each anchor in the embedding space. We found that the anchors connecting different cell types have larger distances than those connecting the same cell type, although those cells are still mutual nearest neighbors. Therefore, we only kept the top 60% of anchors with the smallest distance to eliminate anchors between different cell types, and then transformed the data. This method solves the cell-type mixture problems (fig. S2C, middle).

However, we also noticed that using the distance-based anchor filtering resulted in less integration between donors in some tissues, which means the same cell type of different donors became less mixed (fig. S2D, middle). This is due to the nature of embedding space, where an abundant cell type will explain more variation of the data and occupy larger spaces. Therefore the dominant cell type of a tissue (e.g. epithelial cells) will have a larger distance between single cells. To overcome this limitation, we used Geosketch to sample 1/3 of the cells from the dataset, so that common cell types are down-sampled and rare cell types are preserved (*77*). Next, we fit the SVD/PCA model with selected cells and transform all other cells. Then we selected the anchors based on the distance of cells in the Geosketch embedding space, and kept only the top 60% anchors. This approach allowed better integration between donors as well as less mixture of different cell types (fig. S2D, right).

### Combining the embedding of different modalities

We tested two strategies to combine the mCG and 3C embedding after integration across donors. One is to directly concatenate the embeddings, and the other is to use the weighted nearest neighbor (WNN) framework (*78*). We tested the two methods in different tissues. An example result with transverse colon data was shown in fig. S2E. WNN embedding kept the clustered cells of both mC and 3C modalities close to each other, but they were also expanded according to the variation of the other modalities and no longer formed a cluster (fig. S2E, top panels). On the contrary, the concatenated embedding kept the 3C clusters in almost their original structures but distorted the mC clusters (fig. S2E, bottom panels). In some instances, such as the colon, the mC clustering is not informative for cell types, and we would prefer the algorithm to retain more information from the 3C modality. We reasoned that WNN can quantify the contribution of different modalities if one of the modalities shows a strong separation while the other shows a weak separation. However, when the two modalities separate the cells toward different axes, the algorithm could be confused by quantifying their relative contributions. Given the necessity of dealing with this case when analyzing our data, we used the direct concatenation for joint embedding.

We used 50 dimensions from both modalities for the all cell (level 1) clustering. To decide the number of dimensions to use in the combined embedding for cell clustering within a major type (level 2), we used the Kolmogorov–Smirnov test on adjacent dimensions of the embedding, and used the smallest dimensions k where the distribution of the *k*-th dimension and *k+1*-th dimension were no longer different significantly (p>0.05). However, in some major types the test gives too small a dimension so that the clusters are not fully distinguishable. We thus manually check all the dimensions for each modality separately, and enumerate through 5, 10, 15, 20, 25, 30 dimensions, to choose the smallest number that clustering would only change little after adding five more dimensions. The full list of the number of dimensions used in major type clustering is provided in table S2.

### Initial clustering and merging into clusters and subtypes

We constructed k-nearest neighbors (kNN, k=25) graphs with the joint embedding of all cells (level 1) or each major type (level 2). Consensus Leiden clustering was performed with 500 different random initializations, with a consensus rate of 0.8 and Leiden resolution of 1.0. The initial clusters were then merged based on a supervised learning method iteratively. We train logistic regression models to classify cells from each pair of initial clusters with balanced class weights. The embedding of each modality was separately used as input to the model and performance was evaluated with average AUROC and AUPR whichever is smaller in five-fold cross-validation (separation score). Smaller cell clusters were always used as positive samples, allowing AUPR to penalize the overrepresentation of negative samples. If neither of the two modalities can separate any pair of clusters (separation score<0.9), the two clusters were merged to generate a new cluster. The clustering and merging process was performed again using the merged cluster until all clusters could be separated by at least one modality. This produced 420 clusters (denoted as c*-c*) across all cells. We also used only the mCG or only the 3C features to run the clustering and merging process within each major type, and this resulted in mC clusters (c*-mc*) and 3C clusters (c*-3c*), which were used in Fig. 6A and fig. S19. For Epi-Gas and Epi-Ent cells, the methylation embeddings reflect some global methylation-associated difference rather than variation across cell types, so we used only the 3C clusters as the final clustering result for these two major types. In total, we have 432 clusters defined across all 86,689 cells.

The clusters were further merged into subtypes according to three criteria, including tissue of origin, consistency between modalities, and association with known cell types. The clusters were first annotated if they were composed of cells primarily from one or a few tissues. This strategy is applied to the subtypes of fibroblasts, endothelial cells, perivascular cells, and immune cells, where the same major type has cells from a diverse range of tissues. Next, for each major type, we identified differentially CG-methylated regions (DMRs) and differential loops between clusters. We also processed scRNA-seq datasets from different tissues (table S3) and identified differentially expressed genes (DEGs) between each pair of annotated cell types with pyDESeq2 (*79*). We then compared the expression level of DEGs across cell types with mCG at DMRs or interaction strength at differential loops between clusters to assign the clusters to cell types. A DMR was associated with a gene if the DMR was within 10kb of the TSS, and a loop was associated with a gene if the TSS was within 2kb of the loop anchor. Cosine distances between RNA cell types and m3C clusters were computed across DMR-gene pairs or loop-gene pairs. These two steps assigned the 432 clusters into 166 annotations.

Although some clusters were assigned the same cell type annotation, their separation was supported by both mCG and 3C modalities, which suggested higher confidence in annotating them as meaningful subtypes. Therefore, we further used the iterative supervised learning framework to merge the clusters within each of the 166 annotations if only one modality can separate the pair of clusters. We train logistic regression models to classify cells from each pair of clusters with balanced class weights. The embedding of each modality was separately used as input to the model and performance was evaluated with average AUROC and AUPR whichever is smaller in five-fold cross-validation (separation score). Smaller cell clusters were always used as positive samples, allowing AUPR to penalize the overrepresentation of negative samples. If any pair of clusters could not be separated by one of the modalities (separation score<0.8), the two clusters were merged to generate a new cluster. The clustering and merging process was performed again using the merged cluster until both modalities could separate all clusters. These resulted in 206 subtypes (c*-b*) across all cells.

### Dendrogram of all subtypes

Robust dendrogram(*70*, *80*) was used to compute the dendrogram between major types and between subtypes within each major type. The joint embedding of mCG and 3C were used as input. The same dimensions were used as we performed the hierarchical clustering with all cells or within each major type. Cosine distance and average method were used to build the linkage. The dendrograms were colored based on the performance of supervised models to separate the clusters with embedding of different modalities as input features. Similar to the merging process above, each edge of the dendrogram corresponded to two scores representing the separability of cells in the two leaf nodes using mCG or 3C embedding as features. For each pairwise comparison, logistic regression and five-fold cross validation were used, and the score was assigned as the AUROC or AUPR whichever was smaller. If the mCG embedding score was >0.9 while 3C embedding score was <0.8, we colored the edge in green. If the 3C embedding score was >0.9 while mCG embedding score was <0.8, we colored the edge in orange. Otherwise, the edges were colored in black.

### Comparison with bulk methylation data

Data in the Schultz 2015 paper was downloaded from http://neomorph.salk.edu/SDEC_tissue_methylomes/processed_data/index.html in ALLC format (*24*). Genome coordinates were mapped from hg19 to hg38 using LiftOver default parameters (*81*). Data in the Loyfer 2023 paper was downloaded from GSE186458 in beta format and converted to ALLC format (*32*).

For each of these two datasets, sample-by-1kb autosomal genomic bin mCG matrices were generated with ALLCools generate-dataset. The bins covered by <=20 basecalls or having 50% base pairs overlapping ENCODE exclusion list were removed from further analysis. The subtype-by-1kb matrix was processed with the same process, and cosine distances between subtypes and bulk samples were computed across all remaining 1kb genomic bins and visualized in fig. S7.

### Identification of methylation compartments

For each 10kb bin, we computed the frequency of CpG sites with different mCG levels and used the histogram as features (100 equally sized bins between 0 and 1) to cluster all 10kb bins across the genome. We used the KMeans++ initialization and found the best results among 10 different initializations. We compared it with merging data from all major types and used the clustering result (with 36*100 features) as the initialization for each major type separately, and observed very consistent results. To compare with the chromatin compartment at 100kb resolution, we computed the proportion of 10kb bins assigned to the partially methylated compartment in each 100kb bin as PMD density score in Fig. 5E.

### Identification of DMR

mCG from + and - strands were added together before DMR calling to increase the statistical power. We identified DMRs between all major types, as well as between subtypes within each major type. Major type DMRs with >= 4 differentially methylated CpG sites (DMSs) were used. The cutoff was selected based on the maximum overlap with ATAC peaks in corresponding major types. Hypo CG-methylated DMRs were assigned to each major type. Differentially methylation calling would be affected by global mCG, and more hypo DMRs would be identified in the cell types with PMDs. These methylation differences are likely not due to cell type specificity of regulatory elements, but affected by stochasticity of methylation due to other factors. We excluded the hypo DMRs in Epi TPB, Hema Bmem, Epi-Aci, Hema Tmem, Epi AdrCtx, and Endo Vsc, where >80% hypo methylated DMRs were located in partially methylated compartments. Subtype DMRs within each major type harboring >=2 DMSs were used and merged with the hypo DMRs of that major type as the total DMRs of a major type. The comparison with ATAC peaks, chromatin loops, and motif calling were performed with total DMRs in Figs. 2 and 5H, and figs. S10 and 11.

### Preprocessing of snATAC data

Fragment files of snATAC data were downloaded at http://catlas.org/catlas_downloads/humantissues/fragment/. Only the samples that overlap with m3C profiled samples were used. Integration of snATAC data and mCG data were performed with ALLCools using 5kb bin features.

We fit the LSI model with mCG data *Bm* to derive *Um*. The intermediate matrices *S* and *V* and vector *IDF* were used to transform the ATAC data *B*α to *U*α, by

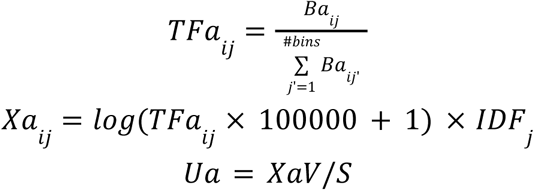

CCA was also performed with the downsampling framework using 100,000 cells from each dataset as a reference and the others as query, but taking the TF-IDF transformed matrices as input. The query cells were projected to the same space using the IDF and CCV of the reference cells. Specifically, *Bm_ref_* and *B*α*_ref_* were converted to *Xm_ref_* and *X*α*_ref_* with TF-IDF, and the CCVs (denoted as *U_ref_* and V*_ref_*) were computed by 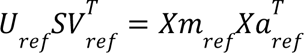. Then *Bm_qry_* and *B*α*_qry_* were converted to *Xm_qry_* and *X*α with TF-IDF using the IDF of reference cells, and the CCVs (denoted as U*_qry_* and V*_qry_*) were computed by 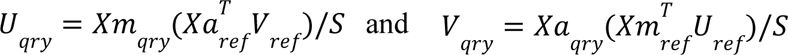. The following steps to find anchors and align *Um* and *U*α were the same as originally implemented in our previous works (*71*, *82*).

Based on the integration results, we match the cell types labeled in the snATAC-seq data to the major types of our m3C annotations (table S4). The celltype-by-peak matrix and peak genome coordinates were downloaded at http://catlas.org/catlas_downloads/humantissues/cCRE_by_cell_type/ and peaks of the celltype assigned to each major type were merged as the peak of the m3C major type. The proportion of peaks within 1kb of a DMR of the same major type were reported in the middle of the Venn plots in Fig. 2K and dark green in fig. S11A, as well as the proportion of DMRs within 1kb of a peak in the same major type. The majority peaks were also used in Fig. 3D to quantify the mCH level at cCREs. The distal peaks (>2kb from TSS) for each cell type were used as cell type distal cCREs, and other peaks that have been assigned to any other cell types were used as other distal cCREs.

### Motif enrichment analysis

Three methods were used for motif enrichment analyses in different situations. PycisTarget (*83*) is very efficient for multiple group comparisons, by creating a database first as a background and performing the tests for all different groups by extracting values from the database. It also has the biggest motif collections (*83*) as the candidate known motifs for the tests. Therefore the motif analyses between DNA groups were performed using pycisTarget. We first build the databases of motif rankings and scores using all DMRs (Fig. 2M), total DMRs of each major type (Fig. 5H), or all DMRs associated with differential loops (Fig. 6G). Then pycisTarget was used to test motif enrichment against the SCENIC+ motif collection (*83*), and normalized enrichment score (NES) >3 was used as the cutoff for enriched motifs in each region group. For visualization of TFs in different major types (Fig. 2M), we randomly select a TF associated with a motif if multiple TFs are mapped to the same motif ID in the pycisTarget database. For the TFs in skeletal muscle (Fig. 6G), we use pseudo bulk RNA expression of skeletal muscle from Micheli et al. 2020(*84*) to rank the TFs and choose the highest expressed TF associated with the motif.

The pycisTarget motif collection contains big motif clusters where a single motif ID matched hundreds to thousands of genes, the TF methylation preference analyses could be confused since the TF genes mapped to the same motif includes both methylPlus and methylMinus(*85*) ones. Therefore, we used AME(*86*) with HOCOMOCO motifs(*87*) for the analysis in fig. S12, D and E (*88*, *89*). 1kb regions from the DMR centers were used and hypo methylated peaks (mCG < 0.1) and hyper methylated peaks (mCG > global mCG + 0.3) were compared against each other.

The pycisTarget motif collection also contains many TF to motif pairs where the TF themself do not have a specific motif but their ChIP-seq shows motif preferences due to cofactors’ binding (e.g. cohesin). To infer the TFs implicated in loop formation, we used Homer (*90*) and its own motif database to perform the enrichment analyses. The DMRs that overlap the anchor of any loop summits by >=1bp (loop DMRs) were used as a foreground, and those that do not were used as a background.

All the motif enrichment analysis used a fixed region size, where we uniformed the foreground and background regions to be ±500bp from the center of the regions before motif scanning. If the significant motifs are more consistent between similar cell types and between methods, we are more likely to trust the result. This is another reason we showed results from HOMER rather than pycisTarget for the loop DMR analysis.

### Repeat-based single-cell clustering

To better assess the repeat-based single-cell clustering without confoundment from the abundance of population of cell types, we downsampled the dataset so that each type has at most 800 cells. The coordinates of repeat were extracted from RepeatMasker (https://www.repeatmasker.org/) labeled hg38 genome. The cell-by-feature matrices were generated for each repeat class (LINE, SINE, LTR, etc) by quantifying mCG fraction at repeat regions. The matrices were further binarized with a threshold of 0.5 and LSI was performed for dimension reduction. t-SNE was applied on the dimension-reduced matrices for visualization. We performed Leiden clustering with a resolution of 2.0 on the dimension-reduced matrices. The clustering results were then compared to the major type clustering using Adjusted Rand Index (ARI) to assess the quality.

### Methylation ratio calculation and ratio-based cell embedding

This method is used to quantify the mCH ratio of genomic bins or genes within each cell (Fig. 3, F and G, and figs. S17 and S18A) or each major type (Fig. 3, C and H, and figs. S15, 16, and S18B), and the mCG ratio of genomic bins and DMRs across subtypes (figs. S6, 7, and S10B) and major types. We use *mc_ij_* and *cov*_*ij*_ to denote the total cytosine basecalls in a specific sequence content in cell *i* and feature *j*. For each cell, we calculated the mean (*m*) and variance (*v*) of the mCH (or mCG) level across the features. A beta distribution was then fit for each cell *i*, where the parameters were estimated by

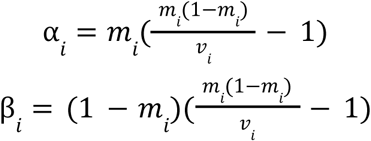

We next calculated the posterior mC level of each bin *j* by

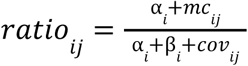

We normalized this rate by the cell’s global mean methylation by

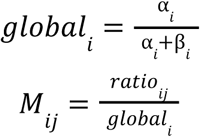

The values greater than 10 in *M* were set to 10. After normalization, *M*_*ij*_ is close to 1 when *cov*_*ij*_ is close to 0.

For cell embedding using mCH, PCA was performed on *M* to generate embeddings, and corrected for donor differences using the integration method above. mCG anchors between donors were used except in Epi Gas, Epi Ent, and Epi TPB, where chromatin contact anchors were used. The corrected embeddings were used for t-SNE visualization.

### Chromatin compartment analysis

Pseudo-bulk contact matrices were generated at 100 kb resolution for each chromosome and used for compartment analysis. To establish a reference, we merged the contact maps from all 86,689 cells and performed principal component analysis (PCA) on these matrices. Before PCA, we filtered out 100 kb bins with abnormal coverage. Coverage for bin *i* on chromosome *c* (denoted as R_c,i_) was defined as the sum of the *i*-th row in the contact matrix for chromosome *c*. We retained bins with coverage between the 99th percentile and twice the median minus the 99th percentile of R_c_. Contact matrices were first distance-normalized, followed by the computation of Pearson correlation matrices. PCA was then applied to the correlation matrices per chromosome, and the first principal component (PC1) was used as the compartment score. The sign of PC1 was adjusted such that regions with higher CpG density had positive scores. We confirmed that PC1 corresponded to the expected plaid pattern of the correlation matrix rather than simply distinguishing chromosome arms.

For each major cell type, contact maps were processed in the same way and projected onto the PCA model derived from the merged dataset. Raw contact matrices were used for PCA fitting, while both raw and imputed matrices were used to compute compartment scores for each cell type—referred to as raw and imputed compartment scores, respectively. Imputed matrices performed better for smaller cell populations, whereas raw matrices offered higher resolution when large numbers of cells were available.

To evaluate whether long-range interactions occurred within or between compartments, we stratified contact distances based on either the difference or the sum of compartment scores at the interaction anchors. A larger difference indicated inter-compartment interactions, while the sum of scores distinguished between AA (positive sum) and BB (negative sum) intra-compartment interactions. Although long-range interactions generally included more inter-compartment contacts, we observed that non-neuronal cells—despite having more long-range interactions—had a higher *proportion* of intra-compartment contacts at long distances compared to neurons, based on distance-normalized contact counts (as shown in fig. S19A). These results reflect relative proportions rather than absolute counts of intra-compartment interactions.

Compartment strengths are computed in the same way as described in Nora et al. 2017 (*91*). Specifically, within each chromosome, we rank all the 100kb bins based on compartment scores, and group the bins into 50 equal-interval groups. The distance normalized interaction strength between each pair of bins were averaged within each group. The axes are ranked by the compartment score of the cell types so that BB interactions are on the top left and AA interactions are on the bottom right.

Compartment strength was calculated following the approach described in Zhang et al., 2019 (*91*, *92*). For each chromosome, 100 kb bins were ranked according to their compartment scores and divided into 50 equally sized groups. Distance-normalized interaction frequencies were then averaged for all bin pairs within and between these groups. To generate all chromosomes-averaged saddle plots, bins associated with B-type compartments (lower compartment scores) were positioned in the upper-left corner of the matrix, while A-type bins (higher scores) were placed in the lower-right. This arranged matrix (denoted as S) ensured that BB interactions appeared in the top-left quadrant and AA interactions in the bottom-right. Then the compartment strength for a cell type was quantified as:

Compartment strength = [Sum(S_1:10,1:10_) + Sum(S_41:50,41:50_)] / [Sum(S_1:10,41:50_) + Sum(S_41:50,1:10_)] This ratio reflects the relative enrichment of same-compartment (AA and BB) versus cross-compartment (AB and BA) interactions, providing a robust measure of compartmentalization.

To identify compartment boundaries at higher resolution, we annotated a subset of domain boundaries as compartment transitions, based on the observation that domain boundaries often coincide with compartment switches. In Fig. 5, F and G, and fig. S21, C and D, we assessed the compartment scores across a 1 Mb window centered on each domain boundary (500 kb on either side). Boundaries were defined as A-to-B or B-to-A if the upstream and downstream compartment scores were separable with an area under the ROC curve (AUROC) > 0.9 or < 0.1, respectively.

### Domain analysis

Raw and imputed contact maps at 25kb resolution were merged into pseudobulk. We used the wrapper of TopDom (*93*) in scHiCluster (*75*) to call domains for all major types.

### Loop and differential loop analysis

Chromatin loops were identified with scHiCluster (*18*, *75*) in each major type, subtype, and cluster, respectively. We only perform loop calling between 50 kb and 5 Mb, given that increasing the distance only leads to a limited increase in the number of significant loops. For each single cell, the imputed matrix of each chromosome Q_cell_ was log-transformed, and Z-score normalized at each diagonal (result denoted as E_cell_) and subtracted a local background between >=30 kb and <=50 kb (result denoted as T_cell_). A pseudo-bulk level t-statistic was computed to quantify the deviation of E and T from 0 across single cells from the cell group, where larger deviations represent higher enrichment against global (E) or local (T) background. E_cell_ is also shuffled across each diagonal to generate E_shufflecell_, and then T_shufflecell_, to estimate a background of the t-statistics. An empirical FDR can be derived by comparing the t-statistics of observed cells versus shuffled cells. We required the pixels to have an average E >0, fold change >1.33 against donut and bottom left backgrounds, fold change >1.2 against horizontal and vertical backgrounds, and FDR <0.01 compared to global (E) and local (T) backgrounds. The loop summits were selected from the loop pixels with a breadth-first search algorithm, where we started from the loop pixels with the largest E, and connected it with all the other loop pixels within 20kb (L0 distance) with smaller E values. The loop pixel with the largest E value in each connected component of loop pixels was defined as a loop summit. We only used the concept of summit during the counting of loop summits, and in all other cases, we used “loop” to represent loop pixels.

To compare the interaction strength of loops between different groups of cells, we adopt an analysis of variance (ANOVA) framework to compute the F statistics for each loop identified in at least one cell group using either Q_cell_ (result denoted as F_Q_) or T_cell_ (result denoted as F_T_). P-values were computed based on the ANOVA statistics, and FDRs were estimated by Benjamini-Hochberg procedure across all loops. Then, we log transformed and Z-scored F_Q_ and F_T_ across all the loops being tested and selected the ones with converted F_Q_ and F_T_ > 1.036 (85th percentile of standard normal distribution) and FDR<0.05 as differential loops.

Aggregate peak analysis (APA) of some of the comparisons is shown in fig. S20D. For each single loop pixel, the imputed contact map from −100kb to +100kb was selected and min-max normalized to the range of 0 to 1, and averaged across all the differential loops that have a folder change of Q and T greater than 1.2 and 1.5, respectively, comparing the average of foreground cell types and the average of background cell types. Z-score at the center pixel was computed, using the mean and standard deviation of the 5×5 matrix at the bottom left corner of the whole 21×21 matrix.

### Data storage and algorithm implementation

Efficient data storage and computation is essential in this project to deal with the large scale datasets. Several engineering strategies were used to resolve the challenges. The methylation analysis is based on ALLCools, which we developed to generate ALLC files, compute methylation ratio, perform cell embedding with ratio based or LSI based method, and call methylation compartment and DMR. Single-cell level methylation basecalls for each cytosine were saved in ALLC files in tsv format. The cell-by-bin or cell-by-gene matrices served as a starting point for clustering and downstream analysis, and were saved in a single methylcytosine dataset (MCDS) file. MCDS format represents a collection of labeled multi-dimensional arrays. The in-memory representation of MCDS builds on the Dataset class in the Xarray package(*94*), which associates raw multi-dimensional data arrays with labeled dimensions, coordinates and attributes. This labeling system is a key feature to allow accessing the data consistently and associating them with the cell or gene metadata. The on-disk serialization of MCDS and RegionDS builds on the zarr package(*95*), which is a storage format for chunked, compressed, multi-dimensional arrays allowing efficient access of data without loading the whole dataset into memory.

The 3D genome analysis is based on scHiCluster, which we developed to achieve raw contact imputation, and compartment, domain, and loop calling, as well as differential loop analyses. The raw contacts were saved in tsv format. The imputed cool files at single cell and pseudobulk level were saved in cooler format. The intermediate files were saved in npz format and deleted when all the downstream files had been generated. The imputation and loop calling were wrapped into a Snakemake pipeline for efficient parallelization(*96*). The imputation was parallelized between cells and chromosomes, and took 30 minutes at 100kb resolution, 2.5 hours at 25kb resolution, and 5 hours at 10kb resolution for 1536 cells with 96 cpus at anvil.rcac.purdue.edu(*97*).

## Acknowledgements

We thank Rajesh Ilango, Scott Newins, Nicholas D. Youngblut, and Dave P. Burke for helping embed the web application into Arc domain names. We thank Brian Plosky and Joseph Caputo for assistance with manuscript preparation. We thank Ruochi Zhang and Siyu He for insightful discussions.

## Author contributions

Conceptualization: JZ, CL, JRD, JRE

Data Production: RGC, AB, JRN, BC, SM, ML, NC, LB, CO

Data Analysis: JZ, YW, HL, WT, DS Visualization Tool: ZZ, GY, JZ, SC

Sample Source: CB, RAeD, AKW, MS, JC-K, SZ, MPS, SP, BR

Writing – original draft: JRD, JZ, YW Writing – review & editing: All authors

## Data and materials availability

Single-cell DNA methylation data generated in this study have been deposited to in https://huggingface.co/datasets/zhoujt1994/HumanCellEpigenomeAtlas_sc_allc/tree/main in tsv format. Single-cell chromatin contact data generated in this study have been deposited to https://huggingface.co/datasets/zhoujt1994/HumanCellEpigenomeAtlas_sc_contact/tree/main in tsv format. Interactive browser and download links for more processed files can be found at https://humancellepigenomeatlas.arcinstitute.org.

---

**Fig. S1.**
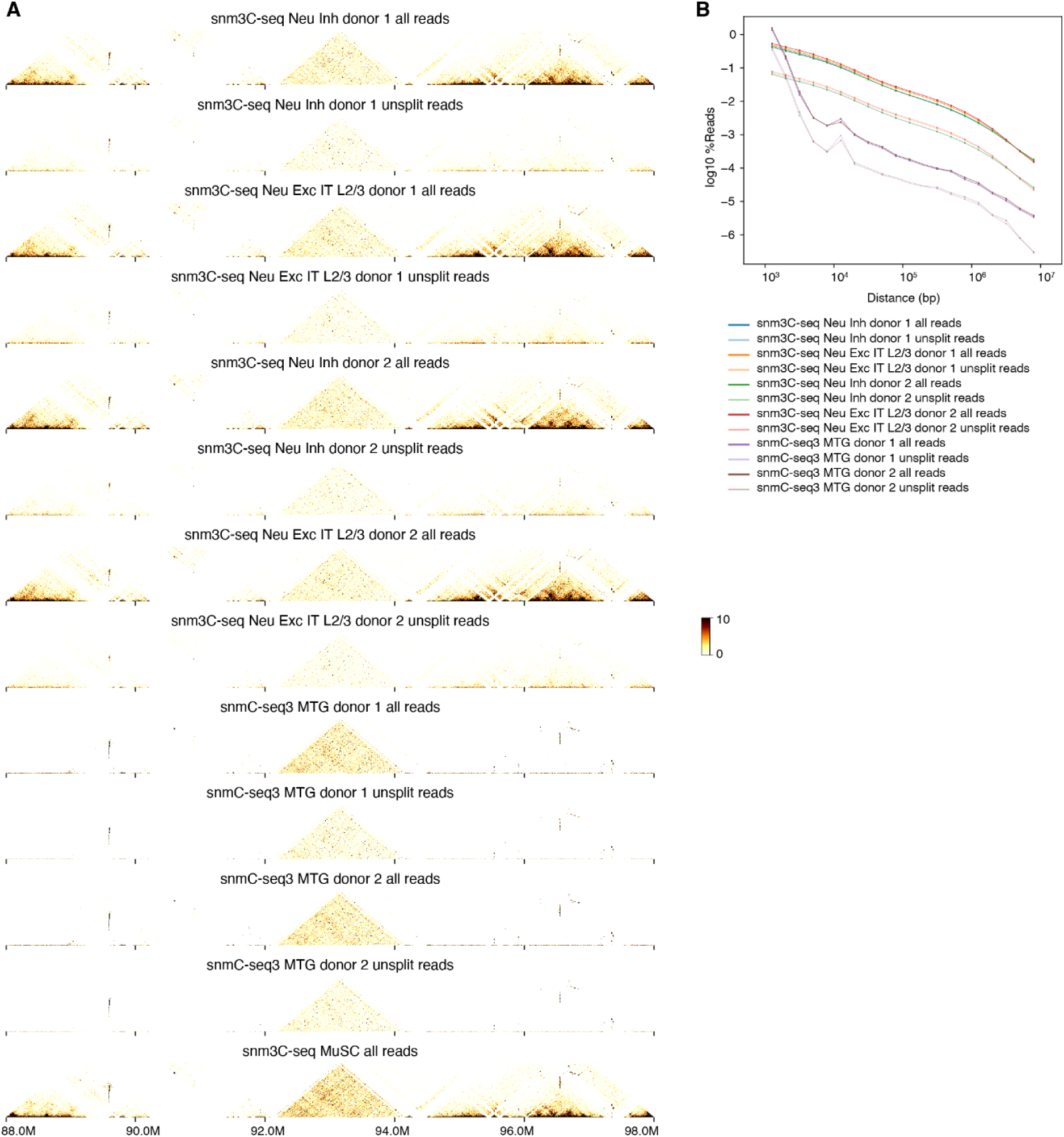
Chromatin contact quality control. (**A**) Contact map of single-nucleus methyl-3C sequencing (snm3C-seq) or single-nucleus methyl-C seq3 (snmC-seq3) from different cell types/tissues (L2/3 excitatory neurons (Neu Exc IT L2/3), inhibitory neurons (Neu Inh), bulk middle temporal gyrus (MTG), muscle stem cells (MuSC)), donors, or mapping methods (with/without read split). (**B**) Chromatin contact decay plots from snm3C-seq or snmC-seq3 datasets.

**Fig. S2.**
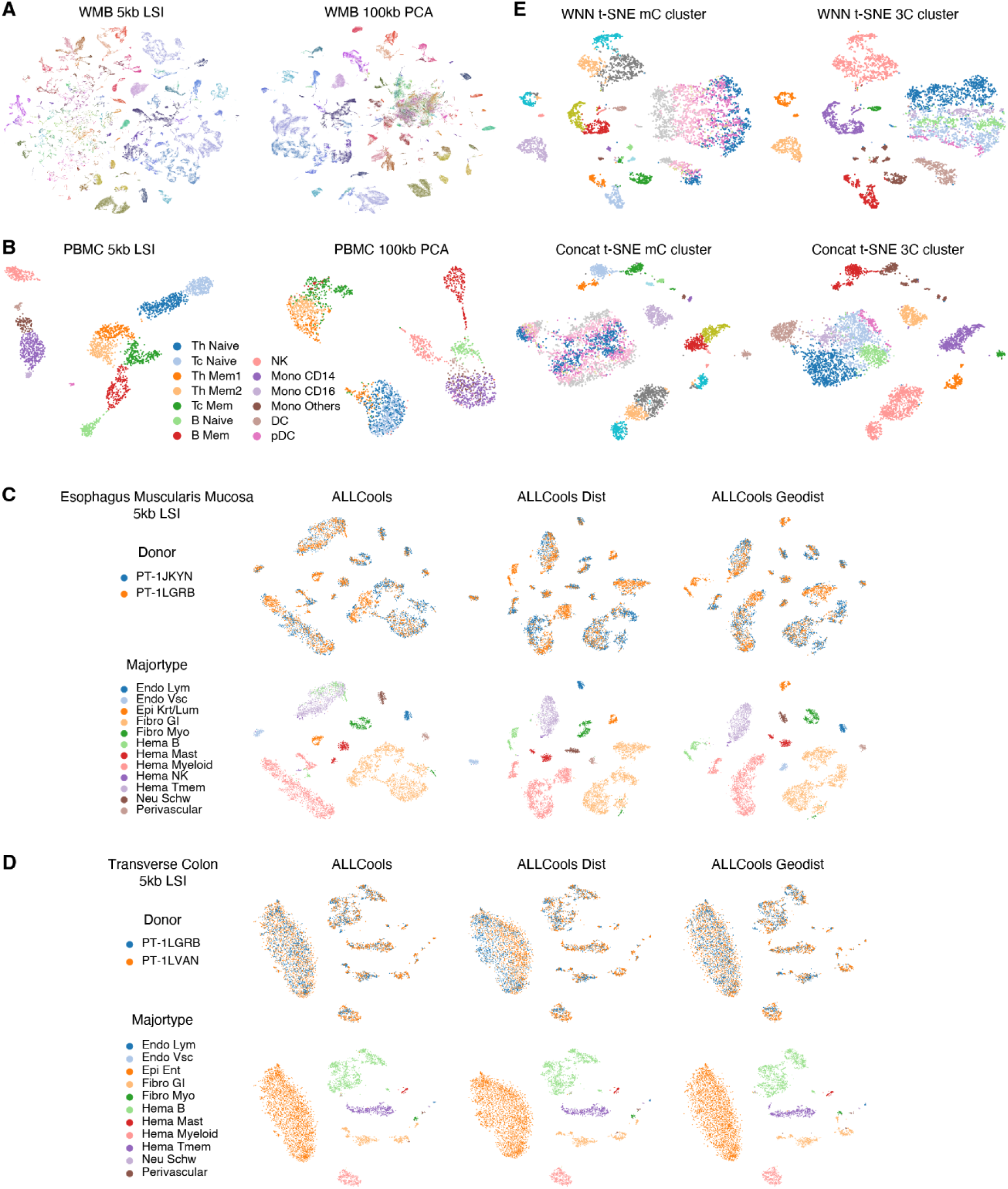
Improved methods for clustering and batch effect correction in snm3C-seq data. (**A,B**) t-SNE of whole mouse brain cells (A) and peripheral blood mononuclear cells (PBMCs; B) using Latent Semantic Indexing (LSI) based on 5kb bins (left) compared with previous approaches using Principal Component Analysis (PCA) based on 100kb bins (right) colored by cell types. (**C,D**) t-SNE of esophagus (C) or colon (D) cells using 5kb mCG with different anchor filtering methods for batch effect correction colored by donors (top) or cell types (bottom). (**E**) t-SNE of colon cells using weighted nearest neighbors (top) or concatenated PCs (bottom) to combine 5kb mCG and 100kb chromatin contact embeddings colored by mCG clusters (left) or chromatin contact clusters (right).

**Fig. S3.**
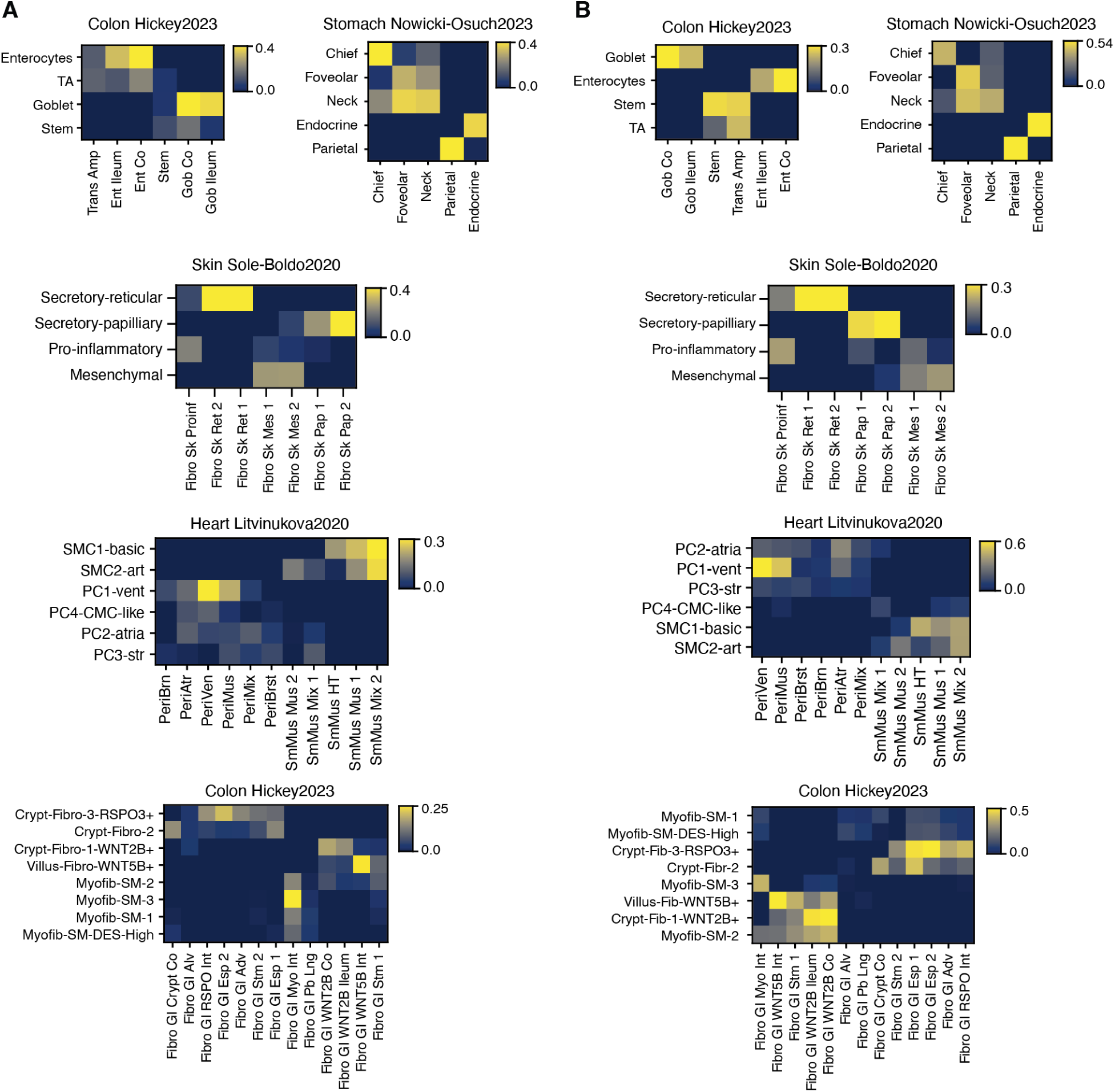
Cell type comparison with RNA-seq. (**A**,**B**) For each pair of subtype (row) and cell type labeled in scRNA-seq data (column), the cosine similarity between expression of DEGs and strength of differential loops (A) or mCG at DMRs (B) across DEG-loop pairs or DMG-DMR pairs.

**Fig. S4.**
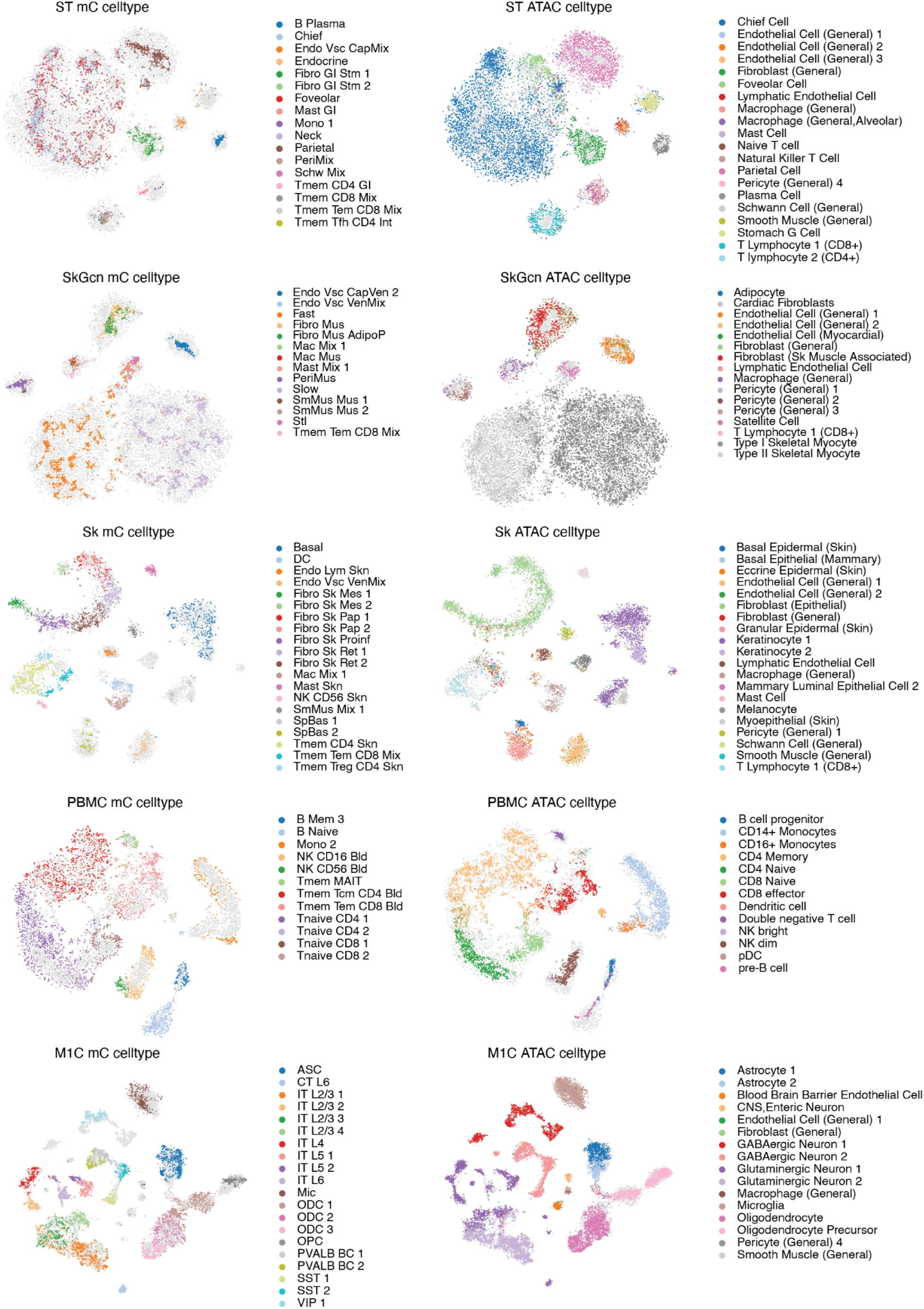
Integration of DNA methylation and ATAC-seq. T-SNE of cells from example tissues using joint embedding of snATAC-seq cells and snm3C-seq cells with 5kb open chromatin and mCG, colored by subtype annotations in our study (left) or the ATAC study (right) with the other modality in grey. A subtype is shown only if it has >=30 cells and is among the 20 most abundant subtypes in each tissue.

**Fig. S5.**
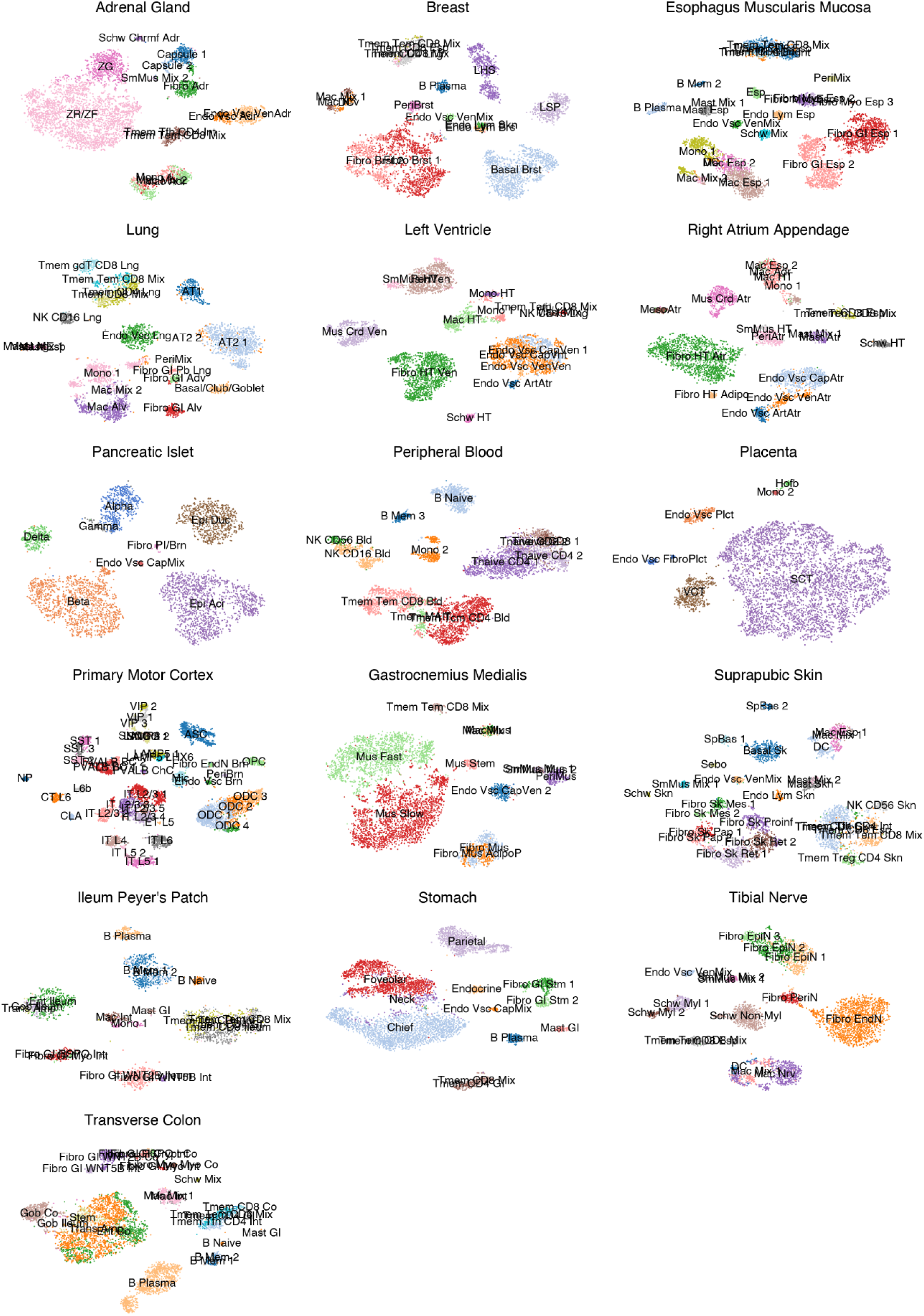
Cell types by tissue. T-SNE of cells from each of the 16 tissues using joint embedding of mCG and chromatin contacts colored by subtypes. Only subtypes with >=30 cells in each tissue were shown.

**Fig. S6.**
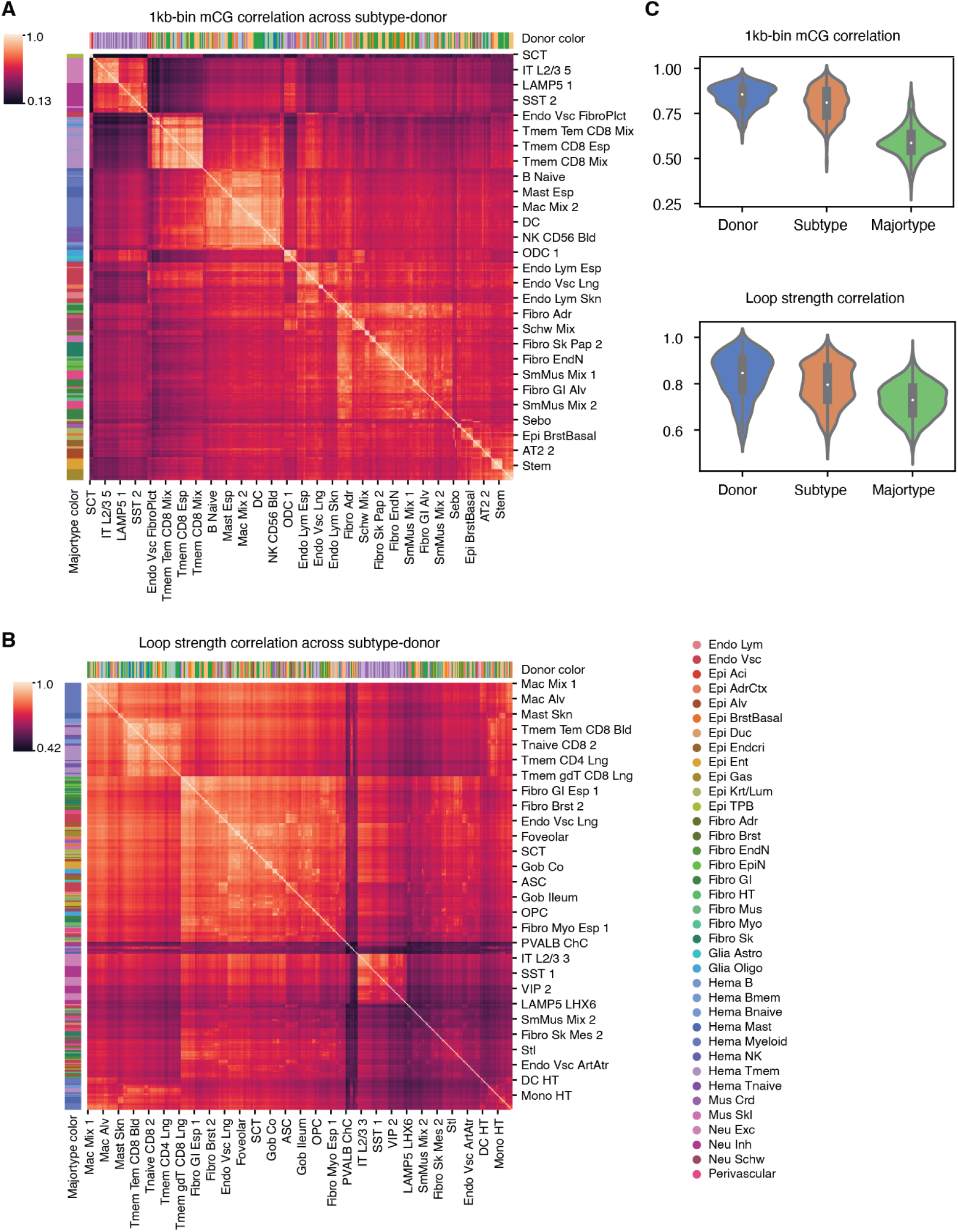
Correlation between cell types and donors. (**A,B**) Pairwise correlations of mCG (A) or loop strength (B) between 715 subtypes-donors with >=30 cells across 1kb bins (A) or all loops identified in major types (B). (**C**) Correlations of mCG (top) or loop strength (bottom) comparing the same subtype between donors (blue), between cell subtypes in the same major type and donor (orange), or between major types within the same donor (green).

**Fig. S7.**
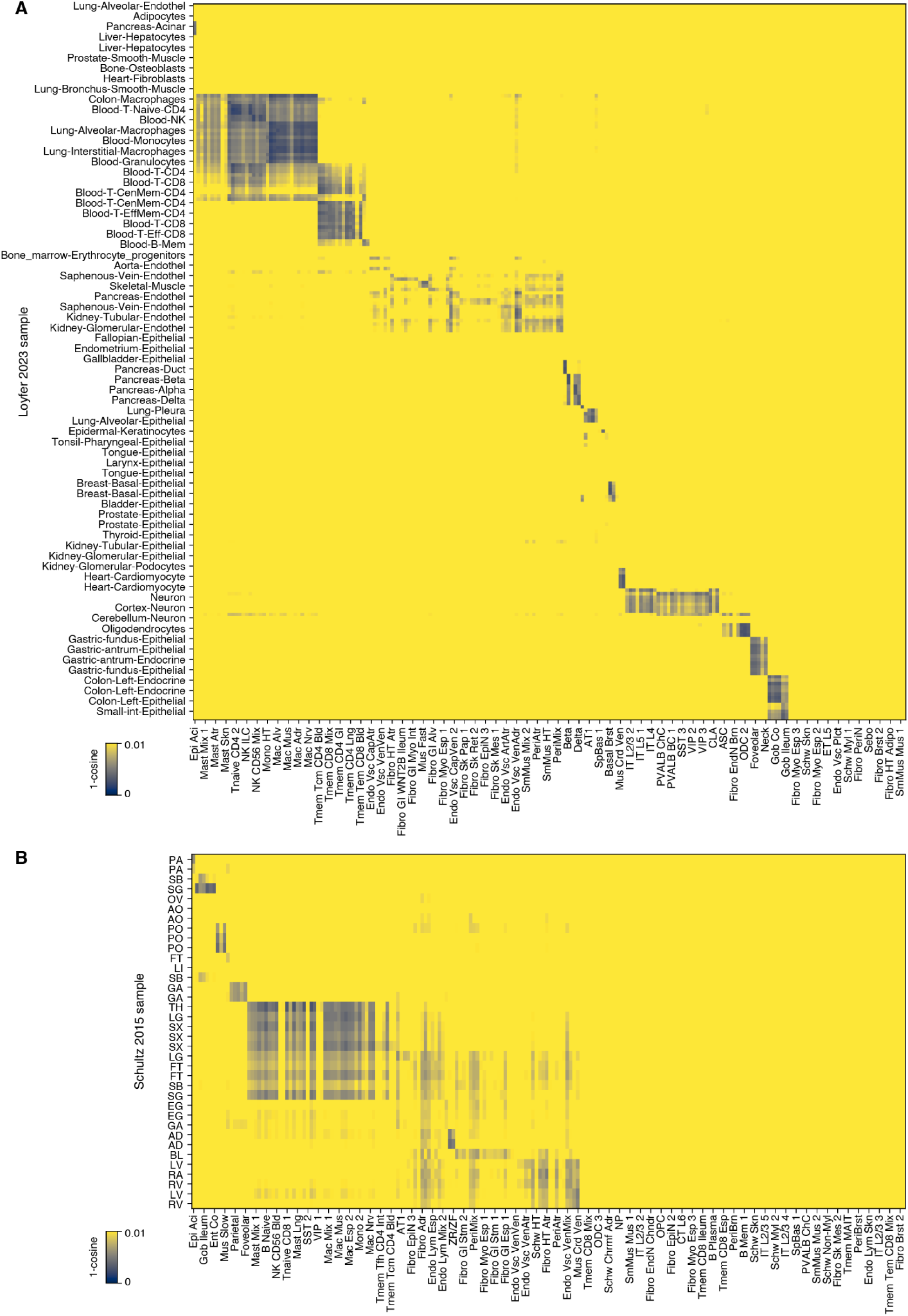
Correlation between snm3C-seq DNA methylation and prior studies. (**A,B**) Correlation of mCG between cell subtypes identified in this study (x-axis) with cell types from prior studies of cells sorted from human tissues (y-axis; A) or from bulk human tissues (y-axis; B) across 1kb bins.

**Fig. S8.**
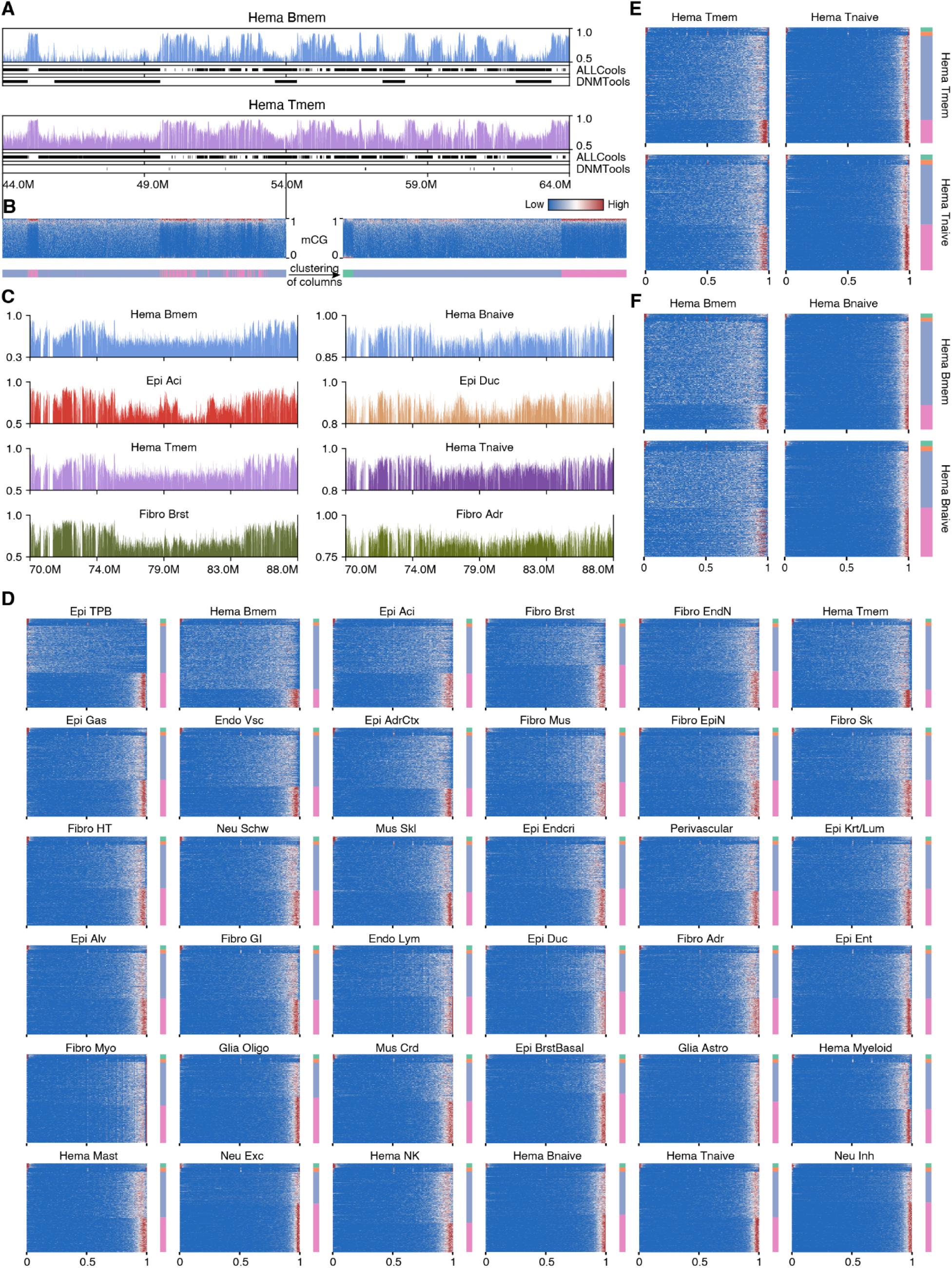
Partially methylated CpG compartments across cell types. (**A**) mCG and calls for partially methylated domains (PMDs) based on methylation compartment calling as part of this study (ALLCools) or DNMTools in two cell types at chr14:44,000,000-64,000,000. (**B**) Distribution of mCG in Hema Tmem over CpGs for 10kb bins (columns of top heatmap) and methylation compartment (bottom) at chr14:44,000,000-54,000,000 ordered by genome coordinate (matching regions in (A); left) or methylation compartment (right). (**C**) Browser views of mCG at chr2:70,000,000-88,000,000 (the same region as Fig. 2C) but different y-axis ranges. (**D**) Distribution of mCG over CpGs for 10kb bins in all major cell types (left) and methylation compartment colorbar (right) ordered by the methylation compartments. (**E,F**) Distribution of mCG in memory (left) or naive (right) cells over CpGs for 10kb bins (row) ordered by methylation compartments called in memory (top) or naive (bottom) cells from T cell (E) or B cell (F) lineage.

**Fig. S9.**
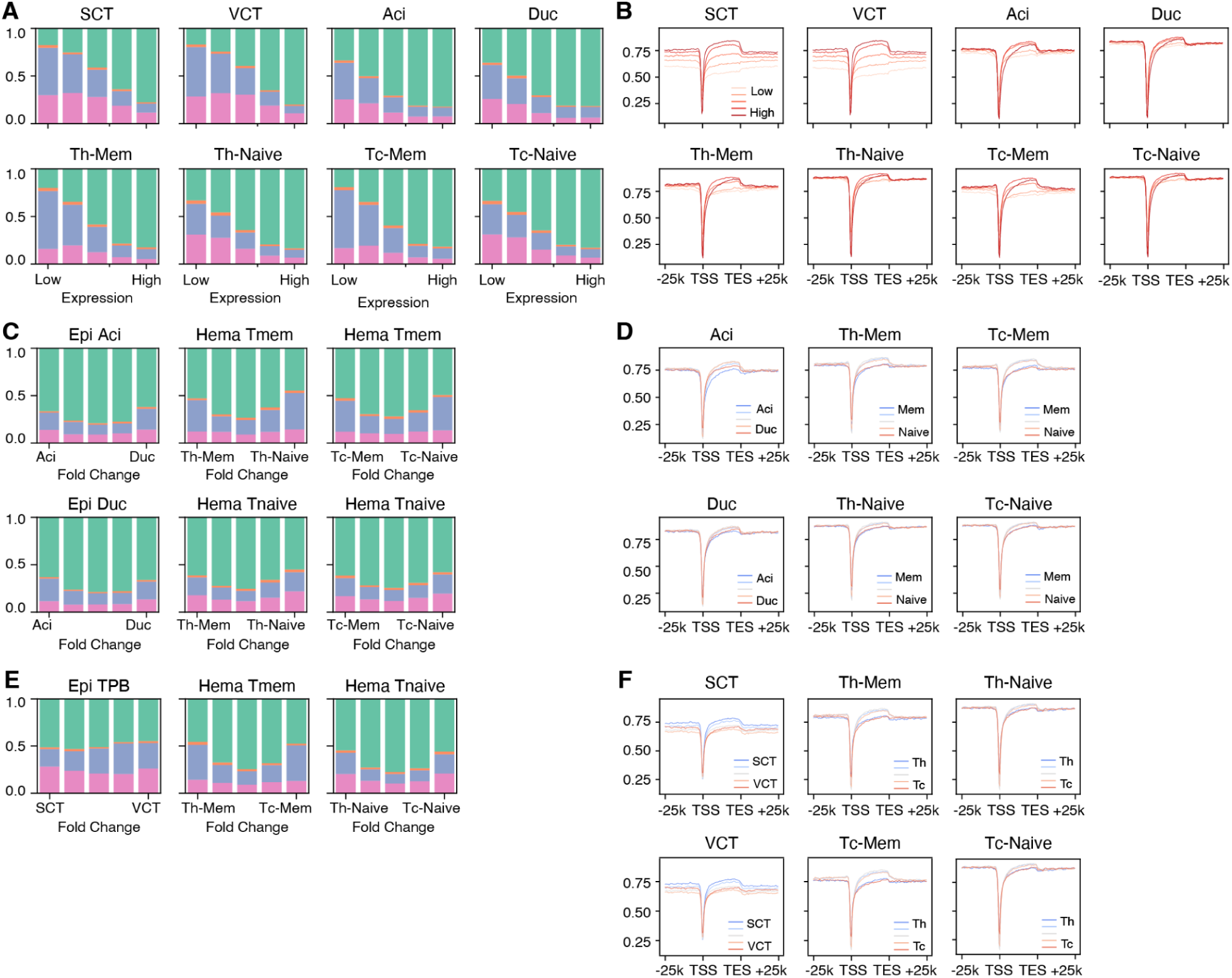
CpG methylation compartments across cell types. (**A-F**) Fraction of genes associated with each methylation compartment (A,C,E) or gene body mCG (B,D,F) stratified by gene expression level (A,B), fold-change between non-PMD cell type and PMD cell type (C,D), or fold-change between subtypes of PMD cell type or non-PMD cell types (E,F).

**Fig. S10.**
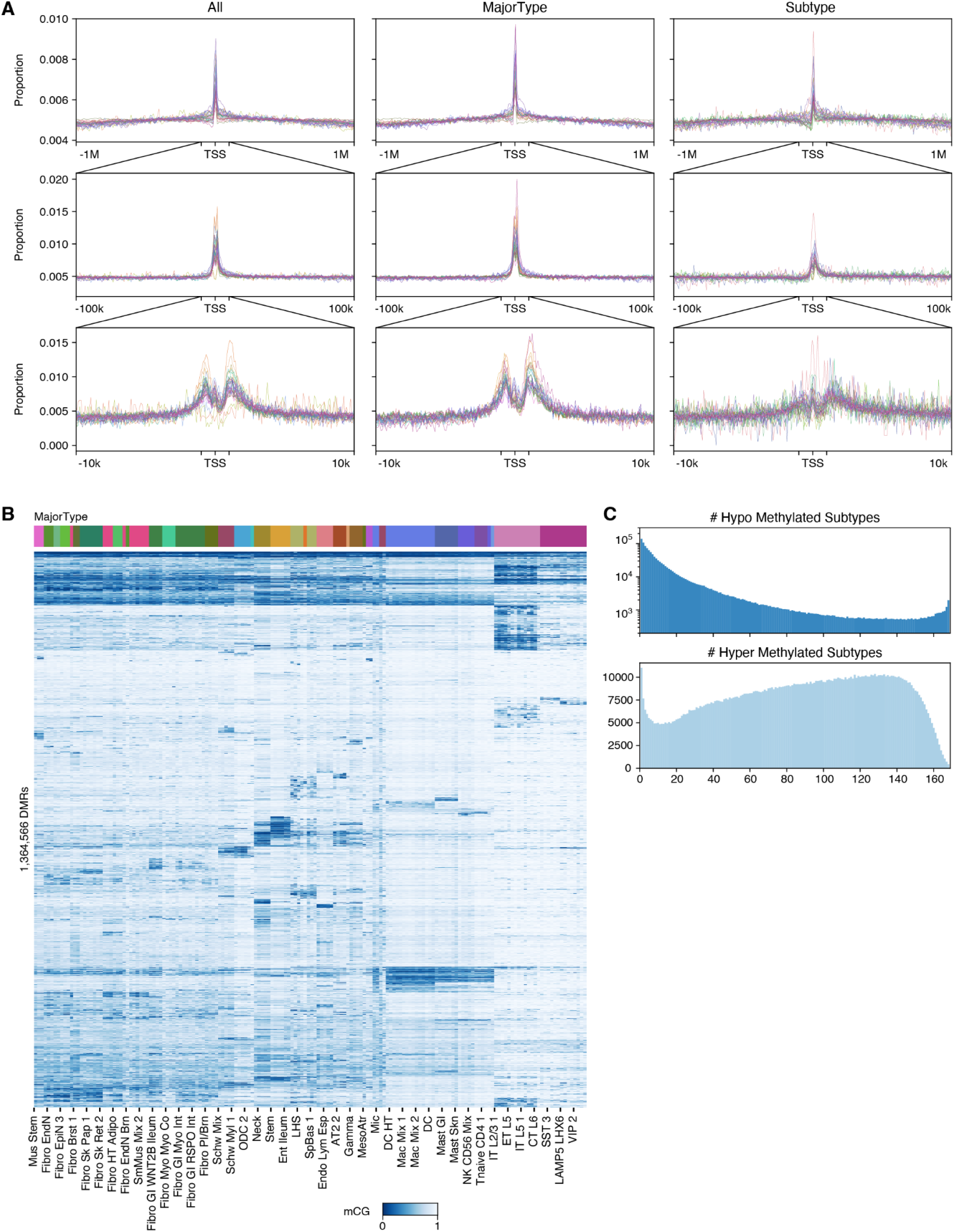
Summary of DMRs. (**A**) Proportion of total DMRs, major type DMRs, and subtype DMRs (Methods) at different distances to TSS. (**B**) mCG of DMRs across 168 subtypes (PMD cell types excluded). Column colors of corresponding major types. (**C**) Proportion of DMRs that are hypo (top) or hyper (bottom) methylated in different numbers of subtypes.

**Fig. S11.**
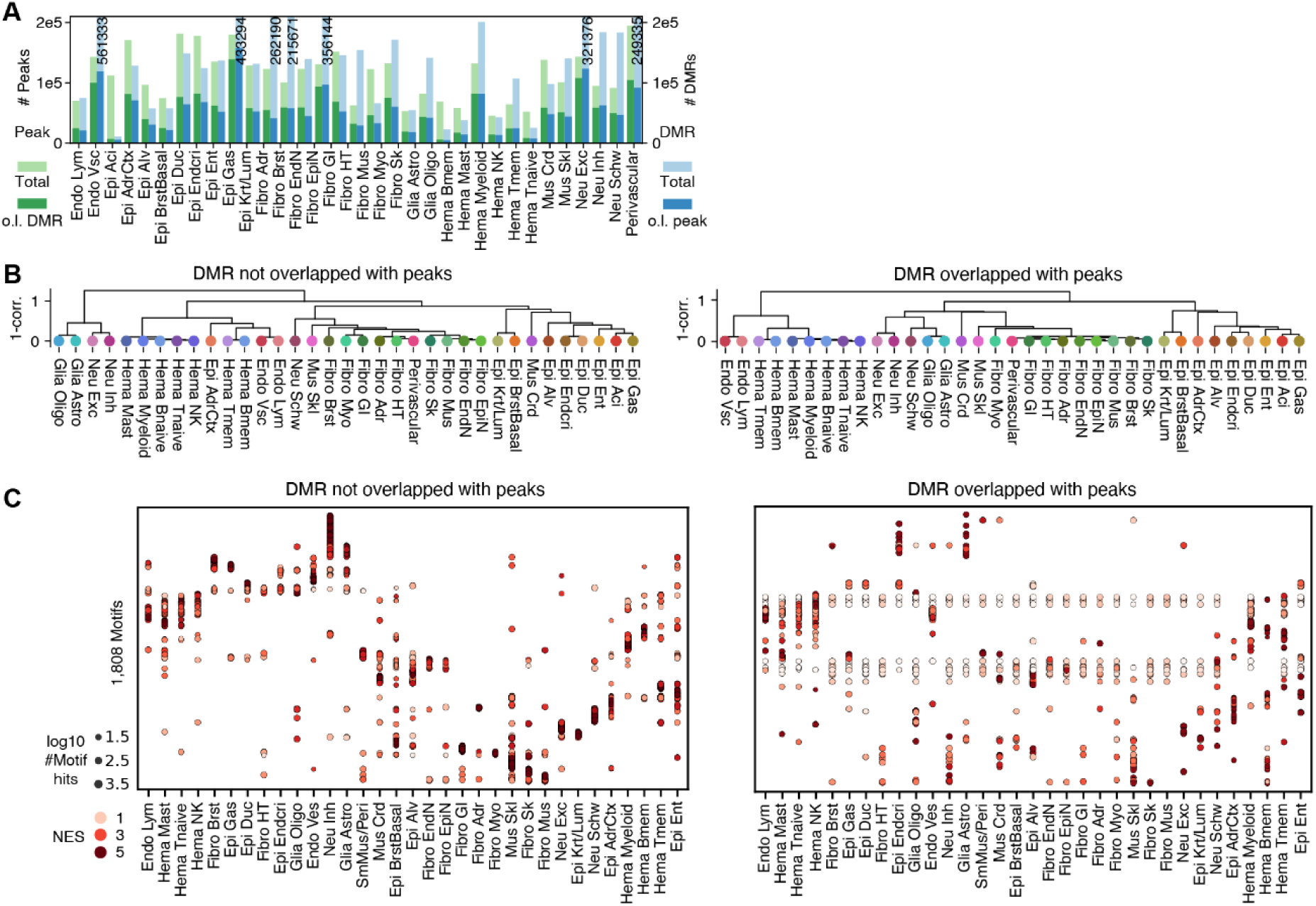
Overlapping between DMRs and ATAC-seq peaks. (**A**) Bar plots showing the overlap between DMRs and ATAC-seq peaks for each major cell type. (**B**) Dendrogram of major types using mCG at non-peak DMRs (left) or peak-DMRs. (**C**) Motif enrichment in non-peak DMRs (left) or peak DMRs (right) of each major type. Only motifs enriched in >=1 major types in non-peak DMRs are shown. Row and columns are ordered by normalized enrichment scores (NESs) from pycisTarget in non-peak DMRs. NESs are row-wise Z-scored.

**Fig. S12.**
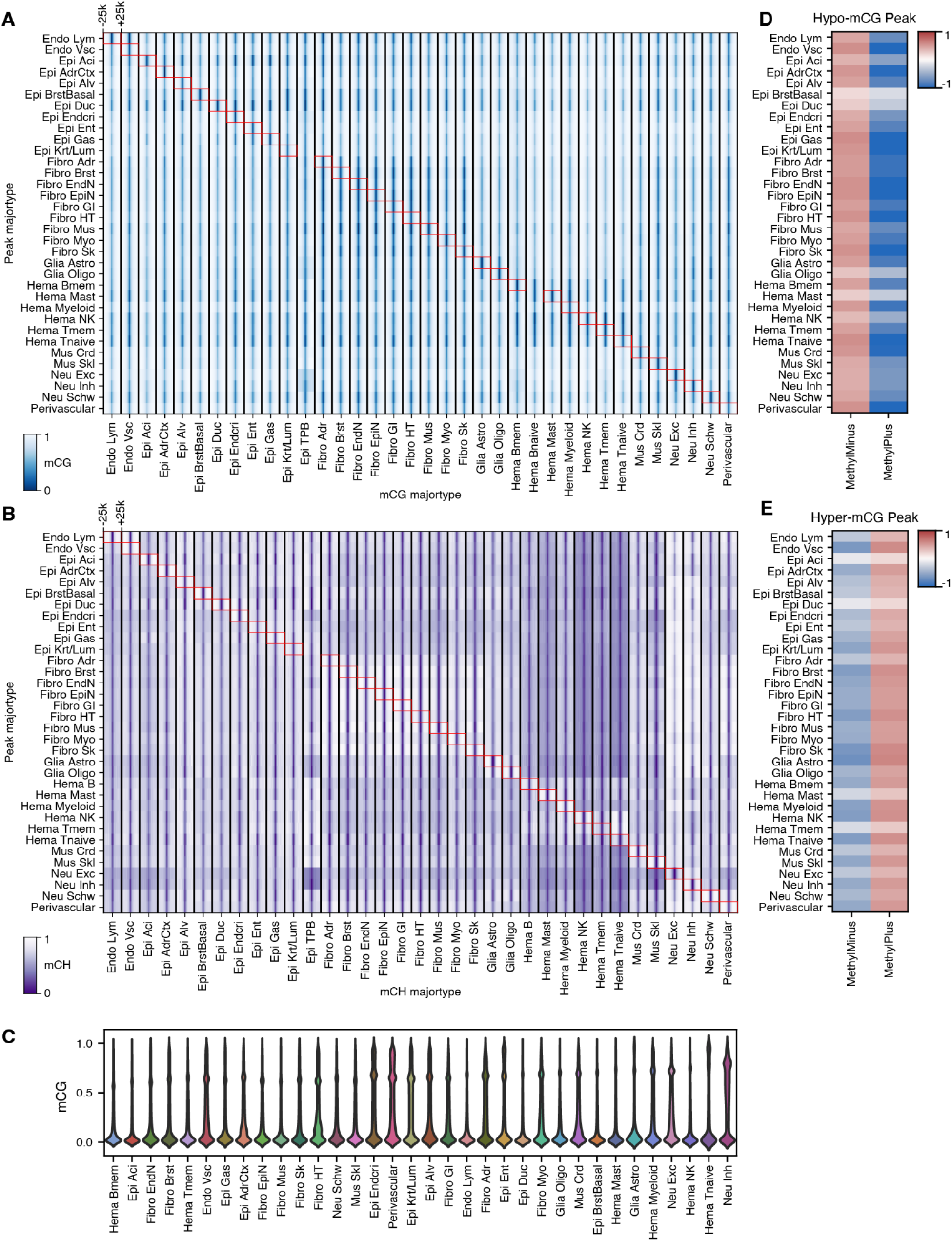
DNA methylation diversity at ATAC-seq peaks. (**A**,**B**) mCG (A) or mCH (B) in y-axis cell types at ±25kb of ATAC peaks called in x-axis major type. Each row is the average over peaks. (**C**) Distribution of mCG across ATAC peaks in each major type. (**D**,**E**) Odds ratio between proportion of methyl plus(minus) TFs among enriched TFs and among all tested TFs.

**Fig. S13.**
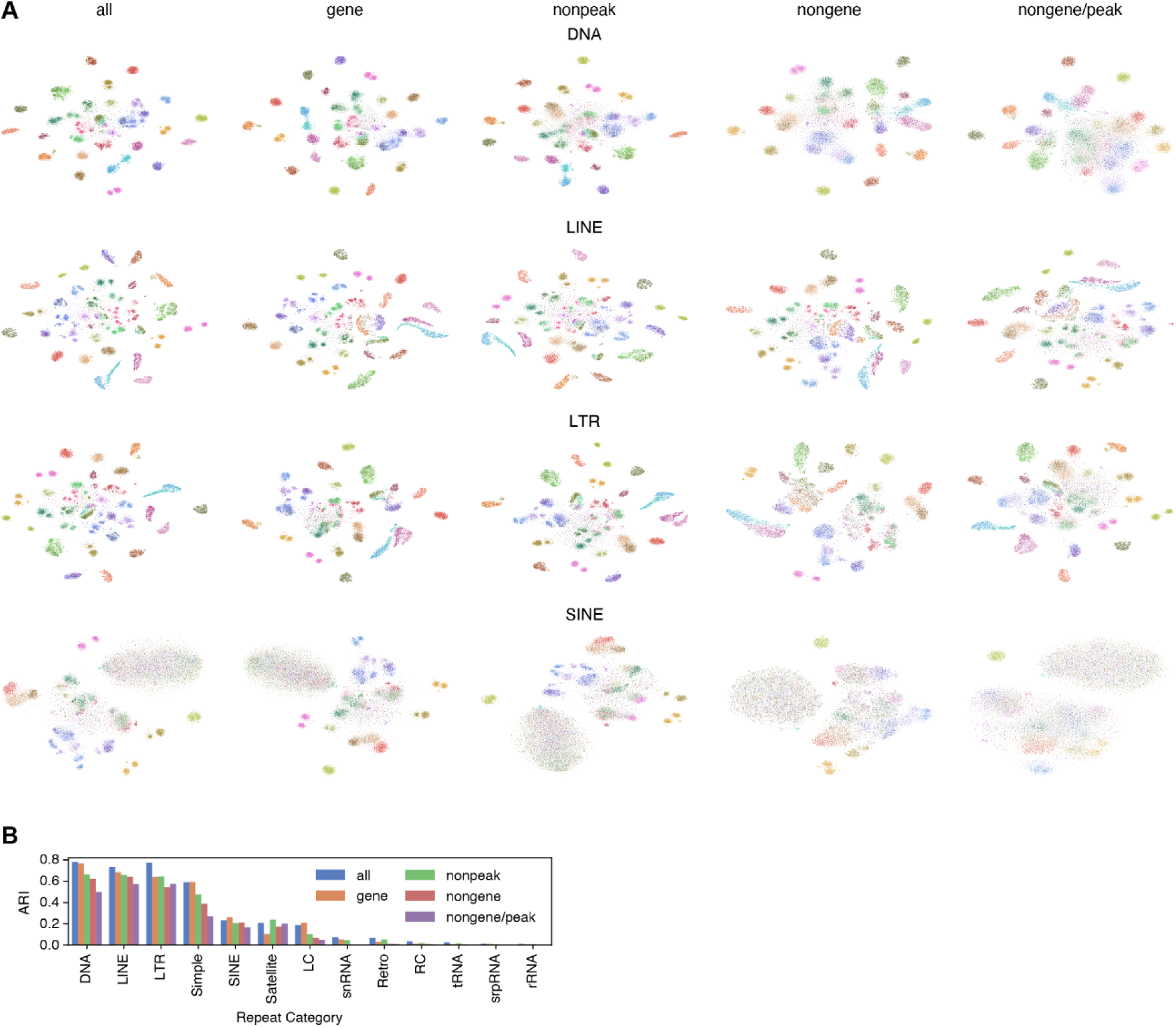
Cell clustering with mCG at repeat regions. (**A**) t-SNE of cells (n=26,423) using mCG of different repeat categories (row; LINE/SINE - long/short interspersed nuclear elements) and region exclusion criterions (column; gene - gene body ± 2k; nonpeak - excluding ATAC peaks of all major types; nongene - excluding all gene bodies ± 2k; nongene/peak - excluding both gene bodies ± 2k and ATAC peaks) colored by major types. (**B**) Barplots of ARI between major types and cell clustering using different repeat categories (x-axis) and region exclusion criterions (color).

**Fig. S14.**
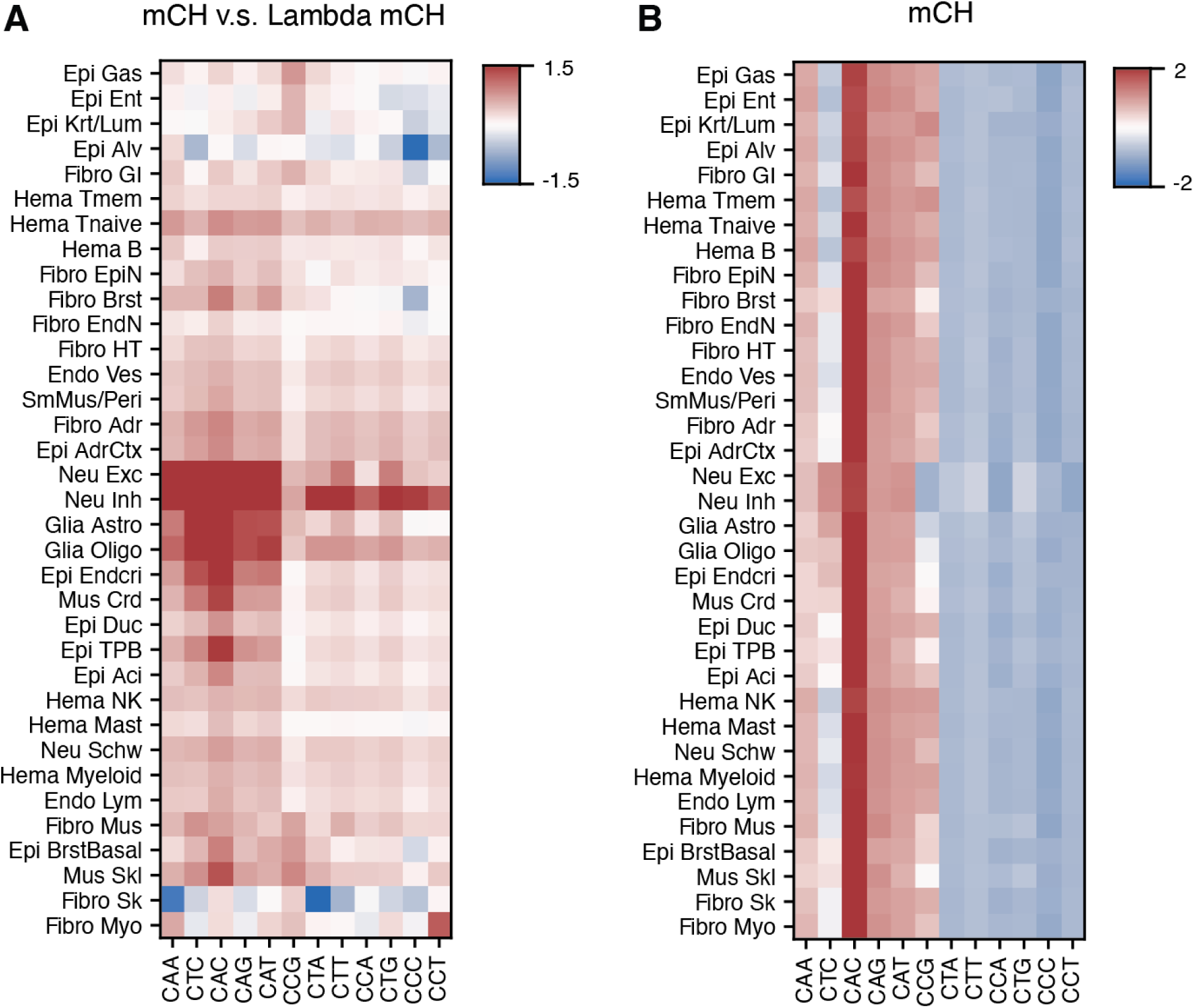
Non-CpG methylation compared with lambda phage controls. (**A**) Natural log fold change of genomic mCH versus control lambda phage mCH by base contexts across major types. (**B**) mCH by base context across major types. Values are row-wise Z-scored.

**Fig. S15.**
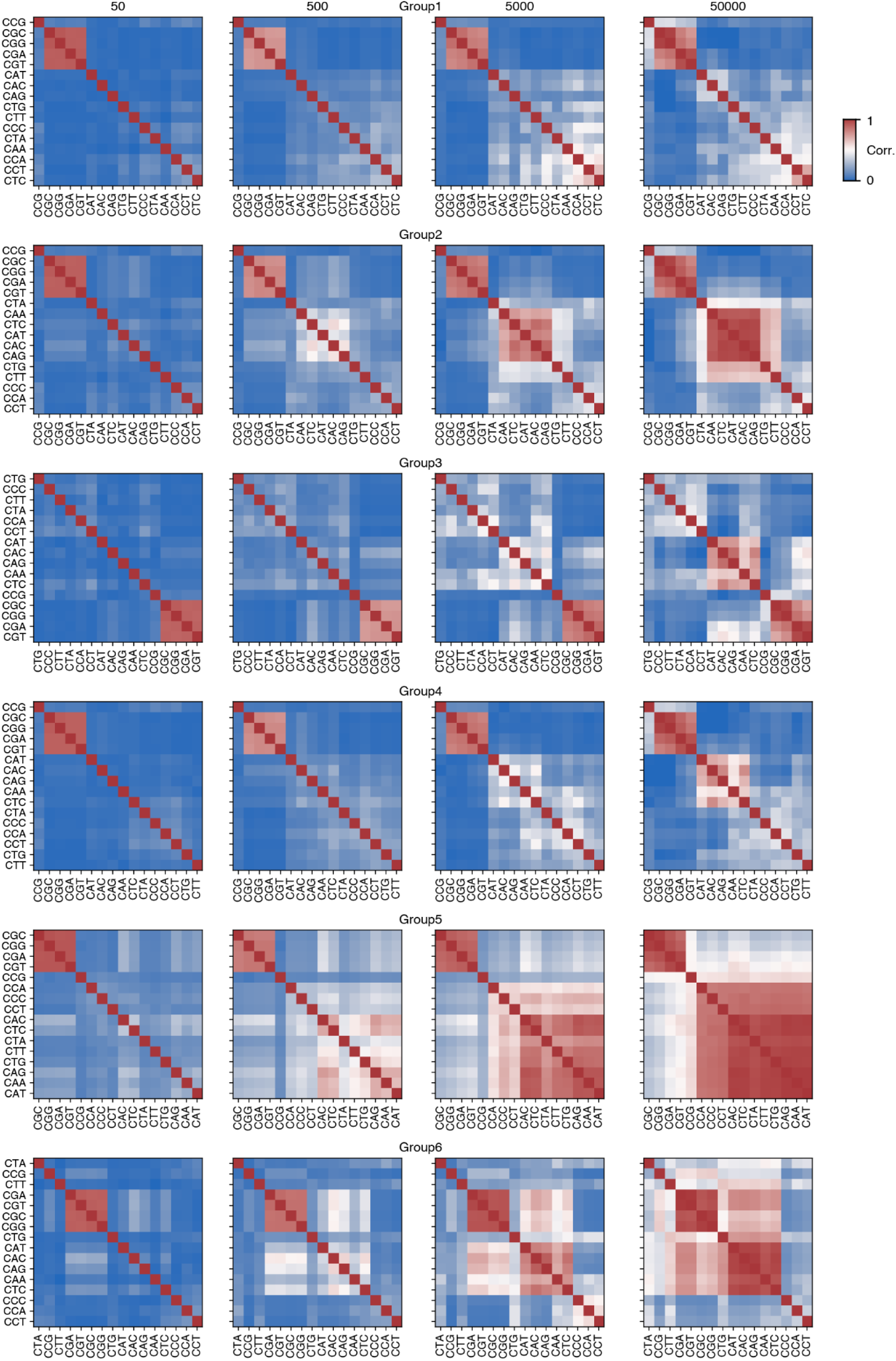
Correlations between methylation levels by trinucleotide contest. Correlations between 16 different trinucleotide contexts across genomic bins of different sizes (columns) for each of the six different major type groups in Fig. 3C (row).

**Fig. S16.**
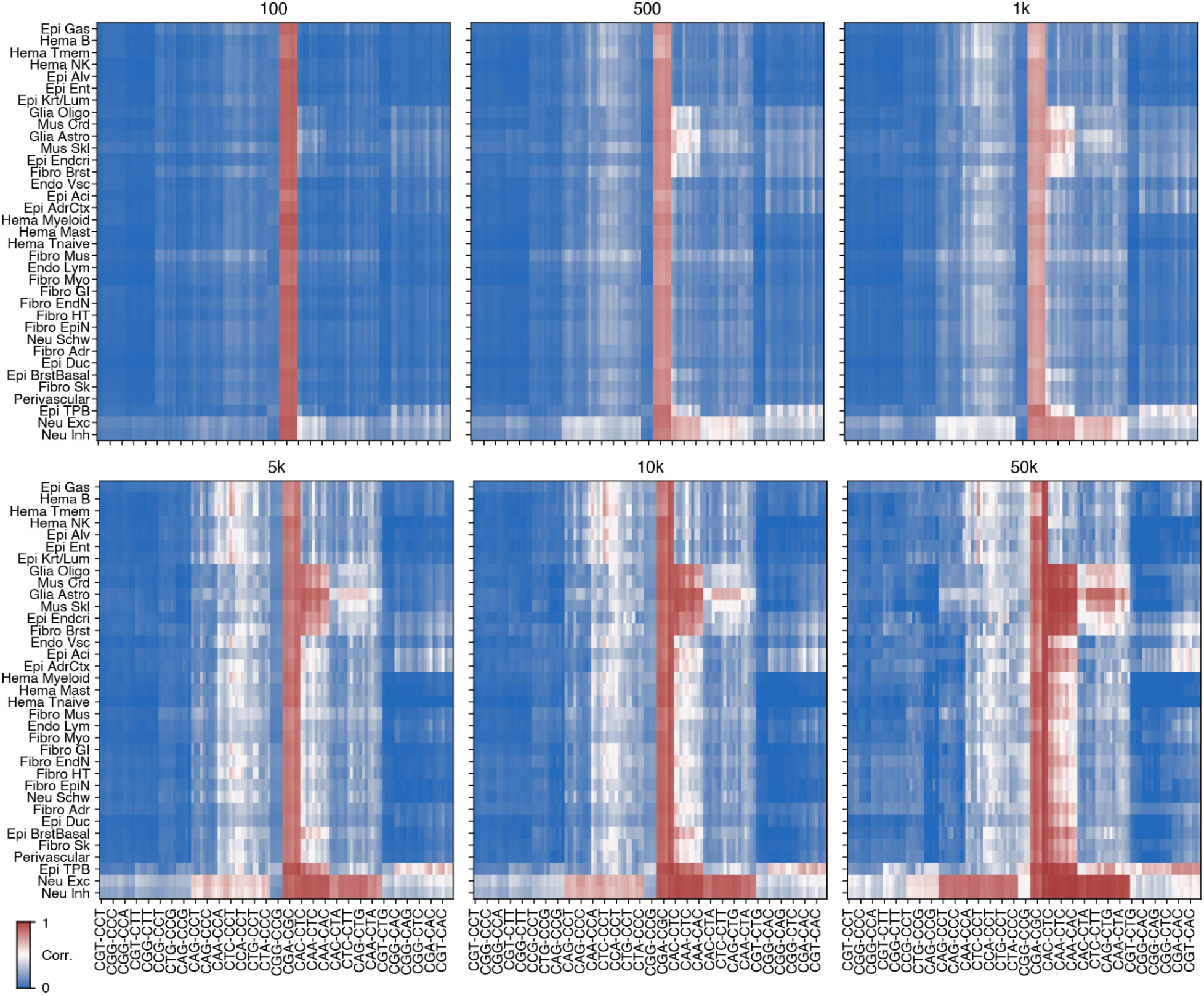
Clustering of trinucleotide correlations across lineages. Correlation of mC between pairwise trinucleotide contexts across genomic bins of different sizes (panel titles) in each cell type. Rows and columns are the same order as in Fig. 3C.

**Fig. S17.**
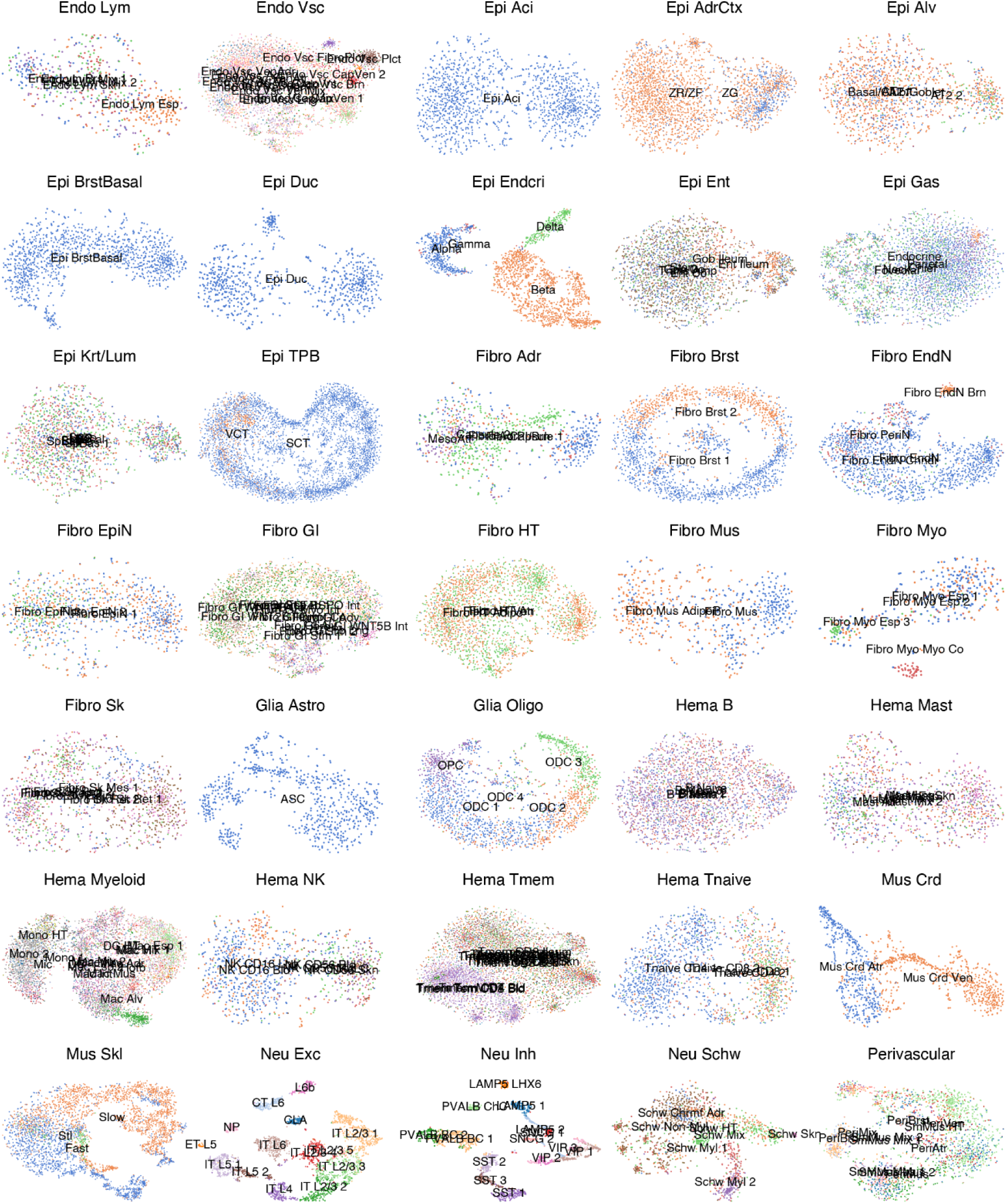
mCH subtype clustering. T-SNE of cells in each major cell type using 100kb bins mCH colored by subtypes.

**Fig. S18.**
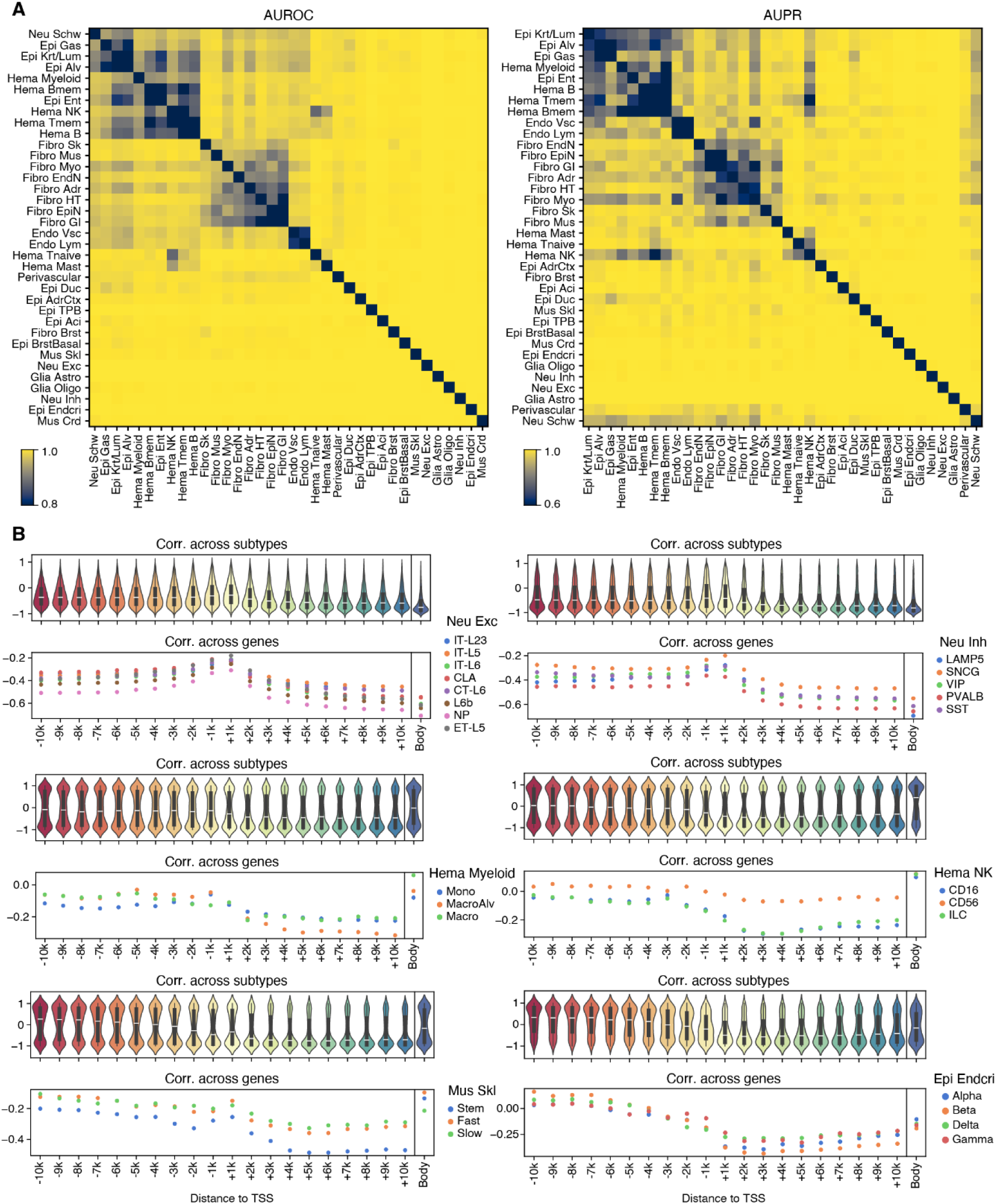
Correlation between gene body mCH and gene expression. (**A**) Area under the receiver operating characteristic (AUROC) and the area under the precision recall curve (AUPR) for logistic regression models to classify each pair of major types using 100kb mCH. (**B**) Correlations between mCH and expression of differentially expressed genes (DEGs) in each major type (legend title) across subtypes for all DEGs (top) or across all DEGs for each cell subtype (bottom). The correlations are calculated using mCH of regions from TSS to different distances on each side of TSS (x-axis) or across the entire gene body (right).

**Fig. S19.**
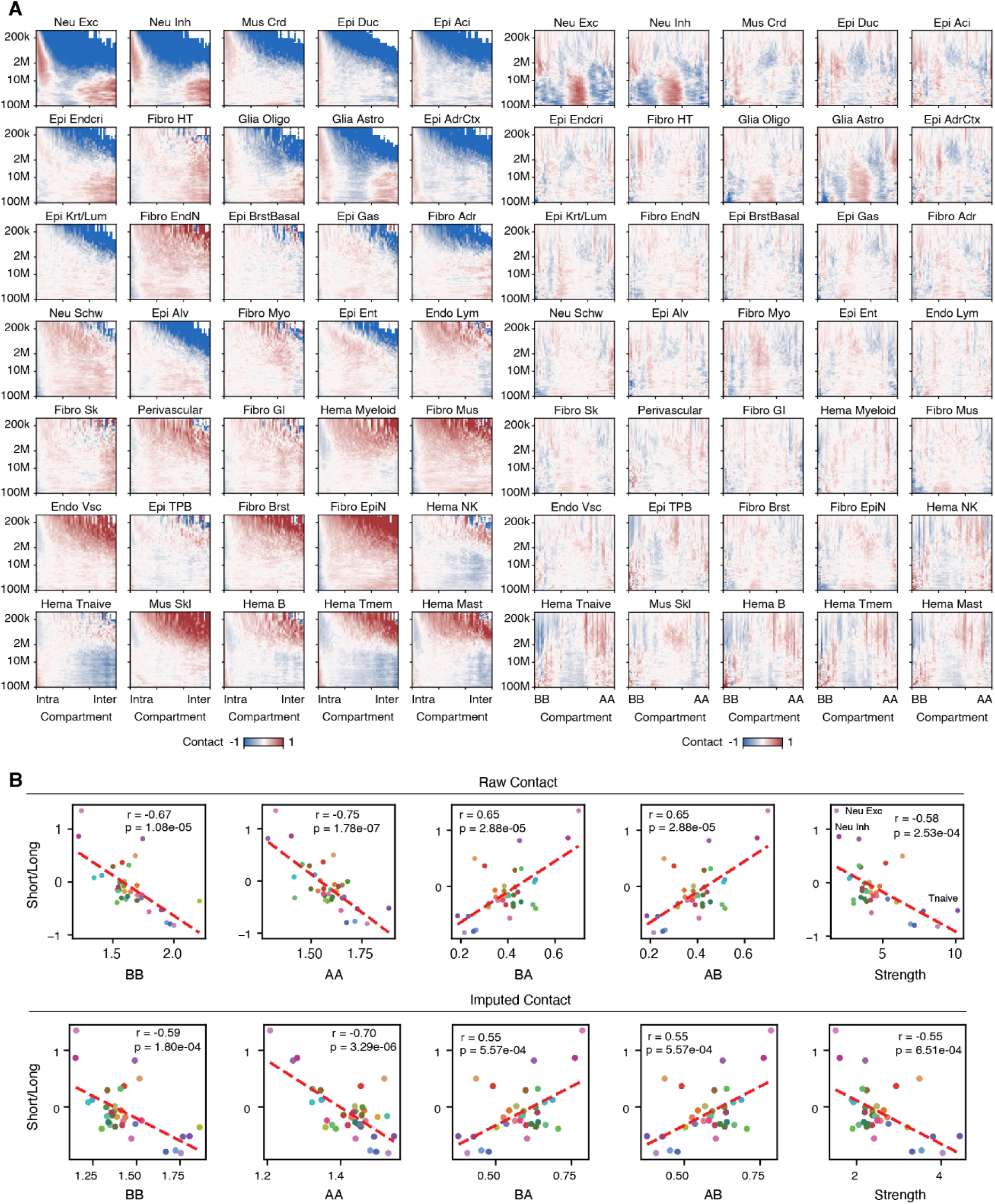
Compartment and domain dominant major cell types. (**A**) For each major type, proportion of contacts at a certain distance (y-axis) stratified by the differences (x-axis; left) or sums (x-axis; right) of compartment scores at the two interacting regions. Values are row-wise Z-scored. (**B**) Comparison of the raw (top) or imputed (bottom) frequency of chromatin compartment interaction types (AA, BB, AB, or BA) or the compartment strength (right most plot) versus the log2 short/long contact ratio across major cell types.

**Fig. S20.**
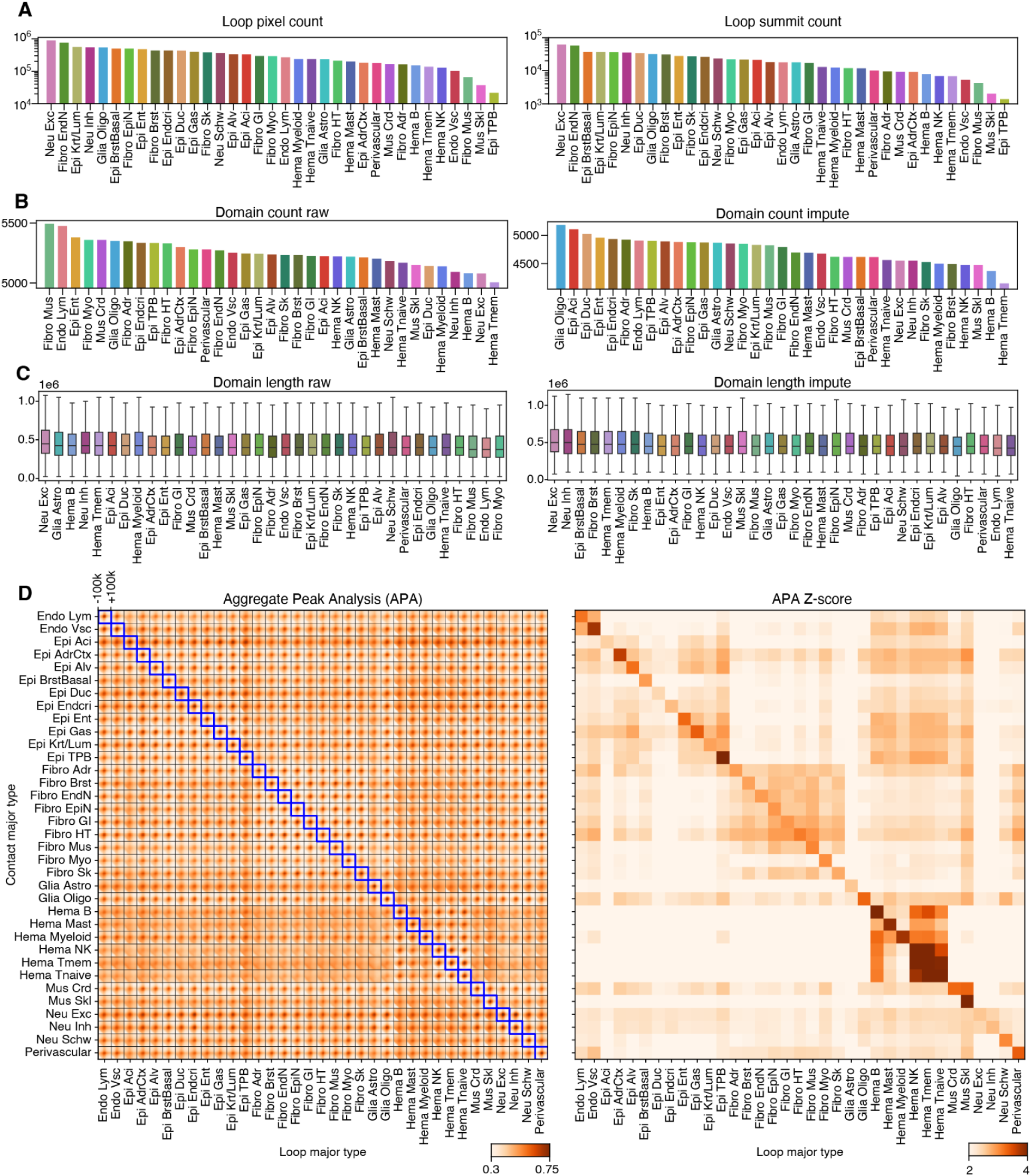
Chromatin loops and domains across lineages. (**A**) Number of unique pixels that are called as loops across major cell types (left) or number of unique loop summits across each lineage after merging adjacent pixels (right). (**B**) Number of domains in each lineage based on raw (left) or imputed (right) contacts in each lineage. (**C**) Distribution of domain lengths from diverse lineages using raw (left) or imputed (right) contacts. (**D**) Aggregate peak analysis (APA) of loops called in each major type (x-axis) using the imputed contacts of each major type (y-axis). The right plot is derived from the left plot, but shows the APA Z-score (Peak relative to lower left 5×5 quadrant).

**Fig. S21.**
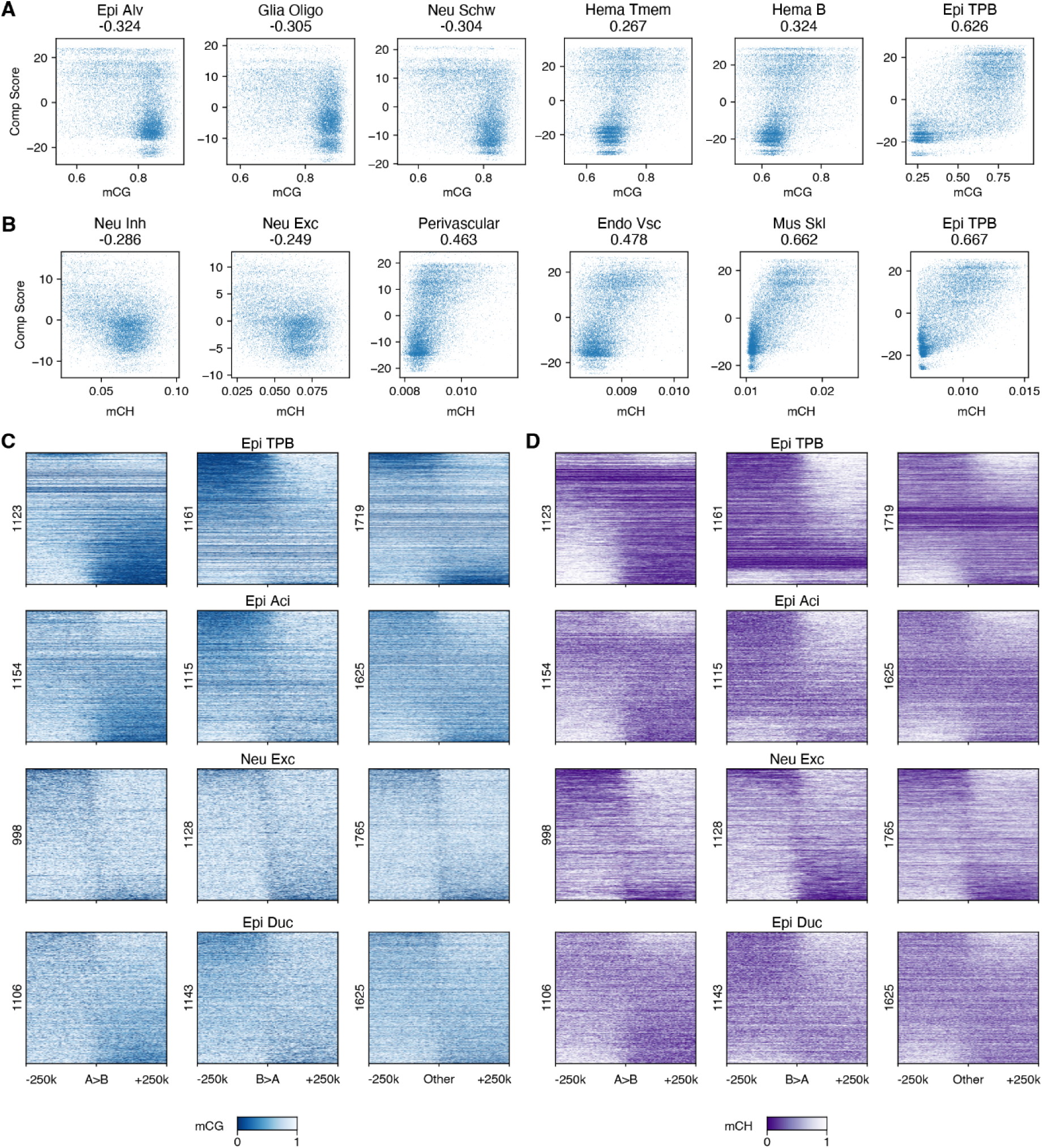
Comparison of DNA methylation, chromatin compartments, and domain boundaries. (**A,B**) Comparison of mCG (**A)** and mCH (**B**) with chromatin compartment scores across 100kb bins. Major types with highest correlation coefficients (both positive and negative) are shown. (**C,D**) mCG (C) and mCH (D) in flanking regions of domain boundaries that separate A and B compartments (A>B or B>A) or that do not (Other). The rows are sorted by the differences of DNA methylation upstream versus downstream of the boundary.

**Fig. S22.**
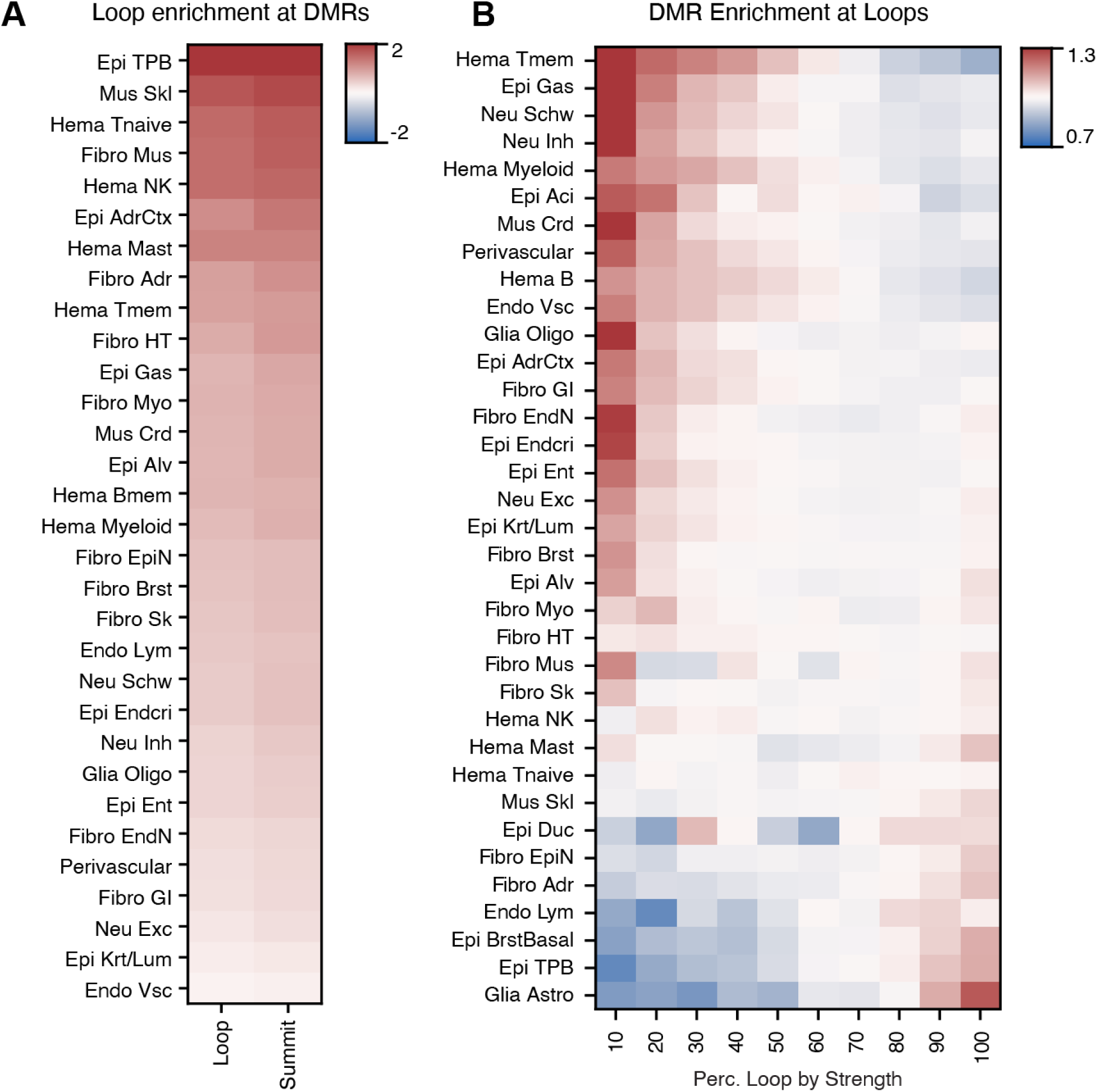
Enrichment of DMRs at loops and differential loops. (**A**) log2 odds ratio between proportion of loop (summit) anchors overlapped with DMRs versus proportion of non blacklist-flanking 10kb bins overlapped with DMRs. Colors are centered at 0, so red means enriched. (**B**) Odds ratio between proportion of DMRs overlapped with differential loop anchors versus proportion of loop anchors in differential loops using different quantile thresholds of Qanova and Tanova for differential loops.

**Fig. S23.**
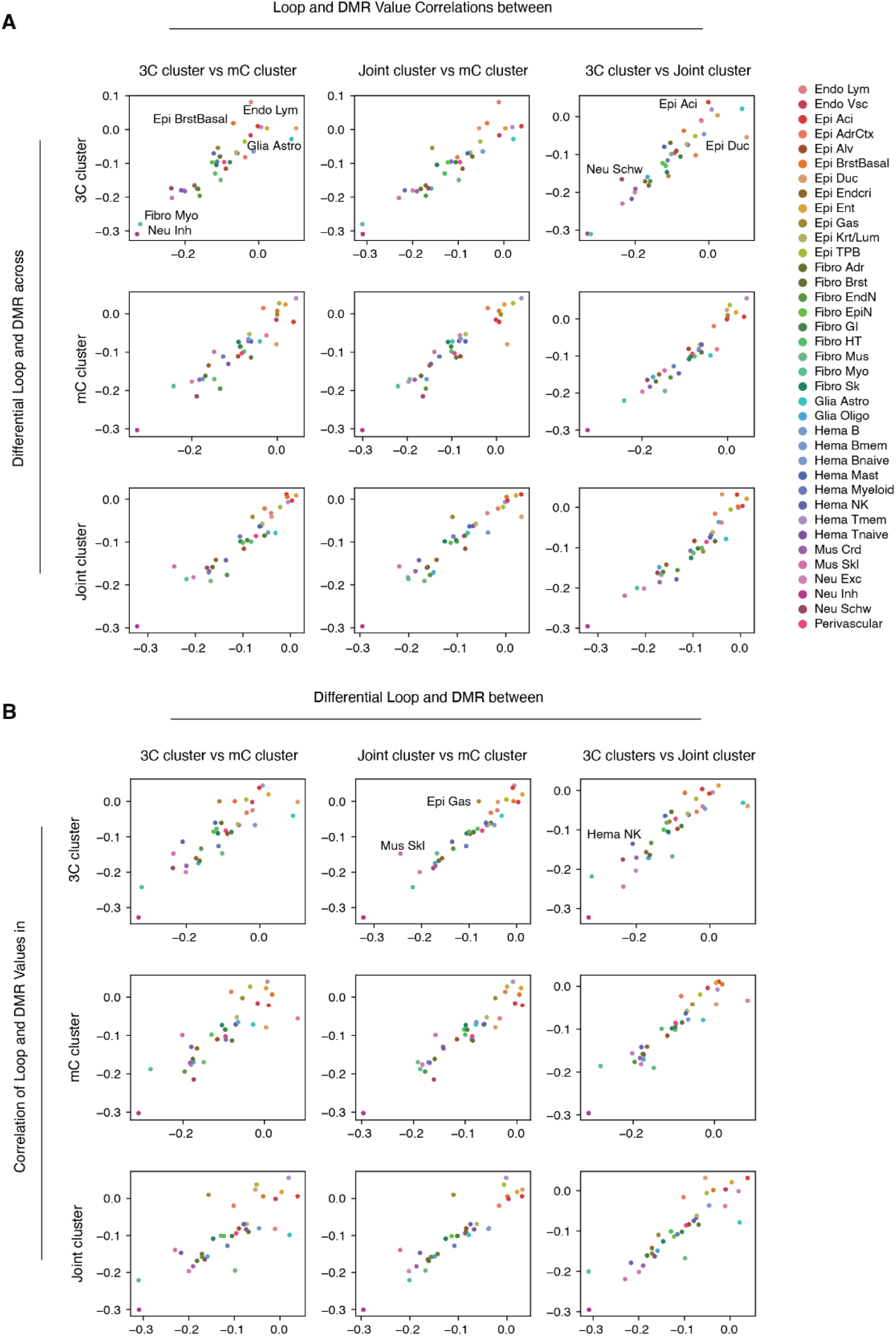
Comparison of loops and DMRs depending on clustering methods. (**A**) Comparison of the correlations between mCG at DMRs and strength of differential loops by the different modalities used for clustering. Within each major cell type, differential loops and DMRs are called based on the clustering using the data type shown in the row label and the coordinates of the differential loops and DMRs are overlapped. Using these coordinates, the correlation between DMRs and loop strength is calculated after independently clustering cells within each major type using clusters based on the column data type labels. The first part of the column label is displayed on the X-axis, and the second part of the column label is on the Y-axis. (**B**) Similar to A, but the DMRs and differential loops are first called using the clustering of data types from the column titles, and the correlations are computed after independently clustering the cells within each major type based on the data modality from the row labels.

**Fig. S24.**
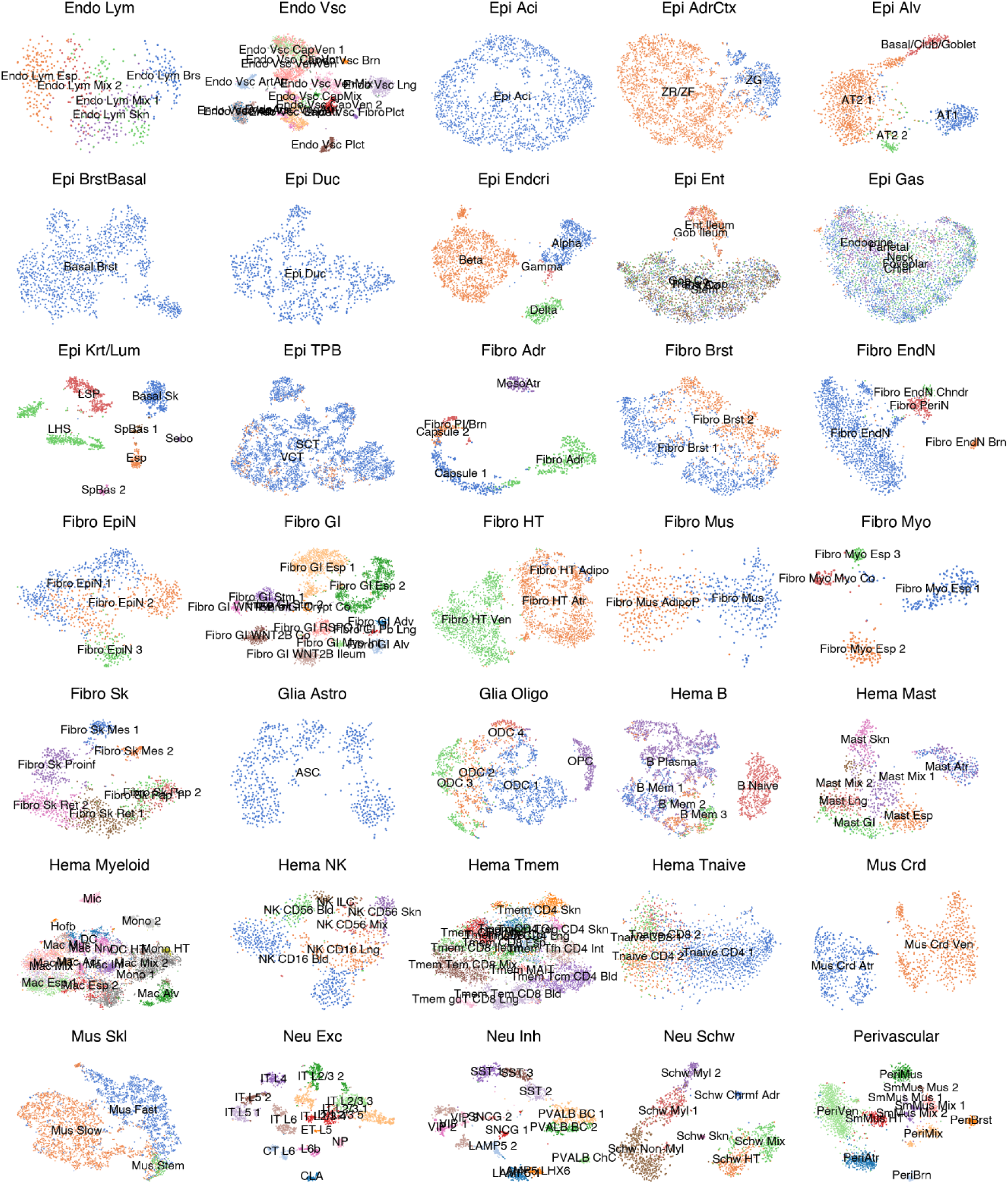
Subtype embeddings using DNA methylation. T-SNE of cells from each major type using mCG colored by subtypes.

**Fig. S25.**
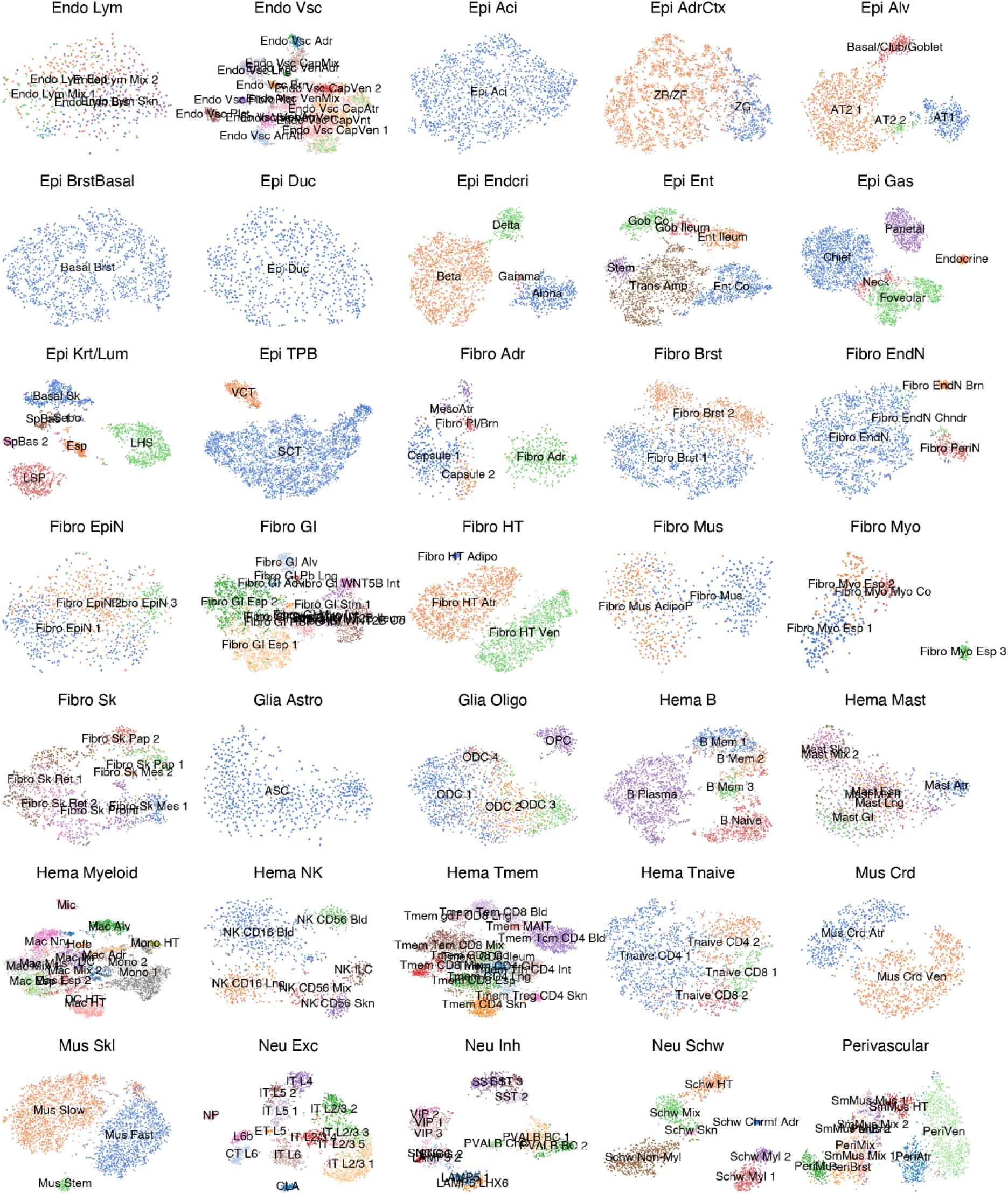
Subtype embeddings using chromatin contacts. T-SNE of cells from each major type using chromatin contacts colored by subtypes.

**Fig. S26.**
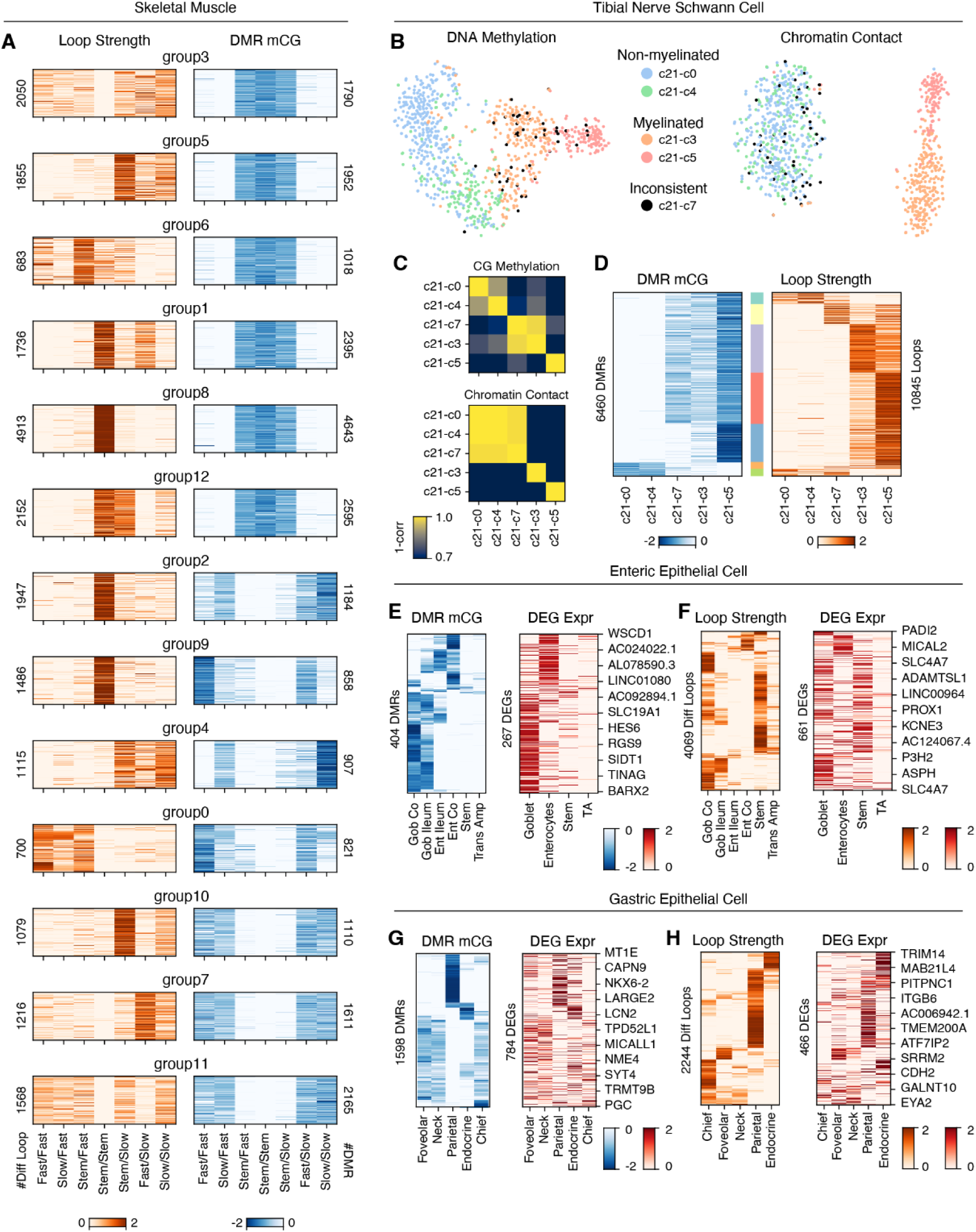
Differences in cell states from chromatin contacts and DNA methylation. (**A**) The same values as in Fig. 6F are shown, but different groups of loop-DMR pairs are separated for better visualization. The number of unique differential loops or unique DMRs in each group is shown as y-axis labels. (**B**) t-SNE of tibial nerve Schwann cells using DNA methylation (left) or chromatin contacts (right) colored by clusters. Black cells are those that show inconsistent associations with myelinated or non-myelinated cells depending on data types. (**C**) Correlation matrix between five Schwann cell clusters based on mCG (top) or chromatin contacts (bottom). (**D**) mCG of DMRs between cell clusters (left) or interaction strength of differential loops with either anchor overlapped with DMR (right). Values are Z-score normalized within each row. Colorbar (middle) shows *k*-means clusters of loop-DMR pairs. (**E-H**) mCG of DMRs (left; E,G) or interaction strength of differential loops (left; F,H) between subtypes of gastric (Epi-Gas; E,F) or enteric (Epi-Ent; G,H) epithelial cells and expression of DEGs (right) whose TSSs are within 2kb of DMRs or either anchor of the differential loops (right). Values are Z-score normalized within each row.

**Table S1. Samples used in the study.** The donor source of each tissue and age of donors are listed.

**Table S2. Number of dimensions used for clustering cells within each major type.**

**Table S3. Single-cell RNA-seq datasets used in the study for cell type annotation.**

**Table S4. Map between cell types annotated in Zhang et al. 2021 and major types in this study.**

